# Leveraging death of drug-sensitive cancer cells to promote immune-mediated bystander killing of subclones of drug-resistant tumour cells

**DOI:** 10.1101/2025.10.27.684825

**Authors:** Mona Tomaschko, KangBo Ng, Christopher Moore, Claire E. Pillsbury, Sareena Rana, James Campbell, Saptaparna Mukherjee, Ania Mikolajczak, Panayiotis Anastasiou, Andrea de Castro, Alicia Alonso de la Vega, Sophie de Carné Trécesson, Nathan W. Goehring, Miriam Molina-Arcas, Julian Downward

## Abstract

The response of lung cancer patients to drugs targeting the G12C mutant form of KRAS is limited by the development of resistance through multiple mechanisms. In order to achieve lasting benefit with these therapies, effective strategies for tackling the evolution of drug resistance are required. We have developed a preclinical model system to mimic the development of resistance to KRAS G12Ci inhibitors (G12Ci) such as adagrasib and RMC-4998. Treatment of tumours containing a minor subpopulation of resistant cancer cells with G12Ci leads to their rapid outgrowth to replace the drug-sensitive cells within a few weeks. However, when combined with therapies that, at least in part, target the immune response, such as SHP2 inhibitors or PD-1 blockade, drug-resistant cells can be eliminated, even by drug combinations that do not impact their growth in the absence of drug-sensitive cells. This bystander killing of drug-resistant cells when drug-sensitive cells are targeted is dependent on an intact adaptive immune system. Mechanistically, these combination therapies lead to profound remodelling of the tumour immune microenvironment, with influx of T cells recognising a tumour associated antigen shared between drug-resistant and drug-sensitive cancer cells. Promotion of immune-mediated bystander killing of drug-resistant cells may provide a paradigm for tackling the problem of drug resistance in cancer more broadly.

## INTRODUCTION

KRAS is the most frequently mutated oncogene in human cancer, with an incidence rate of about 90% in pancreatic adenocarcinoma, 50% in colorectal cancer, and 30% in lung adenocarcinoma^1^. KRAS proteins cycle between a GDP-bound (OFF) state and a GTP-bound (ON) state, the latter being able to interact with and activate downstream effector proteins such as RAF and PI3K^2^. In mutant KRAS proteins, GTP hydrolysis is impaired, leading not only to promotion of aberrant growth and survival signalling, but also evasion of antitumour immunity^3,4^, including suppression of IFN-γ responses^5,6^, aberrant expression of immune checkpoint molecules^7^ and secretion of immunosuppressive molecules by cancer cells^8,9^. This results in recruitment of cells that exert immune regulatory functions, dysfunctional immune phenotypes or promote tumour progression through wound-healing-like programs.

Recent advances in the development of inhibitors that directly target KRAS have opened new treatment opportunities for eligible cancer patients^10,11^. The first clinical compounds developed that were capable of inhibiting KRAS were the KRAS G12C mutant selective inhibitors sotorasib and adagrasib, which irreversibly lock KRAS G12C into its GDP-bound (OFF) state^12,13^. Numerous preclinical studies with these drugs have demonstrated tumour control, durable responses and also reversal of immune suppression^4^. KRAS inhibition restored antigen presentation and IFN-γ responses while reducing expression of immune checkpoint molecules such as PD-L1 and CD47^6,12,14,15^. In the tumour microenvironment (TME), antigen-presenting and CD8^+^ cytotoxic T cell counts were increased, while spatial analysis revealed a greater infiltration depth of tumour-infiltrating lymphocytes (TILs)^16^ and more tertiary lymphoid structures (TLSs)^17^. On a systemic level, tumour control by KRAS inhibitory drugs was superior in immune-competent, compared to immune-compromised, mice and complete responder mice presented improved anti-tumour-directed immune memory^18^. Furthermore, inhibition of oncogenic KRAS signalling was found to sensitise responses towards immune checkpoint inhibitors (ICB)^6,14^.

Phase 2 clinical trials for adagrasib and sotorasib in KRAS G12C mutant lung adenocarcinoma showed promising results, with most patients responding initially, despite prior treatments, leading to accelerated FDA approval^19,20^. However, most patients also developed resistance within months and a subsequent phase 3 trial for sotorasib showed no overall survival advantage over docetaxel^21^. Following on from the development of KRAS G12C(OFF) inhibitors, active-state RAS(ON) G12C-selective inhibitors have also entered clinical trials^22^. The most advanced of these are molecular glues that bind a ubiquitously expressed chaperone, Cyclophilin A, forming a binary complex that selectively and covalently binds RAS G12C-GTP, preventing interaction with downstream targets. Early clinical data show initial responses in most patients, including those who previously progressed on G12C(OFF) inhibitors^23^. Long-term data are not yet available, but resistance is also anticipated to be very likely to occur, although the time to progression will be key. In parallel, many drugs targeting other mutant forms of KRAS and also unmutated RAS proteins are in advanced clinical development^4^.

Resistance to KRAS inhibitors can develop through many different mechanisms, including acquisition of further mutations in oncogenic KRAS proteins or related signalling components^24,25^, genomic amplification or overexpression of pathway components, including KRAS itself^26,27^, or activation of alternative pathways, such as epithelial-to-mesenchymal transition (EMT) or YAP signalling^28,29^. These mechanisms likely emerge from subclonal events and are selected for during therapy, although adaptive resistance due to signalling pathway rewiring can also occur^30^.

To overcome the rapid development of resistance, combination with other therapies is an attractive option^31^. As KRAS inhibitors reverse immunosuppressive features of the TME, one promising approach is to combine with treatments that enhance anti-tumour immunity, such as anti-PD-1/PD-L1 immunotherapy^6,14^. When used as a single agent in lung cancer, immunotherapies lead to a subset of patients achieving durable responses, but the majority of patients display primary resistance to this treatment^32,33^. Inhibition of SHP2 (SHP2i) also presents an interesting therapeutic strategy, by vertically reinforcing inhibition of oncogenic KRAS signalling in cancer cells^27^, while simultaneously also inducing beneficial changes in other TME cell populations, such as reducing cytotoxic T cell exhaustion, shifting macrophages to a less immunosuppressive polarised state, and normalising tumour vasculature^34–36^. While first-generation KRAS inhibitors were limited by off-target toxicities that hindered their use in combination with these immunotherapies, newer generations with improved specificity may overcome these challenges and offer a therapeutic window for effective combination strategies^37^.

Since rapid development of drug resistance is proving to be a key limitation for KRAS inhibitory drugs, we have set out here to explore novel methods for counteracting this phenomenon. As KRAS signalling promotes an immunosuppressive TME and KRAS inhibitors reverse this, at least in the short term, we reasoned that it might be possible that conditions exist whereby KRAS inhibitor induced killing of tumour cells could help prime an adaptive immune response against the tumour itself. Since most drug-resistant subpopulations of tumour cells would be expected to be very closely related to the drug-sensitive parental cells and share common antigens, any such immune priming could potentially act to promote destruction of the drug-resistant cells as well. Here, we set out to develop a mouse lung cancer model that mimics tumours containing a KRAS G12C inhibitor (G12Ci)-resistant subpopulation using isogenic KPAR1.3 cells^18^ harbouring either a KRAS G12C (G12Ci-responsive) or G12D (G12Ci-resistant) mutation co-engrafted in immune-competent mice. While KRAS G12Ci as monotherapy conveyed a proliferative advantage to drug-resistant tumour subpopulations, combination therapies with a SHP2i or anti-PD-1 resulted in immune rejection of drug-resistant tumour subpopulations, complete responses, and enhanced immune memory, even though these combinations did not impact the drug-resistant cells in isolation. Mechanistic studies revealed an increase in inflammatory-associated signatures in KRAS G12Ci-resistant cells and a profound, therapy-mediated remodelling in both innate and adaptive immune compartments in the TME, as well as an increase in memory T cells in tdLNs.

This work establishes the paradigm that it is possible to induce immune-mediated bystander killing of drug-resistant subpopulations by appropriate combinations of oncogene targeted inhibitors and immune targeting agents and provides a platform to explore this novel approach to countering drug resistance in cancer therapy, including gaining further mechanistic insights and optimal strategies for potentiating this effect.

## RESULTS

### A tumour model to mimic development of KRAS G12C inhibitor resistance as a subclonal event in solid tumours

To establish a model system that would allow us to study the onset of drug resistance in KRAS mutant lung cancer, we made use of KPAR1.3, a transplantable lung cancer cell line with an immune “hot” TME that we developed previously^18^, to mimic the presence of minor subpopulations of drug-resistant cancer cells within a predominantly drug-sensitive tumour. While originally from a KRAS G12D mutant genetically engineered mouse model and thus resistant to KRAS G12Ci, we also derived a KRAS G12C mutant version by prime editing, which is sensitive to KRAS G12Ci^18^. To be able to trace isogenic, co-engrafted KPAR1.3 KRAS G12C and G12D mutant cancer cells across both *in vitro* and *in vivo* applications, we first transduced the isogenic cell lines with reporter constructs. To avoid artefactual immune responses directed against reporter construct encoded proteins, we employed the glowing-head (GH) mouse model as recipient, which expresses a Luciferase-enhanced Green Fluorescent Protein (Luc-eGFP) fusion construct in the anterior part of the pituitary gland, thereby rendering the mice immune-tolerant towards these reporters^38^. The KRAS G12D mutant cell line was transduced with a Luc-eGFP construct identical to that used in the GH mouse model. In addition to GFP, the luciferase reporter further enables tracking of the presence of the KRAS G12Ci-resistant subpopulation over time in *vivo,* via bioluminescence scanning (IVIS Spectrum system). To obtain a second colour fluorescent protein for the KRAS G12C mutant cell line, while minimising the chance of aberrant reporter-directed immune responses, amino acid sequence homology across fluorescent proteins was compared and AlphaFold utilised for structural predictions. This revealed that enhanced Blue Fluorescent Protein (eBFP) and eGFP were structurally nearly identical, only differing in two residues, Y67H and Y146F, resulting in a sequence homology of 99.2 % (Figure S1a). Moreover, employing the ‘Immune Epitope Database’ (IEDB) MHC-l complex affinity modelling algorithm predicted that all eBFP-derived peptides containing the two differing amino acids would have very weak MHC-l loading properties (Table S1), so it is unlikely that eBFP would be recognised as foreign in mice with immune tolerance to eGFP.

The transduced cell lines were then subcloned and characterised *in vitro* to check for retention of the same growth rates as the parental line (Figure S1b). Luciferase reporter activity was demonstrated for the KRAS G12D clones (Figure S1c). As expected, only the KRAS G12C clones were sensitive to the KRAS G12C(OFF) inhibitor adagrasib (Figure 1a), while similar responses to MEK inhibitor trametinib were observed in all clones (Figure S1d). To assess how population dynamics between selected isogenic subclones would evolve during continuous adagrasib treatment in *vitro*, mixed KRAS G12C and KRAS G12D co-cultures were set up at ratios of 90:10, 95:5, and 99:1 and population fractions were assessed twice weekly via flow cytometry while on treatment (Figure S1e). In contrast to DMSO-treated control co-cultures, the fraction of KPAR1.3 KRAS G12C cells in adagrasib-treated samples fell near zero within 2-4 weeks, even in the 99:1 co-cultures where the starting fraction of G12D cells was only 1%.

**Figure 1.**
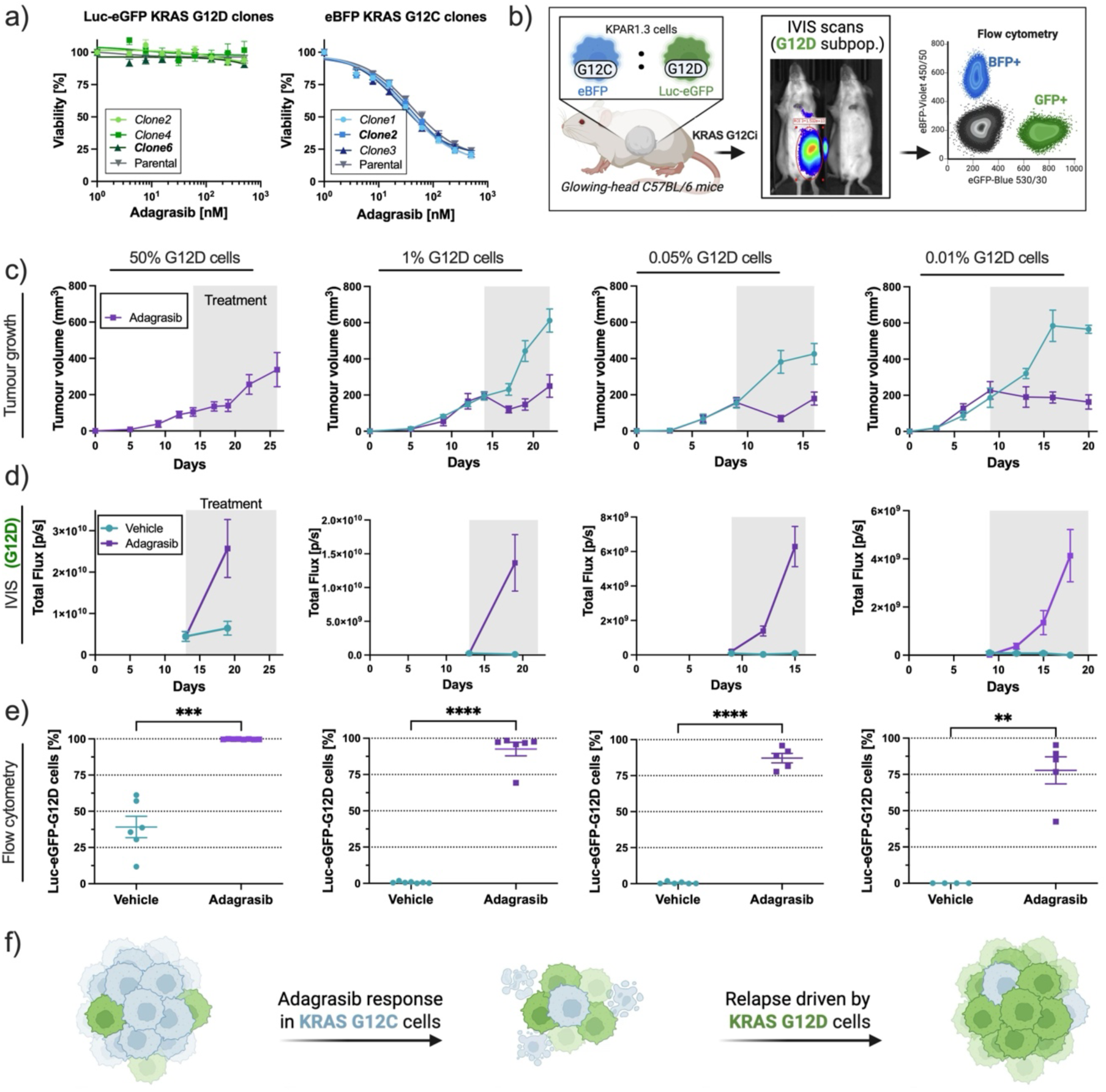
KRAS G12Ci as monotherapy promotes the outgrowth of drug-resistant subpopulations. a) Viability assays of selected KPAR1.3 subclones with the genotypes Luc-eGFP KRAS G12D (left) or eBFP KRAS G12C (right) compared with the parental counterparts. Cells were treated with adagrasib at indicated concentrations for 72 hours. Mean + SEM of 3 biological replicates (n) with 3 technical replicates (tr) each. b) Illustration of the experimental setup for panels (c) to (e). Subcutaneous tumours were engrafted with different ratios of eBFP KRAS G12C and Luc-eGFP KRAS G12D KPAR1.3 cells, indicated on top of the graphs in panel (c). Mice treated with the KRAS G12C inhibitor adagrasib were dosed daily with 50 mg/kg. Grey area on respective plots indicates duration of treatment. n= 6-8 mice per group. c) Grouped tumour growth over time for indicated ratios of KRAS G12C: KRAS G12D cancer cells. d) Bioluminescence (IVIS) scans over time, detecting luminescent signal emitted by luciferase-expressing KRAS G12D tumour cells. e) Flow cytometric analysis of Luc-eGFP KRAS G12D cancer cells of all live cancer cells. All graphs display mean ± SEM; statistics indicate pair-end student t-test with Welch’s correction for varying standard deviation between treatment groups. f) Schematic of subpopulation dynamics.

To confirm that the selected subclones also proliferated at equal rates in *vivo*, the reporter-traced KRAS G12C/G12D mutant cell lines were co-engrafted subcutaneously at a 50:50 ratio in GH-mice and cancer cell fractions from resulting untreated tumours were determined after 14 days by flow cytometry (Figure S2a and S2b). Additionally, tumours were sampled to visualise the spatial distribution of the isogenic cell lines within the tumours via immunohistochemical staining, which revealed an overall interspersed growth pattern between the two populations (Figure S2c and S2d). In these stainings, the GFP antibodies recognise both cancer cell populations as they cross-react with the closely related BFP protein, while the luciferase antibodies recognise only the KRAS G12D cells.

To model the early onset of treatment resistance in a preclinical setting, we next focused on establishing a murine model system harbouring KPAR1.3 KRAS G12C-derived tumours, co-engrafted with a subpopulation of isogenic KRAS G12Ci-resistant (KRAS G12D mutant) tumour cells. It was previously shown that KRAS G12C inhibition in responsive tumour cells can at least partially reverse immune suppression in the tumour microenvironment, and can result in complete tumour regressions in some mice^6^. We therefore investigated whether employing KRAS G12C inhibitors as a monotherapy may promote anti-tumour-directed immunity that would target both KRAS G12Ci-sensitive and -resistant cancer cell clones and prevent the outgrowth of drug-resistant subpopulations through immune-recognition of shared tumour associated antigens or tumour specific antigens. To test this, *in vivo* experiments with four different engraftment ratios of eBFP KRAS G12C and Luc-eGFP KRAS G12D KPAR1.3 cells in a subcutaneous setting were performed (Figure 1b). Responses in vehicle or adagrasib treated mice were assessed via measurement of tumour growth (Figure 1c) and the abundance of KRAS G12D mutant tumour cells over time was measured via bioluminescence scans (Figure 1d). Additionally, the fraction of KRAS G12D mutant cancer cells out of all live tumour cells at the end of treatment was determined by flow cytometry (Figure 1e). Even when engrafting very few G12Ci-resistant KRAS G12D mutant cancer cells (0.01% KRAS G12D cells, equivalent to 15 cells out of 150,000 engrafted cells per tumour), the resistant subpopulation gained a strong proliferative advantage in adagrasib treated tumours, with the majority of tumours being mostly comprised of KRAS G12D mutant cancer cells within two weeks of treatment. From this series of experiments, we conclude that treatment with adagrasib monotherapy promotes the outgrowth of KRAS G12Ci-resistant tumour subpopulations, and that any potential enhanced immune response against shared tumour antigens is not sufficient to generate complete responses, even when only very small numbers of drug-resistant cells are present (Figure 1f).

### Combination therapies enhance treatment responses and promote elimination of KRAS G12Ci-resistant subpopulations

We next wanted to assess whether synergy between targeted KRAS G12C inhibition and further lines of therapies, particularly those known to bolster anti-tumour immunity, may promote immune attack on KRAS G12Ci-resistant tumour subpopulations, with the death of KRAS G12Ci sensitive cells potentially priming an immune response that would target shared tumour antigens on both populations. First, we selected an immune checkpoint blockade antibody targeting PD-1, which primarily acts through preventing T cell exhaustion and enhancing anti-tumour directed T cell effector functions^32^. Secondly, a SHP2 inhibitor (SHP2i), RMC-4550, was chosen. SHP2 is a pleiotropic drug target: in cancer cells, SHP2i can reinforce vertical pathway inhibition, as SHP2 is involved in KRAS activation^27^, while simultaneously acting on immune cells in the TME by preventing T cell exhaustion, repolarising macrophages (MPs) toward an anti-tumour phenotype, and normalising tumour vasculature^34–36^.

Before testing these drug combinations in vivo, we assessed the cancer cell intrinsic sensitivity to SHP2 inhibition for both KRAS G12C and G12D mutant KPAR1.3 cells. It is known that different oncogenic amino acid substitutions in KRAS can convey differing properties on the mutant protein in terms of GTP cycling^39^. While KRAS G12C mutation results in loss of GAP binding, however, the protein-intrinsic GTPase activity remains largely intact. On the other hand, KRAS with a G12D mutation is affected by both impaired GAP binding and also loss of intrinsic GTPase activity, so KRAS G12D remains in the GTP-bound active state for longer compared to KRAS G12C. Thus, KRAS harbouring a G12C mutation may require more frequent upstream reactivation and therefore be more sensitive to SHP2i, compared to KRAS G12D. Indeed, in vitro viability assays revealed that SHP2i elicited a greater response in KRAS G12C mutant cancer cells compared to KRAS G12D mutant cells, especially at reduced serum concentrations (Figure S3a).

To look at these effects in vivo, immunocompetent mice were subcutaneously engrafted with either the KRAS G12C or G12D cell lines. In these studies, we used the active-state RAS(ON) G12C-selective inhibitor, RMC-4998, a tool compound representative of the investigational drug candidate elironrasib (henceforth referred to as KRAS G12Ci)^22^. Pure KRAS G12C-derived tumours were responsive to KRAS G12Ci treatment, with 3 out of 8 complete responders (CR) when treated for two weeks, followed by drug withdrawal (Figure S3b). Pure KRAS G12C-derived tumours also partially regressed when treated with SHP2i (Figure S3c), although tumours grew back after the treatment period and no complete responders were observed. Using anti-PD-1 treatment, pure KRAS G12C-derived tumours progressed on treatment, with only a modest treatment response compared to isotype control being observed (Figure S3d). On the other hand, pure KRAS G12D-derived tumours did not respond visibly to either SHP2i (Figure 2a) or anti-PD-1 treatment (Figure 2b), demonstrating that these tumours are resistant to all three lines of therapy used here, at least when administered as monotherapies.

**Figure 2.**
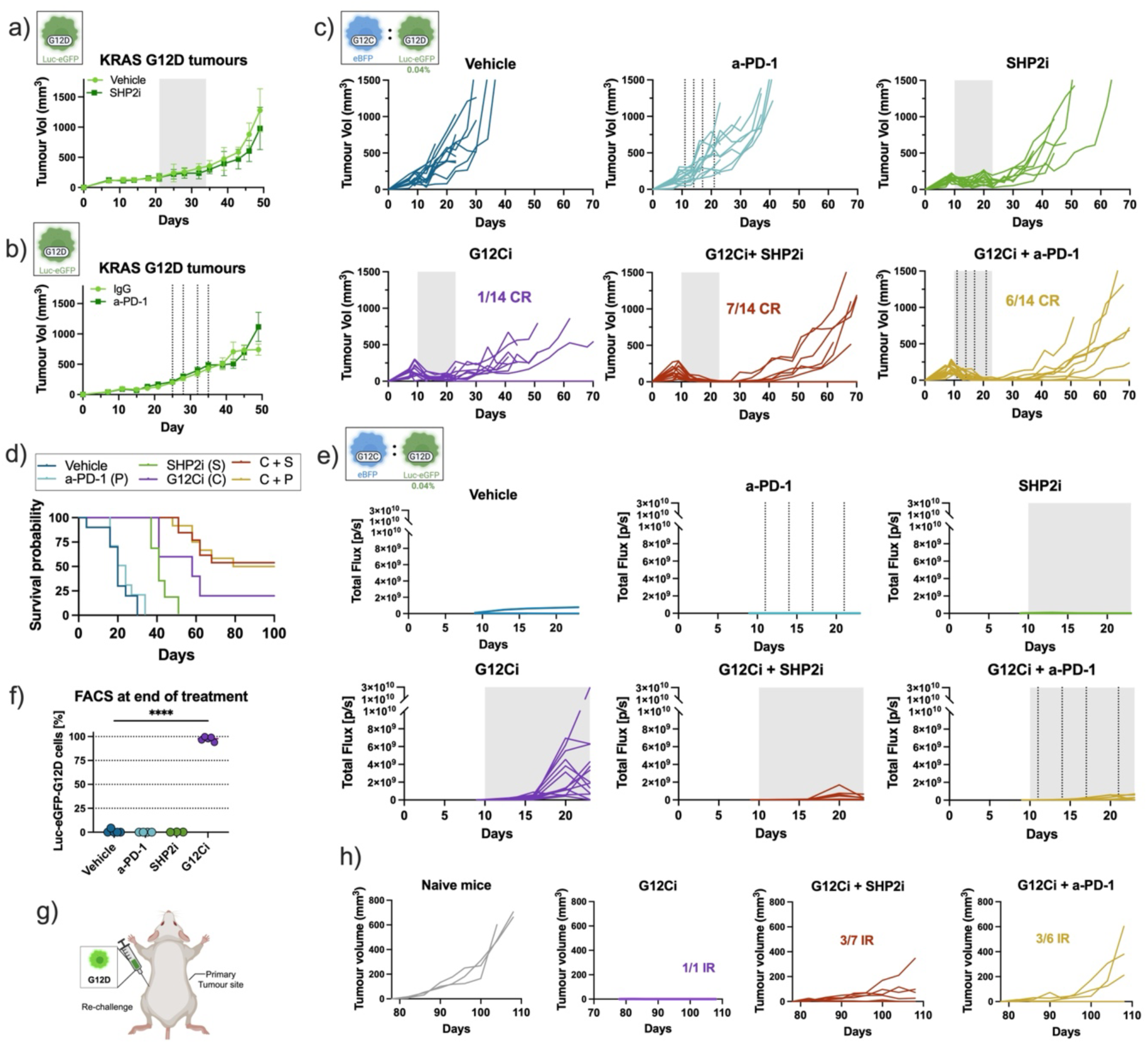
Combination therapies promote elimination of resistant subclones. (a-b) Tumour growth of subcutaneous tumours derived from Luc-eGFP KRAS G12D KPAR1.3 cells. Error bars indicate mean ± SEM. a) Mice were treated with either vehicle (n=7) or RMC-4550 (30 mg/kg; SHP2i; n=8). Treatment period indicated as grey area on plots. b) Mice were treated with either isotype control (10 mg/kg, IgG, n=6) or a-PD-1 (10 mg/kg; n=7). Doses indicated by vertical dotted lines. (c-f) Mixed subcutaneous tumours were engrafted with BFP KRAS G12C cells plus a fraction of 0.04% Luc-eGFP KRAS G12D cells. Mice were treated for two weeks with the RAS G12C(ON) inhibitor RMC-4998 (100 mg/kg; G12Ci) with or without the SHP2 inhibitor RMC-4550 (30 mg/kg; SHP2i) or a-PD-1 (10 mg/kg). RMC-4998 and RMC-4550 were dosed by oral gavage daily (grey area) and a-PD-1 was administered i.p. twice a week (vertical dotted lines). c) Tumour volume over time of individual tumours by treatment group. Mice remaining tumour free for at least 55 days after treatment withdrawn are indicated as complete responders (CR) in respective plots. d) Survival probability across different treatment groups. Tumours taken out for flow cytometry at endpoint were excluded. Cut-off value: 500 mm^3^. e) Bioluminescence scans, indicating the relative abundance of Luc-eGFP KRAS G12D KPAR1.3 cells over time, measured in total flux, photons per second [p/s]. f) Representative tumours were isolated from vehicle and single-therapy treatment groups and fraction of live Luc-eGFP KRAS G12D cells within all cancer cells was determined via flow cytometry. Each dot represents one tumour, mean ± SEM; statistics indicate one-way ANOVA test comparing each of the treatment groups to the vehicle, only significant comparisons are shown. g) Schematic of rechallenge experiment, performed on CR mice from panel (c). Mice were injected in the opposite flank with Luc-eGFP KRAS G12D cells. h) Tumour growth over time of re-challenged tumours in naïve mice or CRs. Mice which were able to immune reject (IR) the re-injected cancer cells out of all mice are indicated. Graph titles indicate the treatment that the primary tumour received.

Next, we assessed whether combinations of these treatments could elicit durable responses in mice engrafted with KRAS G12C mutant tumours containing a small fraction, 0.04%, of treatment-resistant KRAS G12D cells. In this mixed tumour setting, the responses in terms of tumour growth from mice treated with either vehicle, SHP2i, or anti-PD-1 as monotherapy (Figure 2c) was comparable to previous experiments of mice engrafted with KRAS G12C-derived tumours (Figure S3b-S3d). Mice receiving KRAS G12Ci as monotherapy showed tumour regression during the treatment interval, with one mouse achieving a CR. On the other hand, combining KRAS G12Ci with either SHP2i or anti-PD-1 led to complete and durable responses in approximately half of the mice, which further remained tumour-free off treatment for some seven weeks until the end of the experiment. These responses were achieved despite the presence of KRAS G12D mutant tumour subpopulations that did not visibly respond to any of the administered anti-cancer treatments when growing as pure tumours. Synergistic effects between KRAS G12Ci and SHP2i or anti-PD-1 were also evident when visualising survival probabilities across treatment groups (Figure 2d).

Bioluminescence scans revealed further mechanistic insights into the dynamics of the two tumour cell fractions over time (Figure 2e). The amount of KRAS G12D cells in the tumours remained very low on treatment of mice with either vehicle, SHP2i or anti-PD-1 as monotherapies. On the other hand, similar to the previous experiments with adagrasib (Figure 1d), bioluminescence scans again indicated that the KRAS G12D subpopulation increased substantially during RMC-4998 treatment as monotherapy (Figure 2e). However, significantly, bioluminescence scans of combination-treated mice indicated enhanced control of the KRAS G12Ci-resistant tumour subpopulation in addition to palpable tumour regression being observed (Figure 2e). Moreover, bioluminescence signal at the end of treatment correlated with the generation of complete responses (CRs), with most mice achieving CRs showing no clear outgrowth of the KRAS G12D subpopulation (Figure S3e), again despite the KRAS G12D cells being fully resistant to the combination therapies used.

To confirm the changes in tumour cell subpopulation composition that were indicated by bioluminescence scans, we sampled five representative tumours of eligible treatment groups at the end of treatment for flow cytometric analysis for GFP/BFP expression in KRAS G12D or G12C mutant cancer cells. Tumours from combination-treated mice could not be sampled at this time point, as they had regressed to an extent that would not allow for obtaining sufficient sample material. Plotting the fraction of KRAS G12D mutant cancer cells out of all live cancer cells corroborated the findings indicated in bioluminescence scans that only KRAS G12Ci, but not vehicle, SHP2i or anti-PD-1 treatments led to the increased proliferation of the treatment-resistant KRAS G12D tumour subpopulation (Figure 2f). Additionally, to assess whether the bulk of the tumour mass after tumours had grown back after treatment in non-CRs was comprised of KRAS G12C or G12D mutant cancer cells, additional quantification of tumour cell fractions by flow cytometric analysis was also performed once mice had reached a humane endpoint. Unexpectedly, this revealed a high degree of variability in mice administered with G12Ci, either as monotherapy or in combination treatments (Figure S3f). While some tumours contained a mix of both tumour cell populations, other tumours, especially in the combination groups, were mostly comprised of either KRAS G12C or G12D cells. This might indicate that KRAS G12C mutant tumour cells can not only persist during treatment intervals, but further restore proliferation-related programmes and contribute to the tumour mass in relapsed tumours over time in some tumours.

Because complete responses were observed, despite the presence of tumour cells which did not respond to any of the administered treatments directly, we speculated that treatment-enhanced anti-tumour directed immunity may have contributed to the elimination of treatment-resistant cells. If immune suppression within the TME could be alleviated through inhibition of the oncogenic KRAS signalling while increasing immunogenic cell death within the KRAS G12C mutant treatment-responsive tumour compartment, this could prime an adaptive immune response to recognise shared epitopes on both cancer cell populations. Combination treatments with SHP2i or anti-PD-1 could further enhance these responses, thereby aiding immune-mediated elimination of drug-resistant subpopulations. To test this hypothesis, we rechallenged the CR mice, which had remained tumour-free off treatment for over a month, by injecting 150,000 KRAS G12D mutant tumour cells on the opposite flank (Figure 2g). Compared to a group of naive mice where the secondary tumours grew at a speed comparable to primary tumours of previous experiments, mice which were KPAR1.3 cell-experienced were either able to completely reject the reinjected KRAS G12D mutant cancer cells, or in other cases, tumours grew back slower compared to naive mice, indicating the involvement of an adaptive immune response and immune memory (Figure 2h).

Following these results, we wanted to explore whether treatment responses in our mixed preclinical model system could be further enhanced by extending the treatment interval from two to three weeks or via additional synergistic effects in combining G12Ci, SHP2i, and anti-PD-1 therapies. Assessing tumour growth in this experiment (Figure S4a), somewhat fewer CRs were observed in the dual treatment combination groups compared to the previous experiment, regardless of the extended treatment interval. Despite this, 12/14 mice in the triple-combination treatment group were CRs, indicating additional synergistic effects in this therapeutic regimen. This correlates with the bioluminescence scans showing a better control of the KRAS G12Ci-resistant tumour subpopulation at the end of the treatment (Figure S4b). CR mice in this experiment were also rechallenged by injecting KRAS G12D mutant cancer cells in the opposite flank, and results again implied the presence of anti-tumour-directed immune memory, capable of recognising shared antigens on both cancer cell populations (Figure S4c). We speculated that the observed increase in CRs in triple-combination-treated mice could result from additional therapy synergy between SHP2i and anti-PD-1. Indeed, treatment of GH-mice harbouring either KRAS G12C or G12D derived tumours with combined SHP2i plus anti-PD-1 revealed that a degree of tumour control during treatment was achieved in tumours derived from both isogenic cell lines (Figure S4d and S4e), although no CRs were observed in either group.

Taken together, these data indicate that combining KRAS G12C inhibitors with other immune-enhancing treatments could, through synergistic effects, enhance treatment responses in the presence of drug-resistant tumour subpopulations. Moreover, likely by bolstering anti-tumour immunity, combination treatments were able to promote the elimination of KRAS G12D mutant tumour subpopulations which did not respond to any of the individual lines of therapy directly.

### G12Ci induces early formation of proliferative clusters of G12D cells and extensive T cell infiltration

To visualise the spatial distribution of the KRAS G12Ci-sensitive and -resistant cells across the different treatment groups, we performed multiplexed immunofluorescence (IF) analysis, followed by segmentation and spatial analysis (Figure 3a). The antibody panel included anti-GFP, which served as pan-cancer cell marker capable of detecting both eGFP and eBFP, as well as anti-luciferase antibody to specifically identify Luc-eGFP KRAS G12D cells. We also included the T cell lineage markers CD8 and CD4 to assess changes in T cell infiltration. Mice were again engrafted with reporter-traced cancer cells composed predominantly of eBFP KRAS G12C cells with a minor subpopulation (0.04%) of Luc-eGFP KRAS G12D cells. Tumours were treated for six days, coinciding with the time point at which bioluminescence signal begins to pick up (Figure S2e). In contrast to the two-week time point, where KRAS G12Ci-treated tumours were almost entirely composed of KRAS G12D cells, tumours at this stage displayed a mixed population of both G12C and G12D cells (Figure S6a). For the analysis of the stained tissue sections, we developed a novel machine-learning based, high-throughput, computational pipeline to segment and quantify the indicated cell types and activation markers across all treatment groups for a total of 72 tumours (Figure S5). Comparing quantitative output of the automated segmentation to the original IF images, revealed a high degree of accuracy, with few false positive or false negative events.

**Figure 3.**
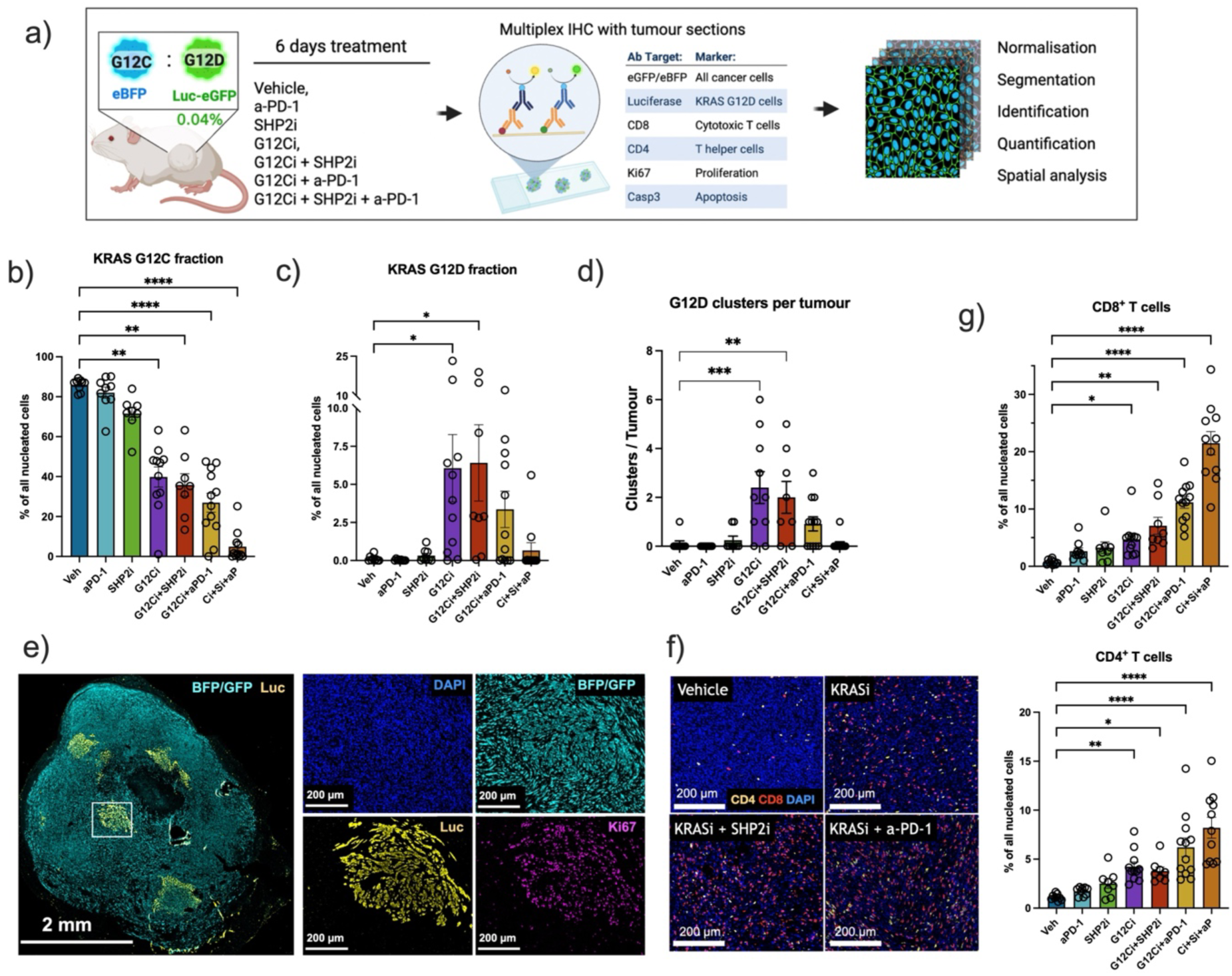
KRAS G12Ci treatment increases the proportion of KRAS G12D cells and T cell infiltration. a) Schematic of the experiment. Mixed subcutaneous tumours were engrafted with BFP KRAS G12C cells plus 0.04% Luc-eGFP KRAS G12D cells. Mice were treated for six days with the RAS G12C(ON) inhibitor RMC-4998 (100 mg/kg; G12Ci / Ci) with or without the SHP2 inhibitor RMC-4550 (30 mg/kg; SHP2i / Si) and/or a-PD-1 (10 mg/kg; aPD-1/aP). After treatment tumours were fixed and multiplex immunofluorescence was performed. Experiment was done twice to obtain two independent biological replicates. For quantifications shown in this figure, each dot represents one tumour, error bars indicate mean ± SD. One-way ANOVA between vehicle and each of the treatment groups was performed; only significant comparisons are indicated. b) Fraction of KRAS G12C mutant cancer cells out of all nucleated cells. c) Fraction of KRAS G12D mutant cancer cells out of all nucleated cells. d) Number of KRAS G12D clusters per tumour section, defined as cell aggregates with more than 100 G12D cells. e) Representative tumour section of a G12Ci-treated tumour section (left) with ROI of a proliferating KPAR1.3 KRAS G12D cluster (right panel). Markers: BFP/GFP as pan-cancer cell marker, Luciferase (Luc) demarcating KRAS G12D mutant cancer cells, Ki67 indicating proliferation. Scale bars indicated on respective images. f) Representative merge images from multiplex IF staining showing CD4^+^ T cells (yellow) CD8^+^ T cells (red) and DAPI; scale bar: 200 μm. Images selected do not include clusters of G12D cells. (g) Fractions of CD8^+^ (top) and CD4^+^ (bottom) T cells out of all nucleated.

As expected, cancer cell quantification revealed a reduction in KRAS G12C cells across different treatments (Figure 3b). Consistent with the bioluminescence data, treatment with KRAS G12Ci also led to an increase in the KRAS G12D fraction (Figure 3c and S6b). At this earlier time point there were not substantial differences between the KRAS G12Ci monotherapy and the combinations with either SHP2i or anti-PD-1. In contrast, the triple combination effectively eliminated both G12C and G12D cells, consistent with the outcomes observed in the survival experiment. Analysis of the spatial distribution revealed that tumours treated with KRAS G12Ci, either as monotherapy or in the double combinations, contained distinctive clusters of KRAS G12D cells (Figure 3d, 3e and S5c). On average, treated tumours contained only two KRAS G12D clusters per section (defined as clusters of more than 100 cells), suggesting clonal expansion of the KRAS G12Ci-resistant subpopulation. Additionally, the majority of the KRAS G12D tumour cells in G12Ci-treated tumours were Ki67-positive (∼60%), in contrast to KRAS G12C cells (∼20%), confirming an increased proliferative activity in the resistant subpopulation (Figure 3e and S6d). Interestingly, the proliferative and spatial patterns were similar between KRAS G12Ci monotherapy and the double combinations, suggesting that immune-mediated control may occur at later time points.

Spatial analysis also revealed increased infiltration of CD4^+^ and CD8^+^ T cells following KRAS G12Ci treatment, which was further enhanced by combination therapies, consistent with previous studies (Figure 3f and 3g)^6,34^. This was accompanied by an increase of Ki67-positive T cells (Figure S6e). Interestingly, combination treatments also led to an increase of cleaved-caspase-3 positive CD8^+^ T cells, although these remain an extremely small fraction of the total; this might reflect activation-induced cell death (Figure S6f). Increased T cell density correlated with depletion of KRAS G12Ci-sensitive cancer cells, an effect most pronounced in triple combination-treated tumours, where T cell numbers exceeded those of cancer cells (Figure S6g). Because G12Ci-resistant cancer cell clusters are highly proliferative and unaffected by KRAS inhibition, we next investigated their impact on T cell infiltration. Comparison of T cell density inside versus outside the resistant clusters showed similar CD8^+^ T cell infiltration, indicating that treatments also result in a CD8^+^ T cell enrichment inside the clusters compared with vehicle-treated tumours (Figure S6h). However, proliferative (Ki67-positive) CD8^+^ T cells were reduced within the clusters relative to surrounding areas (Figure S6i). In contrast, CD4^+^ T cells did not show increased infiltration within clusters compared with vehicle-treated tumours, revealing a distinct spatial pattern (Figure S6j).

Overall, the spatial analysis indicates that the minor G12Ci resistant subpopulation expands into highly proliferative clusters in response to KRAS G12Ci, whether given as monotherapy or in combination. Treatments also triggered a marked influx of T cells, which infiltrated both the G12Ci sensitive tumour bulk and, to a lesser extent, the resistant clusters. The selective loss of KRAS G12D cells seen in long term experiments with G12Ci combination relative to monotherapy (Figure 3) likely occurs later than this six-day treatment time point, and may be driven by an enhanced T cell response.

### Combination therapies increase inflammatory-associated signatures in KRAS G12Ci-resistant cells

To investigate how the different treatments impact KRAS G12Ci-sensitive and - resistant cells in a mixed tumour setting, we performed transcriptomic analysis of the two populations after five days of treatment with KRAS G12Ci, SHP2i, or the combination. To ensure sufficient material for downstream analysis, especially in the vehicle and SHP2i conditions, a higher proportion (7.5%) of KRAS G12D cells was co-engrafted. Following treatment, eBFP KRAS G12C and Luc-eGFP KRAS G12D cancer cells were sorted based on their fluorescent tags and subjected to bulk RNA sequencing (Figure 4a). Principal component analysis (PCA) revealed that both genotype and treatment contributed to variation in gene expression profiles (Figure 4b). As expected, vehicle-treated KRAS G12C samples clustered separately from inhibitor-treated counterparts (Figure S7a). Interestingly, treatment-driven clustering was also evident in KRAS G12D samples, suggesting that these cells undergo transcriptional changes in response to treatment, despite not being directly targeted by KRAS G12Ci (Figure S7b).

**Figure 4.**
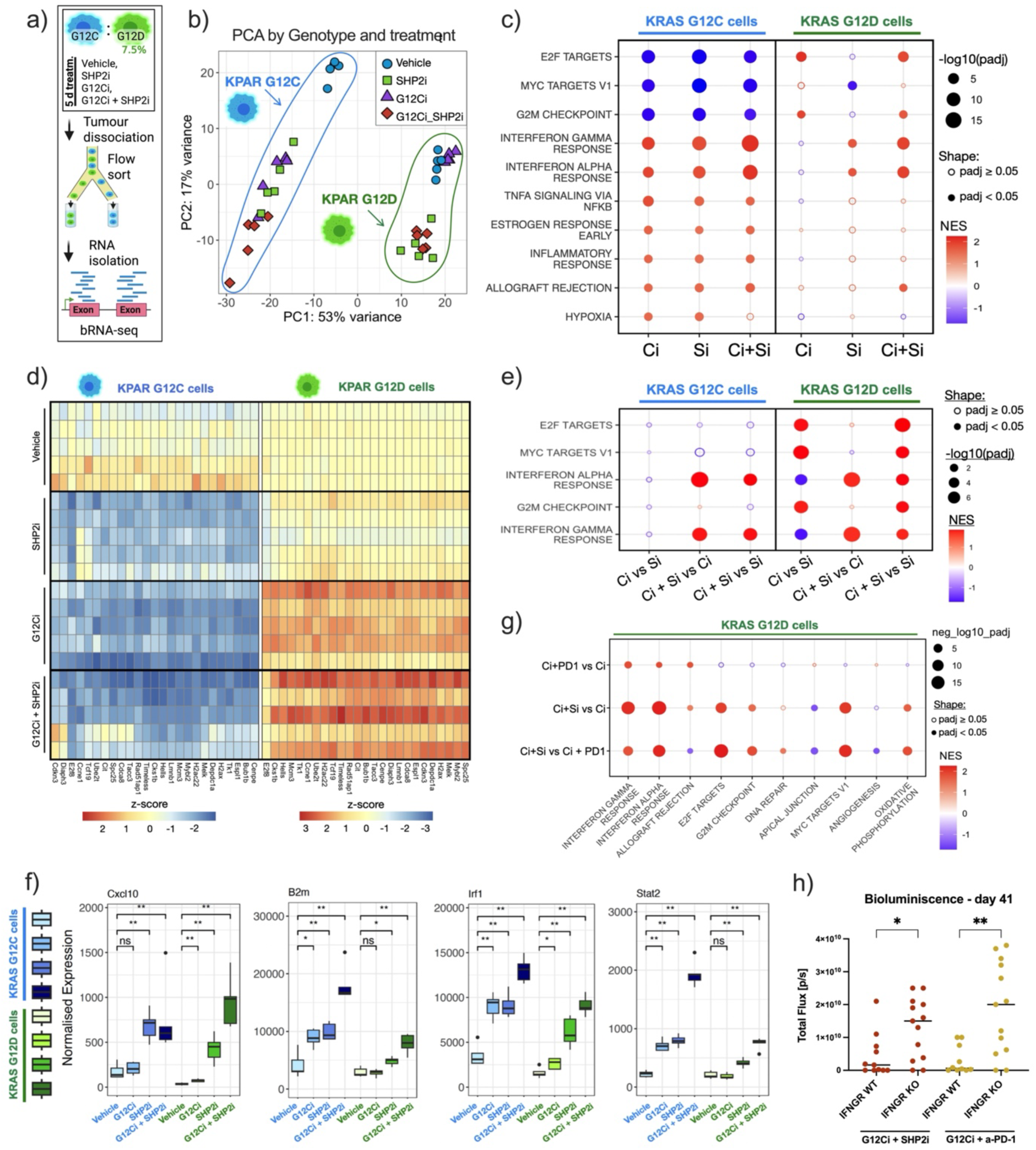
KRAS G12Ci-resistant subpopulation undergo transcriptional changes in response to KRAS G12Ci and combinations. a) Schematic of RNA-seq experiment. Mixed subcutaneous tumours were engrafted with BFP KRAS G12C cells plus 7.5% Luc-eGFP KRAS G12D cells. Mice were treated for five days with the RAS G12C(ON) inhibitor RMC-4998 (100 mg/kg; G12Ci / Ci) with or without the SHP2 inhibitor RMC-4550 (30 mg/kg; SHP2i / Si). After treatment cancer cells were sorted and bulk RNAseq was performed. b) PCA analysis across all samples and treatment groups; genotype of respective samples is indicated. c) GSEA of top 10 hits for differentially regulated gene sets based on msigdb hallmarks gene set collection, with vehicle set as baseline for each comparison indicated on x-axis. Normalised enrichment scores (NES) are indicated by colour scale and adjusted p-values (padj) are indicated by size of respective dots. Hits which were not significant padj > 0.05 indicated as unfilled circles. d) Heatmaps for top 20 leading edge genes of the E2F target collection from comparison between vehicle and G12Ci for KRAS G12C mutant (left) and KRAS G12D mutant cancer cells (right). The average of the 5 vehicle samples was taken to define a baseline for z-score calculations and was set to zero. A positive z-score (red) indicates increased expression of a gene while a negative z-score (blue) indicates decrease in expression, compared to the average expression across vehicle samples. e) GSEA of secondary treatment effects for KPAR G12C and G12D cancer cells. Second treatment on x-axis legends refers to baseline. Scoring as indicated in (c). f) Expression of individual genes related to inflammatory-response genes; with box plots indicate median, interquartile range, whiskers represent largest and smallest quartile and outliers are displayed as individual dots with figure legends consistent between plots. Statistical significance between vehicle-treated and inhibitor-treated samples was assessed using pairwise Wilcoxon rank-sum tests (two-sided). g) Mixed subcutaneous tumours were engrafted with BFP KRAS G12C cells plus 0.5% Luc-eGFP KRAS G12D cells. Mice were treated for five days with the RAS G12C(ON) inhibitor RMC-4998 (100 mg/kg; Ci) with or without the SHP2 inhibitor RMC-4550 (30 mg/kg; Si) or a-PD-1 (10 mg/kg; P). After treatment KRAS G12D cells were sorted and bulk RNAseq was performed. GSEA of top 12 hits for differentially regulated gene sets based on msigdb hallmarks gene set collection, for the comparison indicated in x-axis, the later term is set as baseline on respective plots. Scoring as indicated in (c). h) Mixed subcutaneous tumours were engrafted with BFP KRAS G12C cells plus 0.04% of Luc-eGFP KRAS G12D cells expressing wild-type (WT) or knock-out (KO) IFNGR and treated for two weeks with 100 mg/kg RMC-4998 in combination with 30 mg/kg RMC-4550 (G12Ci + SHP2i) or 10 mg/kg anti-PD-1 (G12Ci + a-PD-1). Graph shows the bioluminescence scans indicating the relative abundance of KRAS G12D cells 18 days after treatment withdrawn (day 41 since the start of the treatment). One-way ANOVA comparing WT vs KO groups for each the treatment group is shown.

To further characterise treatment-associated transcriptional changes in KRAS G12C and G12D cells, we performed gene set enrichment analysis (GSEA) using the Hallmark gene set collection (Figure 4c; Table S2). As expected, treatments suppressed the proliferation-associated signatures E2F targets and G2M checkpoint in KRAS G12C cells. In contrast, these same signatures were upregulated in KRAS G12D cells following KRAS G12C inhibition. Heatmaps of individual samples displaying leading-edge genes contributing to the E2F signature (Figure 4d) and selected proliferation-related genes (Figure S7c), further illustrated the opposing transcriptional responses between KRAS G12C and G12D tumour cell populations under KRAS G12Ci treatment. In contrast, SHP2 inhibition downregulated Myc target gene expression in both KRAS G12C and G12D cells, likely reflecting tumour cell-intrinsic effects of SHP2 inhibition. In fact, although SHP2i alone was not sufficient to inhibit the growth of KRAS G12D tumours (Figure 2a), it still suppressed the expression of RAS target genes both in vitro and in vivo, although the effect size tended to be smaller in G12D cells compared to G12C cells (Figure S7d and S7e).

As previously described, inflammatory response-associated signatures, including interferon-gamma (IFNγ) and interferon-alpha (IFNα) signalling pathways, were upregulated in KRAS G12C cells following treatments (Figure 4c) ^6^. Interestingly, these pathways were also induced in KRAS G12D cells in response to SHP2i or the combination therapy, suggesting that SHP2 inhibition may enhance inflammatory signalling in KRAS G12D mutant cancer cells. This effect could be caused by different mechanisms, including increased IFN secretion due to TME remodelling or modulation of cancer cell-intrinsic responses to IFN. Previous studies have shown that oncogenic KRAS suppresses IFNγ responses, and that KRAS or SHP2 inhibition can restore IFNγ-driven transcriptional programmes ^6,34^. However, our in vitro data indicate that this effect is less pronounced in KRAS G12D cells, consistent with their reduced sensitivity to SHP2 inhibition compared to KRAS G12C cells ^36^ (Figure S8a). These findings would suggest that the observed IFN pathway activation in KRAS G12D cells isolated from mixed tumours is likely mediated primarily by extrinsic signals from the TME. Moreover, the addition of KRAS G12Ci, despite not directly targeting KRAS G12D cells, further enhanced IFN pathway activation in these cells (Figure 4e), again supporting a role for non-cancer cell intrinsic factors in shaping the transcriptional response of KRAS G12D cells. Importantly, this included upregulation of genes involved in antigen presentation (e.g., H2-K1, B2m) and T cell recruitment (e.g., Cxcl9, Cxcl10) that could enhance immune recognition of the KRAS G12Ci-resistant cells (Figure 4f and S8b).

To further assess how combining KRAS G12Ci with SHP2i or anti-PD-1 alters the cell state of KRAS G12D subpopulations compared to G12Ci monotherapy, we performed a second RNA-seq experiment (Figure S8c). Since all treatment groups included KRAS G12Ci, which drives expansion of the KRAS G12D population, we were able to reduce the ratio of engrafted KRAS G12D cells to 0.5%, to more accurately recapitulate the emergence of resistant subpopulations as a subclonal event, while still enabling sufficient material for sequencing. GSEA analysis confirmed that the addition of SHP2i to KRAS G12Ci induced substantial transcriptional changes in KRAS G12D cells, including upregulation of interferon response signatures, in part driven by the effect of the SHP2i treatment (Figure 4g, S8d, S8e and Table S3). In contrast, the transcriptional profile of KRAS G12Ci combined with anti-PD-1 therapy clustered more closely with that of KRAS G12Ci monotherapy, as indicated by the limited gene expression changes (Figure S8e). Nevertheless, GSEA revealed that this combination also enhanced interferon signalling in KRAS G12D cells, although to a lower extent than the combination with SHP2i (Figure 4g).

Given that tumour cell intrinsic IFNγ has previously been shown to be required for antitumour immune responses^40^ and that both combination therapies increased IFN responses in the KRAS G12D cells, we asked whether this pathway contributes to the clearance of these KRAS G12Ci-resistant tumour subpopulations. To test this, we generated IFNGR knock-out (IFNGR KO) Luc-eGFP KRAS G12D cancer cells lacking the receptor for IFNγ and assessed their response to therapy in mixed co-engrafted tumours. Bioluminescence scans revealed that in tumours treated with either combination therapy, the IFNγ-insensitive KRAS G12Ci-resistant subpopulation expanded more rapidly (Figure S8f). This was validated when tumours were analysed after completion of treatment (Figure 4h). These findings suggest that the increased interferon responses observed in KRAS G12D cells may contribute to the immune-dependent control of the resistant subpopulation.

In summary, these data corroborate an enhanced proliferative signature in the treatment-resistant KRAS G12D tumour following KRAS G12C inhibition, either as single treatment or in combination. However, it is notable that combination therapies also enhanced inflammatory responses, including the upregulation of genes involved in interferon responses and antigen presentation, not only in KRAS G12Ci sensitive cells but also in the resistant subpopulation. This immunomodulatory effect may be indicative of a proinflammatory TME induced by the combination therapies and may contribute to the immune-mediated elimination of the treatment-resistant cells.

### Immune-phenotyping reveals treatment-driven shift towards a more inflamed TME

Given the increased proinflammatory signatures in KRAS G12C and G12D tumour cells (Figure 4), along with the higher T cell infiltration observed in the combination-treated tumours (Figure 3), we next investigated how the different therapies remodel the TME. To this end, we performed immune phenotyping by spectral flow cytometry using an antibody panel that included both immune lineage and cell state markers (Figure 5a). Consistent with our previous findings, 6-day treatment with KRAS G12Ci as monotherapy or in combination led to a reduction in the proportion of KRAS G12C cells, accompanied by an increase of KRAS G12D cells (Figure 5b). Additionally, cancer cells upregulated MHC-I and PD-L1 expression following SHP2i or combination therapies, consistent with the enhanced inflammatory signatures observed in these conditions (Figure S9a).

**Figure 5.**
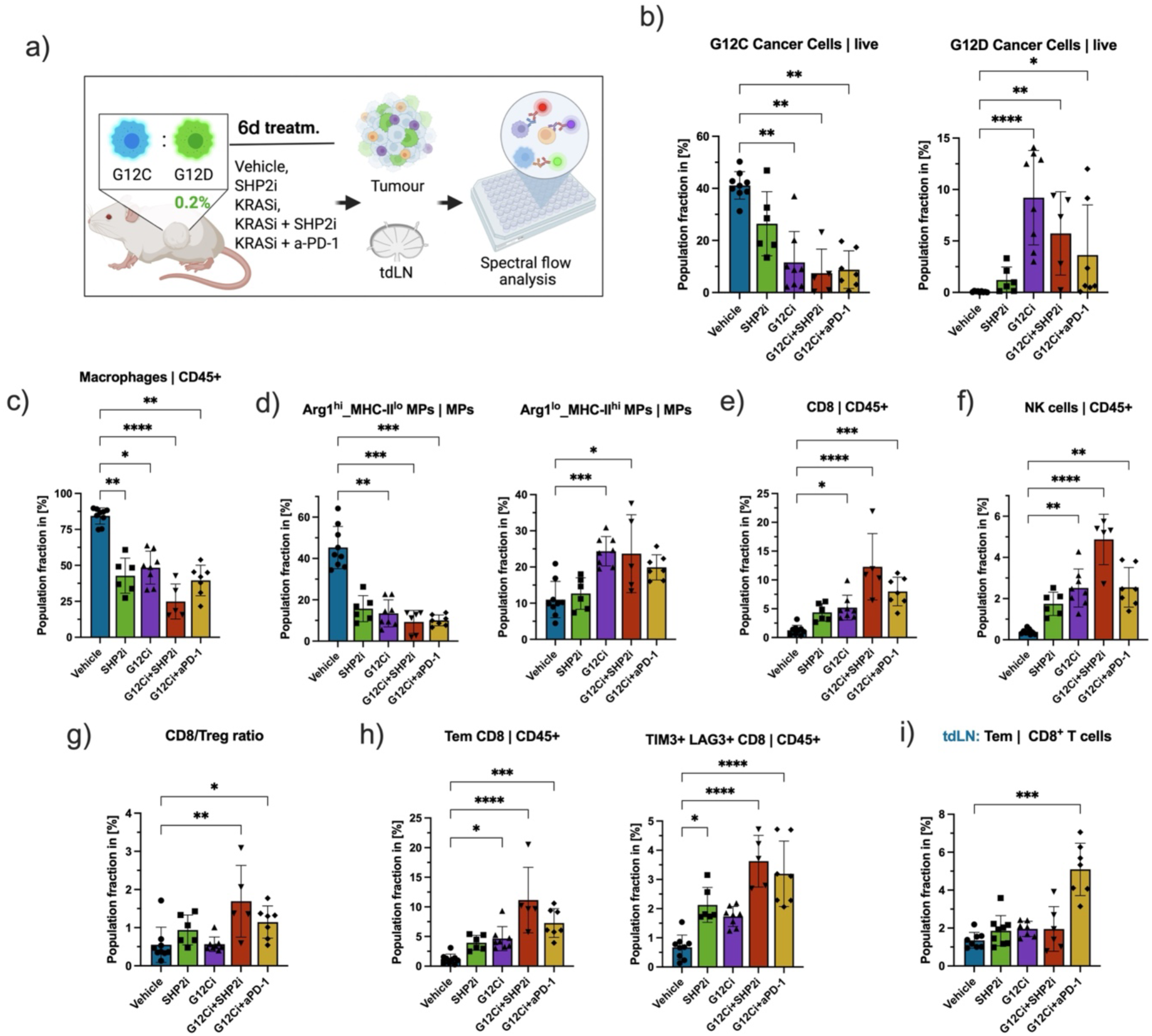
Treatments modulate the immune TME and increase inflammation. a) Schematic of experiment. Mixed subcutaneous tumours were engrafted with BFP KRAS G12C cells plus 0.2% Luc-eGFP KRAS G12D cells. Mice were treated for six days with the RAS G12C(ON) inhibitor RMC-4998 (100 mg/kg; G12Ci / Ci) with or without the SHP2 inhibitor RMC-4550 (30 mg/kg; SHP2i / Si) or a-PD-1 (10 mg/kg). After treatment flow cytometry of tumours and tumour-draining lymph nodes (tdLN) was performed. For all subsequent plots, each dot represents one individual tumour; bar graphs indicate mean ± SD; One-way ANOVA Kruskal-Wallis test comparing vehicle to each of the treated conditions; only significant comparisons are shown. b) Fraction of KRAS G12C (left) and KRAS G12D (right) mutant tumour cells out of all life cells. c) Fraction of macrophages (MPs) out of all immune cells (CD45^+^). d) Macrophage polarisation: Fraction of Arg1 high /MHC-II low (left) and Arg1 low/MHC-ll high (right) MPs out of all MPs. e) Fraction of CD8^+^ T cells out of CD45^+^ cells. f) Fraction of NK cells out of CD45^+^ cells. g) CD8^+^ T cells to Treg (CD4^+^ Foxp3^+^ T cells) ratio by treatment group. h) Fraction of effector memory (Tem; CD62L^-^/CD44^+^; left) and TIM3^+^/Lag3^+^ (right) CD8^+^ T cells out of CD45^+^ cells. i) In tumour draining lymph node (tdLN): fraction of Tem (CD62L^-^/CD44^+^) CD8^+^ T cells out of all CD8^+^ T cells.

Analysis of the immune cell compartment revealed that treatment resulted in a profound remodelling of the TME across multiple immune cell subpopulations (Figure S9b). Within the innate immune cell compartment, a significant reduction in macrophages was observed across all treatment groups, a phenotype that has been associated with a favourable clinical prognosis^41^ (Figure 5c). This was primarily driven by the decrease of Arg1^high^ MHC-II^low^ macrophages, which have been linked to more pro-tumorigenic functions, whereas the percentage of more pro-inflammatory Arg1^low^ MHC-II^high^ macrophages increased (Figure 5d). In addition to the changes in macrophage infiltration and polarization observed in all the treatment conditions, SHP2 inhibition also led to a significant increase in expression of the T cell co-stimulatory transmembrane protein CD86 (Figure S9c), suggesting enhanced immune activation and highlighting additional effects of SHP2 in the myeloid compartment, consistent with previous reports^35^.

Analysis of the lymphoid compartment corroborated the increase of CD8^+^ and CD4^+^ T helper cell infiltration observed in the spatial analysis (Figure 5e and S9d). This was accompanied by an increase of tumour infiltrating NK and NKT cells, which was particularly pronounced in the SHP2i combination group (Figure 5f and S9e). In parallel, regulatory T cells (Tregs) were also increased across treatment groups (Figure S9f), which could potentially dampen anti-tumour immune responses. However, the ratio of CD8^+^ T cells to Tregs, generally considered a favourable prognostic indicator, was elevated in the combination therapies (Figure 5g). Further characterization of CD8^+^ T cells revealed a significant increase in the fraction of effector memory (Tem; CD44+/CD62L^-^) as well as exhausted (Tim3^+^/Lag3^+^) T cells (Figure 5h). However, when expressed as percentages within the CD8^+^ compartment, these increases were not apparent, suggesting that the expansion of activated and exhausted CD8^+^ T cells primarily reflects the overall increase in the influx of T cells (Figure S9g).

To assess whether treatment-induced effects extended beyond the tumour site, we also performed immune phenotyping of inguinal tumour-draining lymph nodes (td-LN) adjacent to the subcutaneous tumours. This analysis revealed a significant increase in Tem CD8^+^ T cells in the mice treated with the combination of KRAS G12Ci and anti-PD-1, suggesting a potential systemic enhancement of T cell priming (Figure 5i). This effect was not observed in the KRAS G12Ci plus SHP2i combination, although we cannot exclude systemic immune modulation of this combination at later time points. These findings are consistent with the observed generation of immune memory capable of eliminating KRAS G12D cells in the tumour rechallenge experiments.

In summary, these results demonstrate that KRAS G12Ci, both as monotherapy and in combination, induces a profound remodelling of the immune TME. This includes a reduction and repolarization of macrophages and an increased infiltration of T and NK cells, as well as an expansion in memory T cells in tdLNs driven by the combination with anti-PD-1. Such enhancement of anti-tumour immunity could facilitate the immune-mediated targeting not only of KRAS G12Ci sensitive cells but also of resistant subpopulations.

### Adaptive immunity is required for durable responses and control of KRAS G12Ci-resistant subpopulations

The data presented above demonstrate the influx of T cells into KPAR G12C tumours upon treatment with KRAS G12Ci, but do not address whether or not these T cells are capable of specifically recognising the tumour cells. In order to address this, we took advantage of the fact that the major tumour associated antigen that determines the immunogenicity of KPAR tumours has been identified as the Emv2 endogenous retroviral envelope protein^18^. Tumour infiltrating T cells from mice bearing KPAR G12C tumours were analysed for their ability to bind to a synthetic tetramer containing the CD8 specific epitope from Emv2. The effect of treatments with G12Ci and combinations with SHP2i or anti-PD-1 were also measured (Figure 6a). T cells specifically recognising the CD8 epitope from Emv2 could be identified in wild-type tumours. In contrast, when mice were transplanted with KPAR1.3 tumour cells from which the Emv2 locus had been deleted, no such reactive T cells were found (Figure 6b and S10a). Although the percentage of CD8^+^ T cells specific for Emv2 did not increase with treatments (Figure S10a), there was an increase in their frequency following treatment with G12Ci combinations when measured as a proportion of total immune cells (Figure 6b). This was accompanied by an additional influx of CD8^+^ T cells not specific for Emv2 (Figure S10b). Therefore, the number of these Emv2 reactive T cells tracks the overall increase in total T cells, suggesting that their influx into the tumours is occurring by the same mechanism, likely cytokine driven. However, compared to the bulk T cell population, the Emv2 reactive T cells did show increased expression of the early activation marker CD69 and the activated cytotoxic T cell marker CD107a, suggesting a more activated phenotype than the non-Emv2-specific CD8^+^ T cells (Figure 6c).

**Figure 6:**
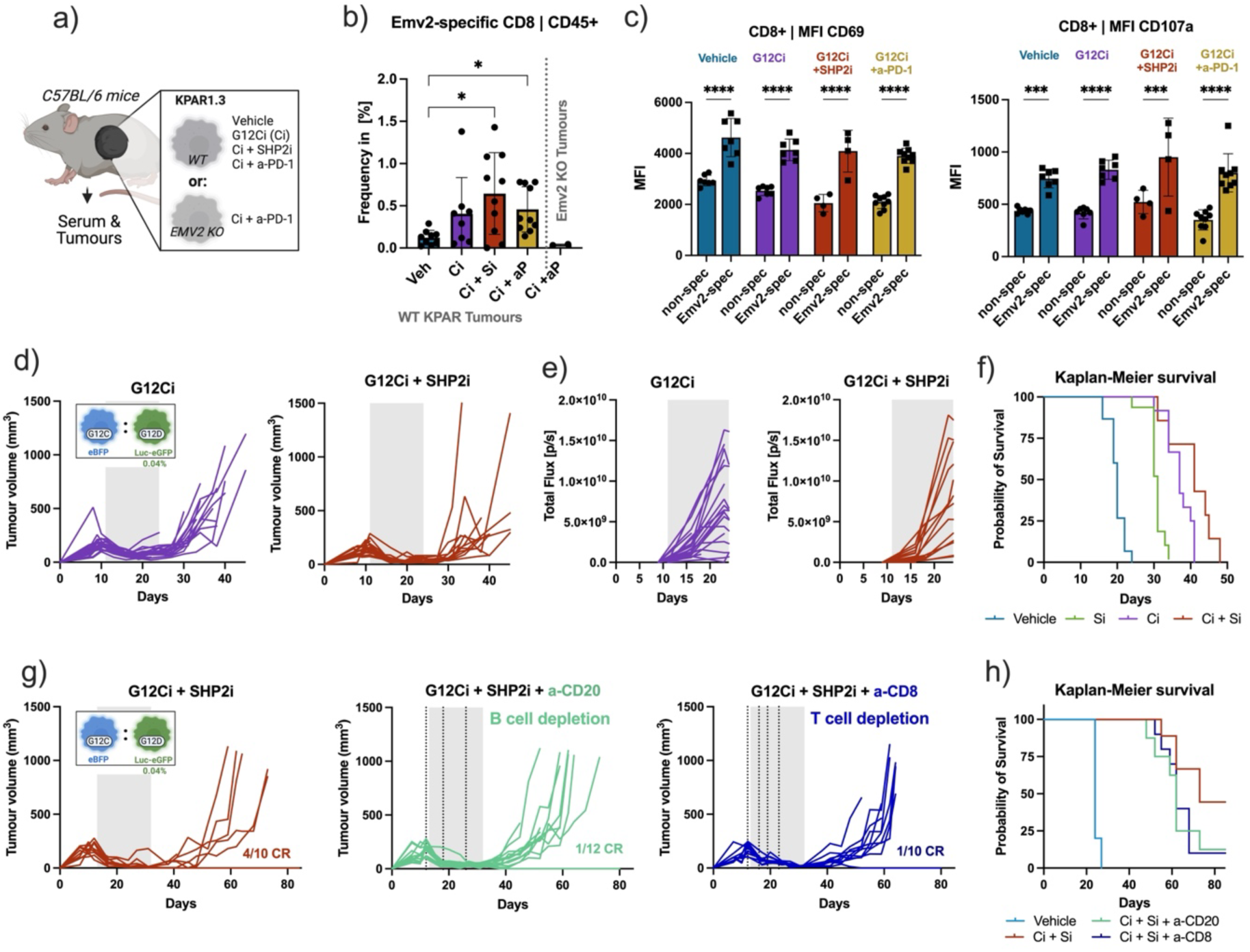
Adaptive immunity is required for durable responses. a) Schematic of experimental design for panels b and c. Mice with KRAS G12C KPAR1.3 wild-type or Emv2 knockout tumours were treated for six days with the RAS G12C(ON) inhibitor RMC-4998 (100 mg/kg; G12Ci / Ci) with or without the SHP2 inhibitor RMC-4550 (30 mg/kg; SHP2i) or anti-PD-1 (10 mg/kg; a-PD-1/aP). b) Fraction of Emv2-specific CD8^+^ T cells of all immune cells (CD45^+^). Each dot represents one individual tumour; bar graphs indicate mean ± SD; One-way ANOVA Kruskal-Wallis test comparing vehicle to each of the treated conditions; only significant comparisons are shown. c) Comparison in activation marker expression between Emv2-specific vs no-specific CD8^+^ T cells for T cell activation marker CD69 (left) and degranulation marker CD107a (right) from tumours treated as indicated in (a). (d-f) Mixed subcutaneous tumours were engrafted with BFP KRAS G12C cells plus 0.04% Luc-eGFP KRAS G12D cells in Rag1^-/-^ GH-mice. Mice were treated for two weeks (grey area) with the RAS G12C(ON) inhibitor RMC-4998 (100 mg/kg; G12Ci) and/or SHP2 inhibitor RMC-4550 (30 mg/kg; SHP2i). Graphs combine the replicate results of n=2 independent experiments. d) Tumour volume over time of individual tumours by treatment group. e) Bioluminescence scans indicating the relative abundance of KRAS G12D cells over time. f) Probability of survival, stratified by treatment group. (g-h) Mixed subcutaneous tumours were engrafted with BFP KRAS G12C cells plus 0.04% Luc-eGFP KRAS G12D cells in immune-competent GH-mice. Mice were treated for three weeks with either vehicle or G12Ci + SHP2i (100 mg/kg RMC-4998 + 30 mg/kg RMC-4550) (grey area). In two of the treated groups, either B cells or CD8^+^ T cells were selectively depleted by administering anti-CD20 (10 mg/kg) or anti-CD8 (12.5 mg/kg) antibodies, respectively; doses indicated by the vertical lines. g) Tumour growth over time for indicated treatment groups. Complete responders (CR) indicated on respective graphs. h) Probability of survival, stratified by treatment group.

To determine whether a primed adaptive immune response is essential for eliminating treatment-resistant KRAS G12D tumour subpopulations and achieving durable tumour control, we generated a strain of albino Rag1^-/-^ GH mice, which lack both functional T and B cells. Similar to the survival experiment performed in immunocompetent GH-mice (Figure 2c-e), mixed subcutaneous tumours containing a small fraction of KRAS G12D cells (0.04%) were engrafted and treated with either KRAS G12Ci, SHP2i or the combination. While initial tumour growth responses were comparable to those observed in immunocompetent mice, all tumours relapsed following treatment withdrawal and no complete responses were achieved (Figure 6d and S10c). Moreover, bioluminescence imaging revealed that, unlike in immunocompetent mice, the combination treatment failed to control the KRAS G12D subpopulation, which proliferated at a similar rate to in mice receiving KRAS G12Ci as monotherapy (Figure 6e and S10d). No long-term survivors were observed, unlike in the immunocompetent setting (Figure 6f, compare to Figure 2d). These findings indicate that adaptive immunity is required to control the growth of KRAS G12Ci-resistant subpopulations and to achieve durable therapeutic responses.

To dissect the relative contributions of T and B cells in mediating this immune control, we performed specific depletion experiments in immunocompetent mice. Interestingly, depletion of either CD8^+^ T cells or B cells individually reduced, but did not eliminate, the generation of complete responses when mice were treated with the combination of KRAS G12C plus SHP2 inhibitors (Figure 6g and 6h), suggesting that both cell types contribute to the elimination of KRAS G12D cells and are required for the generation of a durable anti-tumour immunity. B cell infiltration into the tumours was minor in our model (Figure S9b), so the impact of targeting B cells may be due to effects on promotion of T cell function or antigen presentation, or on production of tumour binding antibodies, which have been reported to play a significant role in this tumour model ^17^. Indeed, serum from mice bearing parental KPAR G12C tumours contained tumour-reactive antibodies that bound both KPAR1.3 G12C and G12D cells lines in vitro (Figure S10e). These antibodies did not bind KPAR1.3 cells lacking the Emv2 locus and were absent in mice bearing these Emv2 knockout tumours, confirming that the tumour-reactive antibodies are specific for the Emv2 endogenous retroviral envelope protein.

These data indicate that elimination of drug-resistant cancer cells on treating mixed tumours with G12Ci combination therapies is dependent on an adaptive immune response, likely involving both T and B cell functions. Emv2 tumour associated antigen specific CD8 T cells are found in tumours and increase in response to these treatments.

### Immune-mediated bystander killing affects tumours with diverse mechanisms of G12Ci drug resistance

The preclinical murine model system used so far recapitulates one of the resistance mechanisms observed in clinical samples from patients treated with KRAS G12C inhibitors^25^, specifically the presence of a minor subclonal population harbouring an alternative KRAS mutation that renders it intrinsically insensitive to KRAS G12C inhibition due to lack of inhibitor binding. However, this model does not reflect adaptive resistance mechanisms, as the cells carrying the KRAS G12D mutation have not undergone selective pressure from prior KRAS G12Ci exposure. To explore whether combination therapies could induce immune-mediated bystander elimination of minor resistant subpopulations arising through alternative resistance mechanisms, we generated a KRAS G12Ci-resistant tumour cell line by treating C57Bl/6 mice bearing orthotopic lung tumours derived from KPAR1.3 KRAS G12C cells with adagrasib five days a week until resistance emerged (Figure S11a). Clonal cell lines were subsequently established from resistant tumours and following initial screening a suitable subclone, named KPAR.E2, was selected for downstream analysis. Sanger sequencing of the KRAS transcript confirmed that the KRAS G12C mutation was retained, and no additional alterations were detected, including previously reported mutations in H95 or Y96 known to disrupt inhibitor binding (Figure S11b)^24^. Viability assays confirmed that KPAR.E2 cells were resistant to both inactive- and active-state KRAS G12C inhibitors (Figure 7a). Furthermore, the cell line was also resistant to MEK inhibition, while remaining sensitive to PI3K inhibition (Figure S11c). Finally, assessment of the response to adagrasib in vivo showed that, in contrast to the parental KPAR1.3 KRAS G12C cell line, KPAR.E2-derived subcutaneous tumours continued to progress on treatment (Figure S11d and S11e).

**Figure 7:**
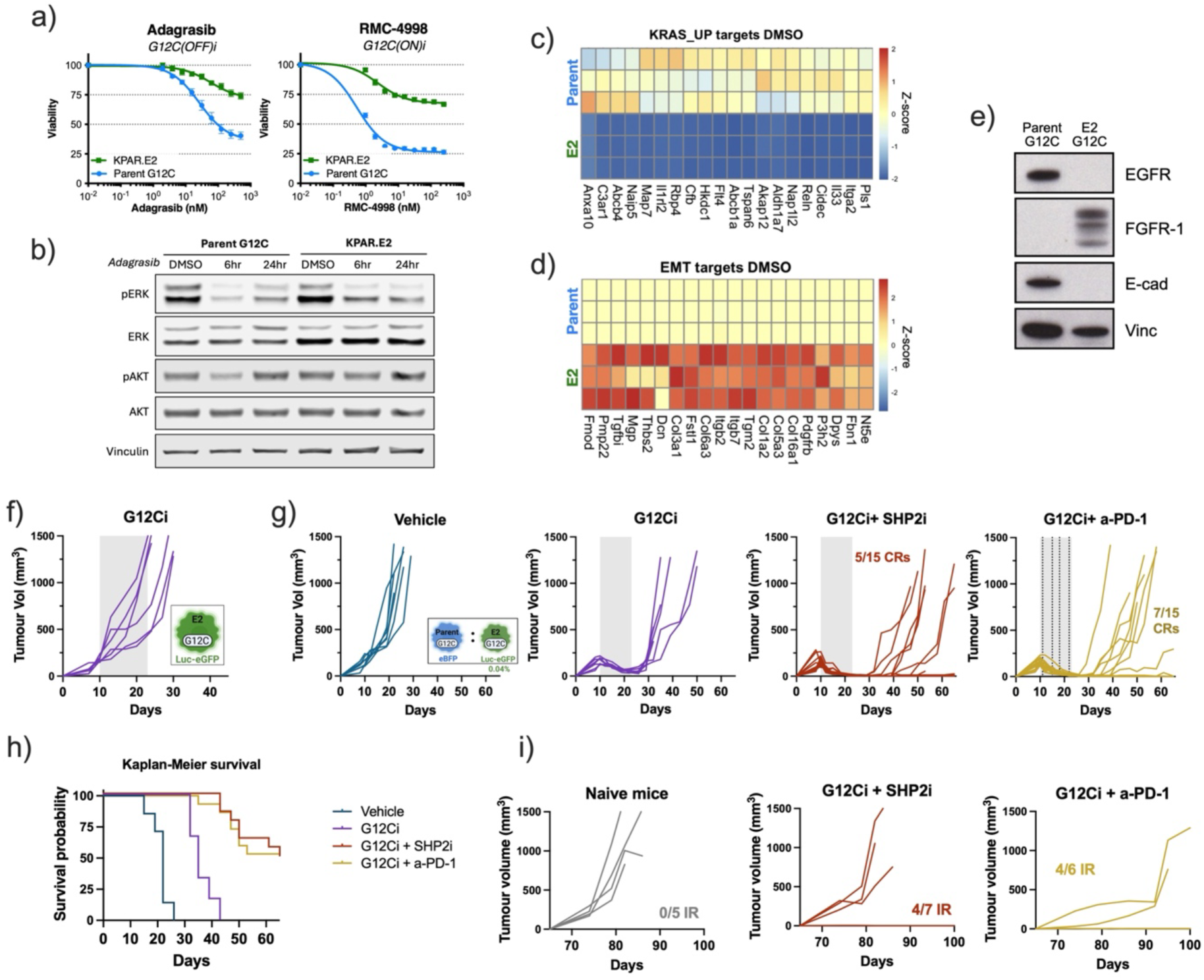
Immune-rejection of mechanistically distinct G12Ci-resistant tumour subpopulations. a) Viability assays of the KPAR.E2 cell line compared with the parental KPAR.1.3 KRAS G12C cells (Parent G12C). Cells were treated for 72 hours with 2-fold serial dilutions of the inactive or active-state KRAS G12C inhibitors, adagrasib and RMC-4998, respectively. Error bars indicate mean ± SEM, n = 3. b) Westen blot of cells treated at different time points with 100 nM adagrasib. c-d) RNA-seq analysis comparing gene expression at baseline (DMSO-treated samples) between parental KPAR1.3 KRAS G12C cell line (Parent) and G12Ci-resistant KPAR.E2 cells. Heatmaps for leading edge genes in GSEA analysis for hallmarks gene set collections c) KRAS_UP signature and d) EMT target signature. For both plots, z-score was normalised to average gene expression in “Parent” experimental group. e) Western blot comparing basal protein expression between parental KPAR1.3 and KPAR.E2 cells, using the indicated antibodies. f) Tumour growth of Luc-eGFP KPAR.E2 subcutaneous tumours treated for 2 weeks (grey area) with RMC-4998 (100 mg/kg; G12Ci). Growth of vehicle-treated mice is shown in Figure S13a. (g-h) Mixed subcutaneous tumours were engrafted with BFP KRAS G12C cells plus a fraction of 0.04% Luc-eGFP KPAR.E2 cells. Mice were treated for two weeks with the RAS G12C(ON) inhibitor RMC-4998 (100 mg/kg; G12Ci) with or without the SHP2 inhibitor RMC-4550 (30 mg/kg; SHP2i) or a-PD-1 (10 mg/kg). g) Tumour growth of individual tumours.Complete responders (CR) indicated in respective plots. h) Survival probability across different treatment groups, cut-off value: 500 mm^3^. i) Re-challenge of complete responders (CRs) from panel (g) with untraced, KPAR.E2 cells injected on day 65, compared to a group of age-matched naive control mice. Immune-rejected (IR) secondary tumours are indicated as fraction of all CRs. Graph titles indicate the treatment that the primary tumour received.

To investigate the mechanism of resistance, we evaluated the effect of KRAS G12C inhibition on downstream signalling. ERK activity was suppressed in response to KRAS G12Ci treatment and, unlike the parental cell line, KPAR.E2 cells did not exhibit a rebound in ERK phosphorylation at 24 hours, a well-described compensatory mechanism to KRAS inhibition (Figure 7b and S11f)^27^. These results indicate that resistance in KPAR.E2 is not driven by reactivation of the RAS-MAPK pathway. To further characterise the mechanisms underlying resistance to KRAS G12Ci, despite intact suppression of downstream signalling, we performed RNA sequencing on both KPAR.E2 and the parental KPAR1.3 cell lines, at baseline and following adagrasib treatment. PCA analysis revealed a clear separation of samples based on both genotype and treatment (Figure S12a). Consistent with the in vitro data, KRAS G12C inhibition led to suppression of transcriptional targets downstream of RAS signalling in both cell lines (Figure S12b). However, GSEA indicated that KPAR.E2 cells displayed lower basal KRAS signalling (Figure 7c and S12c). In addition, GSEA also revealed significantly enriched epithelial-to-mesenchymal transition (EMT)-associated gene signatures in KPAR.E2 compared with parental cells (Figure 7d and S12d), which was further supported by the upregulation of canonical EMT markers (Figure 7e and S12e). Enrichment of mesenchymal features has previously been associated with reduced KRAS dependency ^42^, and EMT induction is a known mechanism of resistance to KRAS and other MAPK pathway inhibitors^28,43^. Moreover, in line with previous findings, EMT was accompanied by rewiring of receptor tyrosine kinase expression and while the parental KPAR1.3 cells expressed EGFR, the resistant KPAR.E2 cells expressed FGFR-1 (Figure 7e)^44,45^. Altogether, these findings suggest that the KPAR.E2 cell line has undergone EMT and lost its dependency on RAS signalling, thereby conferring resistance to KRAS G12Ci. Therefore, this new cell line provides a suitable model to study an alternative and clinically relevant mechanism of KRAS G12Ci resistance in our co-engraftment tumour setting.

Next, we transduced the KPAR.E2 cell line with the Luc-eGFP reporter to allow tracking when engrafted as a KRAS G12Ci-resistant subpopulation. Treatment of Luc-eGFP KPAR.E2 tumours confirmed the resistance to the active-state RAS(ON) inhibitor RMC-4998 (KRAS G12Ci) (Figure 7f) along with only modest response to anti-PD-1 and SHP2i treatments, with all tumours progressing following treatment withdrawal (Figure S13a and S13b). After establishing the KPAR.E2 cells as a suitable alternative KRAS G12Ci-resistant subpopulation, eBFP KRAS G12C subcutaneous tumours containing a small fraction (0.04%) of Luc-eGFP KPAR.E2 cells were engrafted and treated with KRAS G12Ci alone or in combination with SHP2i or anti-PD-1 for two weeks. No complete responders were observed with KRAS G12Ci as monotherapy, and all tumours rapidly progressed after treatment withdrawal. In contrast, both combination therapies led to complete responses despite the presence of the KPAR G12Ci-resistant subpopulation (Figure 7g). These effects were also reflected in improved survival (Figure 7h). Moreover, consistent with the previous model, bioluminescence imaging showed that combinations effectively controlled the outgrowth of the resistant cells (Figure S13c). Furthermore, mice achieving complete responses presented an enhanced immune memory compared to naïve mice after tumour rechallenge in the opposite flank (Figure 7i), supporting the role of adaptive immunity in eliminating resistant cells and establishing long-term immune protection.

These data show that the immune-mediated bystander killing of drug-resistant tumour cells is not limited to a single mechanism of resistance, but can also be seen in a setting where tumour cells have become resistant to therapy in vivo due to a phenotypic switch from epithelial to mesenchymal state, raising the prospect of more general applicability of this therapeutic approach to tackling drug resistance.

## DISCUSSION

While initial clinical responses to recently approved RAS inhibitory drugs have been encouraging, it has become clear that acquired drug resistance is a massive problem that is severely limiting the clinical utility of these agents. Relatively rapid evolution of drug resistance has been a serious issue impacting the effectiveness of many targeted oncology drugs: this urgently needs to be addressed if these agents are to achieve lasting tumour control. In the case of mutant KRAS targeted agents, a huge diversity of resistance mechanism has been identified, making it hard to design combination therapies that effectively shut off signalling pathways within the cancer cells that are being exploited during the evolution of drug resistance.

In this study, we hypothesised that a better approach to tackling the problem of the rapid development of resistance to KRAS oncoprotein targeted drugs might be to exploit the fact that KRAS inhibitors at least partially reverse the KRAS pathway induced immunosuppressive tumour microenvironment in order to promote immune mediated attack on the drug-resistant cancer cells. Indeed, it has been shown that KRAS inhibitors cause changes in the tumour microenvironment that tend to reduce the ability of the cancer cells to evade immune mediated attack^4^: this should impact both the bulk drug-sensitive tumour and also minor drug-resistant subclones sharing the same tumour antigens and located within the same tumour. In addition, the killing of large numbers of cancer cells by the KRAS inhibitors is likely to result in the elevated uptake of tumour antigens by antigen presenting cells, which may provide a further “in situ” vaccination effect to boost the immune response to the tumour.

To explore this possible approach to tackling the emergence of drug resistance, we have developed a mouse cancer model that allows the study of the behaviour under selective pressure of minor drug-resistant clones within a bulk drug-sensitive tumour. We have focused particularly on the use of treatments combining KRAS inhibitors and agents that, at least in part, target the immune system. Based on the p53 deleted KPAR1.3 murine lung cancer cell line^18^, which on syngeneic transplantation forms tumours with a relatively inflamed (“hot”) tumour microenvironment, the KRAS oncogene has been engineered to either be sensitive (G12C) or insensitive (G12D) to KRAS G12C inhibitory drugs. The two populations of tumour cells can be distinguished by different fluorescent markers (BFP or GFP) and, in the case of the drug-resistant cells, growth can be followed in vivo using the expression of luciferase.

Using this system, we see that monotherapy with G12Ci results in a rapid outgrowth of the minor resistant cancer cell population, comparable to the responses often observed in patients. By six days of treatment, small islands of drug-resistant cells are seen to emerge which are proliferating, while the bulk drug-sensitive tumour has lost proliferative capacity. The outgrowth of drug-resistant cells can be visualised by bioluminescence and within two weeks the cancer cell population is almost entirely made up of KRAS G12D cells, even when they made up only as little as 0.01% of the original tumour. The drug-resistant cells grow more rapidly when the tumour is under selective pressure from G12Ci, likely due to removal of competition from the bulk drug-sensitive population, although it is also possible that trophic factors released from dying drug-sensitive cells may promote the growth of drug-resistant cells^46^.

Use of G12Ci monotherapy leads to profound remodelling of the TME of sensitive tumours, with influx of CD8^+^ T cells and NK cells, and changes in the phenotypic state of macrophages within the tumour, with reduction in immune suppressive cells (Arg1^high^ MHC-II^low^) and increase in inflammatory macrophages (Arg1^low^ MHC-II^high^). Despite this, G12Ci monotherapy is not sufficient to allow effective immune control of tumours with minor populations of drug-resistant cells as the G12D cells proceed to grow out, re-establish an immunosuppressive TME and form drug-resistant tumours that will lead to the death of the animal. However, when an immune targeted agent, such as anti-PD-1 immune checkpoint blockade or SHP2 inhibitor, which acts both on the TME and also the RAS pathway within the tumour cells, is added to the G12Ci, complete destruction of the drug-resistant, as well as the drug-sensitive, tumour cells is achieved in a high proportion of animals. This is despite the fact that the drug-resistant cells growing in the absence of drug-sensitive cells are not sensitive to G12Ci combinations with anti-PD-1 or SHP2i. This effect requires an adaptive immune system, with contributions from both T and B cells. We therefore consider this to be an immune-mediated bystander killing of the drug-resistant cells induced when the drug-sensitive cells are killed.

Mechanistically, it is likely that this immune-mediated bystander killing of the drug-resistant cells is at least in part mediated by the targeting of tumour associated antigens shared between the closely related drug-resistant and drug-sensitive cancer cell populations. In the case of the KPAR1.3 cells, we believe the major tumour associated antigen recognised by the immune system is the Emv2 envelope protein of the MLV endogenous retrovirus^18^, which can be recognised both by CD8^+^ and CD4^+^ T cells and also by antibodies. The proportion of Emv2 reactive CD8^+^ T cells within the tumour increases with G12Ci and combination treatments, providing a possible mechanism for elimination of drug-resistant tumour cells.

Transcriptomic analysis of the separate populations of drug-sensitive and drug-resistant cells within the tumour reveal that the drug-sensitive KRAS G12C cells go out of cycle in response to G12Ci and combinations, while drug-resistant KRAS G12D cells do not, as expected. In addition, drug-sensitive KRAS G12C cells demonstrate transcriptional responses characteristic of interferon signalling, as we have reported previously^6^. Interestingly, KRAS G12D show a stronger interferon response to G12Ci plus SHP2i compared to SHP2i alone, suggesting that the enhanced interferon signalling in these drug-resistant cells is due to the action of G12Ci on the drug-sensitive cells nearby. Removal of IFNγ receptor from these KRAS G12D makes them less responsive to immune-mediated bystander killing, indicating that this is at least in part dependent on the tumour cell–intrinsic ability to respond to IFNγ. While some of the interferon production in response to G12Ci will be coming from the drug-sensitive cancer cells^6^, it is likely that other cells within the TME that infiltrate the tumour in response to KRAS G12Ci are also contributing to this.

Further research will be needed to understand the full molecular details for the enhanced immune-mediated bystander killing of drug-resistant cancer cells achieved by combinations of G12Ci with anti-PD-1 or SHP2i, compared to G12Ci alone, but the involvement of tumour associated antigen specific T cells and also interferon signalling is clear. We have extended the model system to include a different mechanism of drug resistance, that caused by the KRAS G12C cells undergoing epithelial to mesenchymal transition under prolonged drug selection, and find that the immune mediated-bystander killing can still be seen in this setting, even though the two cell populations will be markedly more divergent than in the original model, although still sharing the Emv2 tumour associated antigen.

In terms of more immediate clinical translation, while combination of G12Ci with SHP2i potently promotes elimination of drug-resistant cells in this mouse model, it now appears to be relatively unlikely that SHP2 inhibitory drugs will enter routine clinical use in the foreseeable future due to significant toxicity problems. On the other hand, anti-PD-1 immune checkpoint blockade is used very widely, although trials of combination therapies with first generation KRAS G12C inhibitors have been limited by severe toxicity, particularly immune mediated hepatotoxicity^47^. However, it is likely that these toxicities are off target with regard to KRAS G12C, so there is reason to be hopeful that combinations of anti-PD-1 immune checkpoint blockade with more specific, second generation KRAS G12C inhibitors, such as divarasib, olomorasib and elironrasib, may allow the clinical exploration of the potential of combination KRAS G12C inhibition and anti-PD-1 to control the development of drug-resistant disease in the relatively near future. The potential for this approach to impact on drug-resistant disease is likely to be best in the setting of tumours with an inflamed TME, comparable to the model used here.

We believe that the model system described here firmly establishes the phenomenon of immune-mediated bystander killing of drug-resistant cancer cells. It also provides a platform for determining optimal approaches to enhancing the strength and impact of this effect, which will be particularly important for more immune evasive tumours that will not be so responsive to the combination of KRAS G12Ci with anti-PD-1. It is possible that adding further immune targeted agents may significantly potentiate the effect, such as using CTLA4 or CCR8 depleting antibodies to remove intratumoural regulatory T cells, which we know increase in response to G12Ci treatment^48^. The effect of targeting other aspects of the cancer immunity cycle could also be explored in this system, such as CXCR1/2 inhibitor for immunosuppressive myeloid cells and 41BB agonistic antibody for dendritic cells^49^. Alternatively, and more speculatively, the process of presentation of tumour associated antigens from dying cancer cells to immune effector cells may be another area that could be optimised, for example by intervening in phagocytosis or making the mechanism of cell death more immunogenic. This may represent a paradigm for eliminating the development of disease that is resistant to oncogene targeted agents through engagement of the immune system.

## MATERIALS AND METHODS

### Cell culture conditions and treatments

KPAR1.3 G12D and KPAR1.3 G12C cell lines were generated as previously described^18^. IFNGR knock-out and Emv2 knock-out KPAR1.3 G12D cells were generated as previously described^9,17^. All KPAR1.3 cell lines, including derivatives established for this study, were maintained in DMEM (Gibco, 41966052) supplemented with 10% fetal calf serum (FCS), 4 mM Glutamine, and 100 U/ml Penicillin/Streptomycin and were passaged using trypsin (Gibco, 25200072). Cell lines were tested regularly for mycoplasma by the Cell Science facility at the Francis Crick Institute.

For the different in vitro experiments, cells were plated at appropriate densities and treated 24 hours after seeding. RMC-4998 was provided by Revolution Medicines under a sponsored research agreement. RMC-4550 was provided by Revolutions Medicines under a former collaboration agreement with Sanofi. Adagrasib (MRTX849) was obtained from MedChemExpress and trametinib (GSK1120212) and pictilisib (GDC-0941) from Selleckchem. Recombinant mouse IFNγ was obtained from Biolegend.

### Generating reporter-traced KPAR1.3 cell lines

The lentiviral plasmids pLARRY-eGFP (#140025) and pFUGW-FerH-ffLuc2-eGFP (#71393) were obtained from Addgene. The nucleotide sequence of pLARRY-eGFP was altered to encode eBFP by substitution of two individual nucleotides via site-directed mutagenesis using the In-Fusion Snap Assembly Master Mix (TaKaRa, 638947) and the primers Tyr67 to His (Fwd ccctgacccacggcgtgcagtgcttcag; Rev cgccgtgggtcagggtggtcacgagg) and Tyr146 to Phe (Fwd gtacaacttcaacagccacaacgtctatatcatg; Rev ctgttgaagttgtactccagcttgtgccc). PCR conditions for rolling cycle amplification were performed following manufacturer’s instructions. Edited plasmids were transformed into *Stbl2* E. coli using a standard heat-shock protocol for selection and subsequent plasmid purification.

Reporter constructs were introduced into KPAR1.3 cells via lentiviral transduction. Lentivirus were generated by co-transfecting HEK293T cells with reporter plasmids (pLARRY-eBFP, pFUGW-FerH-ffLuc2-eGFP) and packaging plasmids (pCMV-dR8.2, pCMV-VSV-G) using the jetPRIME® transfection kit (Polyplus, 101000015). After 48 hours, virus-containing supernatant was collected, filtered (0.45 μm), and used for spinning infection of KPAR1.3 cells (600 rpm, 90 min, RT). Fluorescent reporter-expressing cells were single-cell cloned by FACS into 96-well plates.

### Characterising reporter-traced KPAR1.3 subclones in vitro

For growth-rate, viability, and luciferase activity assays, cells were plated into cell culture-treated 96 well plates with 1500 cells in 100 μl of the standard cell culture media, described above. For viability assays, inhibitors were added in serial dilution, as indicated in respective experiments. Viability was measured by adding 5 µl CellTiter-Blue® (Promega, G8081), incubating for 90 min at 37°C, and detecting fluorescence with an EnVision® 2105 multimode plate reader (PerkinElmer). For growth-rate assays, confluency was monitored using the S3 IncuCyte (Sartorius) at 20× magnification. Wells were imaged every 6 hours for up to 5 days or until confluency. Confluency over time was quantified using the S3 Live-Cell Imaging & Analysis Software (Sartorius). In *vitro* luciferase activity was assessed using the Pierce™ Firefly Luciferase Glow Assay Kit following manufacturer’s instructions (Thermo Fisher, 16176). Bioluminescence was measured with an EnSight® multimode plate reader (PerkinElmer).

### Western blotting

Cells were plated in 6-well plates in standard culture media with adjusted FCS concentrations as indicated in respective experiments. At the end of the treatment, cells were lysed in Cell lysis buffer (Cell Signaling, 9803) supplemented with Complete Mini protease inhibitor cocktail and PhosSTOP phosphatase inhibitors (Roche). Lysates were purified via centrifugation at 15,000 rpm for 15 min at 4°C, supernatant was collected, and protein concentration was measured for normalisation using the DC protein assay kit (Bio-Rad, 5000111). 20 µg of protein was then prepared with 4× NuPAGE LDS Sample Buffer (Invitrogen, NP0007), separated on NuPAGE 4-12% Bis-Tris gels (Invitrogen), and transferred to PVDF or nitrocellulose membranes (Merck Millipore), for chemiluminescence or fluorescence detection, respectively. Membranes were incubated with the following primary antibodies: from Sigma-Aldrich: Vinculin, #V9131; from Cell Signaling: AKT, #2920; pAKT (S473), #4060; ERK, #9107; pERK1/2 (T202/Y204), #9101; FGFR1, #9740; E-cadherin, #14472; from Abcam: EGFR, ab52894. Bound primary antibodies were incubated with the appropriate HRP-conjugated secondary antibodies and protein expression was detected via chemiluminescence using Immobilon Western HRP substrates (Merck). Alternatively, membranes were incubated with secondary antibodies compatible with infrared detection at 700nm and 800nm and scanned using the Odyssey Infrared Imaging System (Odyssey, LICOR).

### Quantitative reverse-transcription PCR

RNA was extracted using the RNAeasy Mini Kit (Qiagen) following manufacturer’s instructions. cDNA was generated using the Maxima First Strand cDNA synthesis kit (Thermo Fisher Scientific). qPCR was performed using Fast SYBR Green reagents (Applied Biosystems). Gene expression changes relative to the housekeeping genes were calculated using the ΔΔCT method. List of primers used is shown in Table S4.

### In vivo techniques

All in vivo experiments were conducted in accordance with a UK Home Office-approved project licence and following the welfare guidelines provided by the biological research facility at the Francis Crick Institute. Transplantation experiments were performed using either C57BL/6J mice or albino “Glowing Head” C57BL/6J mice (GH; C57BL/6-Tyr^c-Brd^ Tg(Gh1-luc/EGFP)D8Mrln/J)^38^. Alternatively, for experiments in immune compromised mice GH mice were crossed with Rag1^-/-^ mice.

For subcutaneous engraftment studies, 7-12 week-old mice were engrafted with a cell suspension of 150,000 KPAR1.3 cells, which were prepared in 100 μl of 50% PBS and 50% Geltrex (Gibco, A1413202). Tumour growth was measured twice weekly via calliper, and volume was calculated as [Length × Width²]/2. Mice were culled via cervical dislocation when tumours reached a 1.2 cm - 1.4 cm in diameter, or if any other humane endpoint was reached. Mice that rejected their primary subcutaneous tumour and remained tumour-free for ∼4 weeks post-treatment were classified as complete responders (CRs). These mice were re-challenged with 150,000 KPAR1.3 Luciferase-eGFP KRAS G12D cells injected into the opposite flank.

For in vivo imaging of Luciferase-eGFP-expressing KPAR1.3 KRAS G12D cells, mice were injected intraperitoneally with 100 μl 30 μg/ml XenoLight D-Luciferin (PerkinElmer, 122799), dissolved in PBS and anaesthetised via inhalation of 2% - 2.5% isoflurane, vaporised in O_2_. Bioluminescence was imaged in mice using the IVIS SpectrumCT scanner (PerkinElmer), and data was analysed with Living Image Software (versions 4.4–4.5, PerkinElmer).

The inhibitors adagrasib (MedchemExpress, 50 mg/kg), RMC-4998 (Revolution Medicines, 100 mg/kg), and RMC-4550 (Revolution Medicines, 30 mg/kg) were prepared in a 50 mM sodium citrate buffer (pH 5) containing 10% w/v Captisol (Ligand). Drugs were administered via oral gavage, once daily for a maximum of 6 days a week. Anti-PD-1 (BioXCell, clone RMP1-14, 10 mg/kg) was prepared in PBS and administered twice weekly via intraperitoneal injection. Combinations and durations of treatments are indicated in respective experiments.

### Flow cytometric techniques

Tumours were first minced with a scalpel, followed by enzymatic dissociation using mouse Tumour Dissociation Kit (Miltenyi Biotech, 130-096-730). The resulting cell suspension was filtered through a 70 μm MACS SmartStrainer (Miltenyi, 130-110-916), red blood cells were lysed with 5 ml ACK Lysing Buffer (Gibco, A1049201), followed by a washing step with FACS buffer (PBS containing 2% FBS, 1mM EDTA). For quantification of tumour cell subpopulations via their fluorescent protein (FP) expression, live cell suspensions were stained with viability dye DRAQ7 (1:100) and analysed on BD LSRFortessa™ X-20 Cell Analyzers (BD Biosciences) using corresponding BD FACSDiva™ and BD FlowJo analysis Software (version 10.8.1 - 10.8.3).

For immune phenotyping via spectral flow cytometry, additional single cell suspensions from spleens and tdLNs were obtained by directly pushing respective tissues through a 70 μm MACS SmartStrainers, followed by ACK lysis for spleen suspensions. All samples were then transferred to conical 96-well plates and normalised to ∼1.5*10^6^ cells per well. Before staining, cell suspensions were blocked with CD16/CD32 Mouse BD Fc Block (BD Biosciences, 553141) for 15 min. A staining panel master mix for extracellular staining was prepared (antibodies and viability dye listed in Table S5) and cells were stained for 20 min at RT. Single staining controls were prepared using UltraComp eBeads (Invitrogen, 01-2222-42). Cells were then fixed/permeabilised using Foxp3 Fixation/Permeabilization Buffer (Invitrogen, 00-5523-00) for 30 minutes at 4°C and washed with the buffer provided in the same kit. Samples were then stained with the intracellular antibody master mix (Table S5) for 20 minutes at RT, washed and resuspended in FACS buffer. Immune-phenotyping was performed on the Cytek Aurora spectral flow cytometer, with spectral unmixing using Cytek SpectroFlow Software and further analysis in FlowJo (versions 10.8.1 – 10.10.1).

For the analysis of Emv2-specific T cells, initial sample processing was performed as described above. Once samples were transferred to 96 well plate, cells were first incubated in PBS containing CD16/CD32 Mouse BD Fc Block (BD Biosciences, 553141) and Zombie NIR viability dye for 15 minutes at RT. Next, samples were stained with T-select H-2Kb MuLV p15E Tetramer-KSPWFTTL (2x) for 40 min at 4°C. Subsequently, extracellular master mix, prepared with the antibodies listed in Table S6 diluted in Foxp3 Fixation/Permeabilization Buffer (Invitrogen, 00-5523-00), was added and samples were incubated for 30 min at 4°C. Fixation, intracellular staining and sample immune-phenotyping on Cytek Aurora spectral flow cytometer was performed as described above.

### Quantification of tumour-binding antibodies

To assess whether serum from tumour-bearing mice contained tumour cell-opsonising antibodies, serum samples were collected from mice harbouring KPAR1.3 KRAS G12C tumours, from mice with KPAR1.3 KRAS G12D Emv2 KO tumours, and from naïve mice without tumours. To test for the presence of tumour cell–specific antibodies, 3 × 10⁵ KPAR1.3 cells were plated per well in a 96-well plate for each isogenic version of the cell line: (1) KRAS G12C (2) KRAS G12D, and (3) KRAS G12D Emv2 KO. Samples were incubated with 3 µl of mouse serum, diluted in 100 µl FACS buffer per well for 1 hour at 4 °C. Cells were then stained with FITC-conjugated anti-mouse IgG antibody (BioLegend, Cat. No. 406001) for 30 min at 4 °C. DAPI was added at working concentration to exclude dead cells. Samples were acquired on a BD LSRFortessa™ X-20 Cell Analyzer (BD Biosciences), and data was analysed using FlowJo (v10). The median fluorescence intensity (MFI) of FITC was used as a measure of antibody binding. For each condition, FITC MFI values were normalised to those obtained from serum of naïve mice.

### Bulk RNA sequencing

Subcutaneous tumours were extracted, dissociated into single-cell suspensions, and red blood cells were lysed using ACK lysis buffer (Gibco, A1049201). To eliminate additional auto-fluorescent events in the eBFP+ sort gate, CD45+ immune cells were removed priorly via magnetic pulldown, using an a-CD45 microbead kit (Miltenyi Biotec, 130-052-301). Cancer cells were then flow cytometrically sorted via their expressed FPs and RNA was purified using the RNeasy Micro Kit (Qiagen, 74004). Alternatively, for in vitro characterization of the KPAR.E2 cell line, parental KPAR1.3 G12C and KPAR.E2 cells were treated for 24 hours with DMSO or 100 nM adagrasib and RNA was extracted using the RNAeasy Mini Kit (Qiagen) following manufacturer’s instructions. The Genomics Science Technology Platform (STP) at the Francis Crick Institute prepared reverse stranded sequencing libraries from 50 ng of RNA using either NEBNext Ultra II Directional PolyA mRNA (New England Biolabs) or the Watchmaker mRNA Library Prep Kit (modules K0105 and K0078), following the manufacturer’s protocols. Paired-end sequencing (100 bp) was performed on an Illumina NovaSeq X platform, with a minimum depth of 25 million reads per sample.

Sequencing results were first pre-processed utilising the nf-core pipeline (v3.14.0), selecting the *star_rsem* aligner and the mouse reference genome version *Mus_musculus.GRCm38.95 for alignment of sequenced reads.* Data analysis was conducted using the DESeq2 pipeline, and the hallmark gene set collection from the MSigDB website (https://www.gsea-msigdb.org/gsea/msigdb) was used for GSEA analysis.

### Multiplex Immunohistochemistry

Stainings of Formalin-Fixed Paraffin-Embedded (FFPE) tissue sections were performed by the Experimental Histopathology (EHP) core facility at the Francis Crick Institute. Tumours were fixed in 10 % neutral buffered formalin (NBF) for 48h, stored in 70% EtOH, and embedded in paraffin wax using a Tissue-Tek VIP® 6 AI processor (Sakura). 3 μm sections were cut and mounted on glass slides, which were baked at 60°C for 1 hour to adhere the tissue and melt the wax. Slides were then transferred to a Leica Bond Rx platform (Leica, 3498240), where a 0.1% BSA solution for protein blocking and a 3% hydrogen peroxide solution for inhibiting endogenous peroxidases were applied to prepare the slides for staining. Antigens were then sequentially stained using HRP-conjugated antibodies and various Opal dyes (Akoya). The antibodies targeting the following epitopes were used: eGFP/eBFP, clone A-6455, ThermoFisher; CD4, clone EPR19514, Abcam; Caspase3, clone D3E9, Cell Signaling; CD8a, clone EPR21769, Abcam; Luciferase, NB100-1677, Novus Biologicals; Ki67, ab15580, Abcam. Between each antigen staining, slides underwent antibody stripping and antigen retrieval by heating at 95°C for 20 minutes with Epitope Retrieval Solutions (Leica, AR9961 and AR9640). Nuclei were then stained with DAPI (Thermo Scientific, 62248) and coverslips mounted using ProLong Gold Antifade reagent (Invitrogen, P36934). Finally, slides were scanned on a PhenoImager HT (Akoya, VP2143N2122) in MOTiF scanning mode using Vectra Polaris software (version 1.0.13) and unmixing of fluorescent spectra was performed via appurtenant InForm software (version 2.6.0).

Using QuPath (version0.4.3), tumour areas for analysis were first outlined to avoid edge effects. Larger necrotic regions were identified based on Casp3 expression, in consultation with experimental histopathology, and were also excluded from analysis. Additional technical artefacts such as wrinkles, rips, or tissue folds were similarly annotated and excluded from downstream processing. Because stain intensity varied across sections, a mean background subtraction to balance uneven illumination was performed. A gaussian blur was also performed to smoothen out features and improve downstream analysis. Nuclei were segmented with StarDist^50–52^, a U-Net–based approach trained and benchmarked on our dataset. In parallel, semantic masks for CD4, CD8, GFP/BFP, luciferase, Ki67, and Casp3 were generated using a trained random-forest classifier (DOI 10.5281/zenodo.3555620). Cell types were assigned based on the intersection between nuclear masks with the corresponding marker masks (CD4, CD8, GFP/BFP, luciferase). Conflicts between markers were resolved using pre-specified thresholds. Cytoplasm was delineated without a membrane marker under three constraints. (i) Cytoplasm surrounds its nucleus and is spatially bounded; we created a mask based on the position of each positive nucleus and expanded it into their corresponding, overlapping marker masks. (ii) Cytoplasm from different cells does not overlap; overlaps were resolved by Euclidean Voronoi reassignment of pixels to the nearest cell centre of mass, with nuclei reinforced as true positives. (iii) Cytoplasm is contiguous; after Voronoi assignment, fragmented pixel islands were reassigned to the adjacent cell with the greatest connected support. Isolated pixels with no neighbouring cells were discarded.

### Statistical analysis

Data were compared using pair-end Student’s t-test or analysis of variance (ANOVA) if more than two experimental groups were analysed. Test used for each experiment is indicated in the figure legends. Significance was determined at p<0.05 (*p<0.05, **p<0.01, ***p < 0.001, ****p<0.0001).

## DATA AVAILABILITY

The RNA-seq data for this study has been deposited in the European Nucleotide Archive (ENA) at EMBL-EBI under the accession numbers PREB101476, PRJEB101500 and PRJEB101588. The remaining data are available within the article, Supplementary Information or Source Data provided with this paper.

## ACKNOWLEDGEMENTS

We thank the core facilities at the Francis Crick Institute including the Biological Research Facility, Scientific Computing, Flow Cytometry, Experimental Histopathology, Cell Science, and In Vivo Imaging Science Technology Platforms (STP). We acknowledge the Genomics STP, and particularly Deb Jackson, Daniel Leonce and Marg Crawford, for their contributions to mRNA library preparation and sequencing. We acknowledge the Bioinformatics and Biostatistics STP, and particularly Probir Chakravarty and Gavin Kelly for their analysis support. We thank the members of the Oncogene Biology Laboratory for their discussions and critical reading of the manuscript. We thank Elsa Quintana, Cristina Blaj and Jan Smith at Revolution Medicines for their advice and support.

This work was supported by the Francis Crick Institute which receives its core funding from Cancer Research UK (CC2097 and CC2119), the UK Medical Research Council (CC2097 and CC2119) and the Wellcome Trust (CC2097 and CC2119). This work also received funding from the European Research Council Advanced Grant RASImmune and from Revolution Medicines, Inc. under a collaborative research agreement.

## AUTHOR CONTRIBUTIONS

M.T., K.B.N., M.M.-A. and J.D. designed the study, interpreted the results and wrote the manuscript. M.T., C.P., P.A., S.R., C.P., A.dC., A.A.V., S.C.T., A.M. and S.M. performed the biochemical experiments. M.T. and C.M. performed in vitro studies. M.T., K.B.N., J.C. performed bioinformatic and computational analyses. J.D. and N.G. contributed with interpretation and resources. All authors contributed to the manuscript revision and review.

## COMPETING INTERESTS

J.D. has acted as a consultant for AstraZeneca, Jubilant, Theras, Roche and Vividion and has received research funding from Bristol Myers Squibb, Revolution Medicines, Vividion, AstraZeneca and Novartis. S.C.T. has acted as a consultant for Revolution Medicines. The other authors declare that they have no competing interests.

## REFERENCES

1. Prior, I.A., Hood, F.E., and Hartley, J.L. (2020). The Frequency of Ras Mutations in Cancer. Cancer Res 80, 2969–2974. 10.1158/0008-5472.CAN-19-3682.

2. Downward, J. (2003). Targeting RAS signalling pathways in cancer therapy. Nat Rev Cancer 3, 11–22. 10.1038/nrc969.

3. Hamarsheh, S., Groß, O., Brummer, T., and Zeiser, R. (2020). Immune modulatory effects of oncogenic KRAS in cancer. Nat Commun 11, 5439. 10.1038/s41467-020-19288-6.

4. Molina-Arcas, M., and Downward, J. (2024). Exploiting the therapeutic implications of KRAS inhibition on tumor immunity. Cancer Cell 42, 338–357. 10.1016/j.ccell.2024.02.012.

5. Liao, W., Overman, M.J., Boutin, A.T., Shang, X., Zhao, D., Dey, P., Li, J., Wang, G., Lan, Z., Li, J., et al. (2019). KRAS-IRF2 Axis Drives Immune Suppression and Immune Therapy Resistance in Colorectal Cancer. Cancer Cell 35, 559–572.e557. 10.1016/j.ccell.2019.02.008.

6. Mugarza, E., van Maldegem, F., Boumelha, J., Moore, C., Rana, S., Llorian Sopena, M., East, P., Ambler, R., Anastasiou, P., Romero Clavijo, P., et al. (2022). Therapeutic KRASG12C inhibition drives effective interferon-mediated anti-tumour immunity in immunogenic lung cancers. Science Advances 8, eabm8780.

7. Coelho, M.A., de Carne Trecesson, S., Rana, S., Zecchin, D., Moore, C., Molina-Arcas, M., East, P., Spencer-Dene, B., Nye, E., Barnouin, K., et al. (2017). Oncogenic RAS Signaling Promotes Tumor Immunoresistance by Stabilizing PD-L1 mRNA. Immunity 47, 1083–1099 e1086. 10.1016/j.immuni.2017.11.016.

8. Pylayeva-Gupta, Y., Lee, K.E., Hajdu, C.H., Miller, G., and Bar-Sagi, D. (2012). Oncogenic Kras-induced GM-CSF production promotes the development of pancreatic neoplasia. Cancer Cell 21, 836–847. 10.1016/j.ccr.2012.04.024.

9. Boumelha, J., de Castro, A., Bah, N., Cha, H., de Carné Trécesson, S., Rana, S., Tomaschko, M., Anastasiou, P., Mugarza, E., Moore, C., et al. (2024). CRISPR-Cas9 Screening Identifies KRAS-Induced COX2 as a Driver of Immunotherapy Resistance in Lung Cancer. Cancer Res 84, 2231–2246. 10.1158/0008-5472.Can-23-2627.

10. Ostrem, J.M., Peters, U., Sos, M.L., Wells, J.A., and Shokat, K.M. (2013). K-Ras(G12C) inhibitors allosterically control GTP affinity and effector interactions. Nature 503, 548–551. 10.1038/nature12796.

11. Moore, A.R., Rosenberg, S.C., McCormick, F., and Malek, S. (2020). RAS-targeted therapies: is the undruggable drugged? Nat Rev Drug Discov 19, 533–552. 10.1038/s41573-020-0068-6.

12. Canon, J., Rex, K., Saiki, A.Y., Mohr, C., Cooke, K., Bagal, D., Gaida, K., Holt, T., Knutson, C.G., Koppada, N., et al. (2019). The clinical KRAS(G12C) inhibitor AMG 510 drives anti-tumour immunity. Nature 575, 217–223. 10.1038/s41586-019-1694-1.

13. Hallin, J., Engstrom, L.D., Hargis, L., Calinisan, A., Aranda, R., Briere, D.M., Sudhakar, N., Bowcut, V., Baer, B.R., Ballard, J.A., et al. (2020). The KRAS(G12C) Inhibitor MRTX849 Provides Insight toward Therapeutic Susceptibility of KRAS-Mutant Cancers in Mouse Models and Patients. Cancer Discov 10, 54–71. 10.1158/2159-8290.Cd-19-1167.

14. Briere, D.M., Li, S., Calinisan, A., Sudhakar, N., Aranda, R., Hargis, L., Peng, D.H., Deng, J., Engstrom, L.D., Hallin, J., et al. (2021). The KRAS(G12C) Inhibitor MRTX849 Reconditions the Tumor Immune Microenvironment and Sensitizes Tumors to Checkpoint Inhibitor Therapy. Mol Cancer Ther 20, 975–985. 10.1158/1535-7163.MCT-20-0462.

15. Hu, H., Cheng, R., Wang, Y., Wang, X., Wu, J., Kong, Y., Zhan, S., Zhou, Z., Zhu, H., Yu, R., et al. (2023). Oncogenic KRAS signaling drives evasion of innate immune surveillance in lung adenocarcinoma by activating CD47. J Clin Invest 133. 10.1172/jci153470.

16. van Maldegem, F., Valand, K., Cole, M., Patel, H., Angelova, M., Rana, S., Colliver, E., Enfield, K., Bah, N., Kelly, G., et al. (2021). Characterisation of tumour microenvironment remodelling following oncogene inhibition in preclinical studies with imaging mass cytometry. Nat Commun 12, 5906. 10.1038/s41467-021-26214-x.

17. Ng, K.W., Boumelha, J., Enfield, K.S.S., Almagro, J., Cha, H., Pich, O., Karasaki, T., Moore, D.A., Salgado, R., Sivakumar, M., et al. (2023). Antibodies against endogenous retroviruses promote lung cancer immunotherapy. Nature 616, 563–573. 10.1038/s41586-023-05771-9.

18. Boumelha, J., de Carne Trecesson, S., Law, E.K., Romero-Clavijo, P., Coelho, M., Ng, K., Mugarza, E., Moore, C., Rana, S., Caswell, D.R., et al. (2022). An immunogenic model of KRAS-mutant lung cancer enables evaluation of targeted therapy and immunotherapy combinations. Cancer Res 82, 3435–3448. 10.1158/0008-5472.CAN-22-0325.

19. Janne, P.A., Riely, G.J., Gadgeel, S.M., Heist, R.S., Ou, S.I., Pacheco, J.M., Johnson, M.L., Sabari, J.K., Leventakos, K., Yau, E., et al. (2022). Adagrasib in Non-Small-Cell Lung Cancer Harboring a KRAS(G12C) Mutation. N Engl J Med 387, 120–131. 10.1056/NEJMoa2204619.

20. Skoulidis, F., Li, B.T., Dy, G.K., Price, T.J., Falchook, G.S., Wolf, J., Italiano, A., Schuler, M., Borghaei, H., Barlesi, F., et al. (2021). Sotorasib for Lung Cancers with KRAS p.G12C Mutation. N Engl J Med 384, 2371–2381. 10.1056/NEJMoa2103695.

21. de Langen, A.J., Johnson, M.L., Mazieres, J., Dingemans, A.C., Mountzios, G., Pless, M., Wolf, J., Schuler, M., Lena, H., Skoulidis, F., et al. (2023). Sotorasib versus docetaxel for previously treated non-small-cell lung cancer with KRAS(G12C) mutation: a randomised, open-label, phase 3 trial. Lancet 401, 733–746. 10.1016/s0140-6736(23)00221-0.

22. Schulze, C.J., Seamon, K.J., Zhao, Y., Yang, Y.C., Cregg, J., Kim, D., Tomlinson, A., Choy, T.J., Wang, Z., Sang, B., et al. (2023). Chemical remodeling of a cellular chaperone to target the active state of mutant KRAS. Science 381, 794–799. 10.1126/science.adg9652.

23. Jänne, P.A., Bigot, F., Papadopoulos, K., Eberst, L., Sommerhalder, D., Lebellec, L., Voon, P.J., Pellini, B., Kalinka, E., and Arbour, K. (2023). Abstract PR014: Preliminary safety and anti-tumor activity of RMC-6291, a first-in-class, tri-complex KRASG12C (ON) inhibitor, in patients with or without prior KRASG12C (OFF) inhibitor treatment. Molecular Cancer Therapeutics 22, PR014–PR014.

24. Awad, M.M., Liu, S., Rybkin, II, Arbour, K.C., Dilly, J., Zhu, V.W., Johnson, M.L., Heist, R.S., Patil, T., Riely, G.J., et al. (2021). Acquired Resistance to KRAS(G12C) Inhibition in Cancer. N Engl J Med 384, 2382–2393. 10.1056/NEJMoa2105281.

25. Riedl, J.M., Fece de la Cruz, F., Lin, J.J., Parseghian, C., Kim, J.E., Matsubara, H., Barnes, H., Caughey, B., Norden, B.L., Morales-Giron, A.A., et al. (2025). Genomic landscape of clinically acquired resistance alterations in patients treated with KRAS(G12C) inhibitors. Ann Oncol 36, 682–692. 10.1016/j.annonc.2025.01.020.

26. Xue, J.Y., Zhao, Y., Aronowitz, J., Mai, T.T., Vides, A., Qeriqi, B., Kim, D., Li, C., de Stanchina, E., Mazutis, L., et al. (2020). Rapid non-uniform adaptation to conformation-specific KRAS(G12C) inhibition. Nature 577, 421–425. 10.1038/s41586-019-1884-x.

27. Ryan, M.B., Fece de la Cruz, F., Phat, S., Myers, D.T., Wong, E., Shahzade, H.A., Hong, C.B., and Corcoran, R.B. (2020). Vertical Pathway Inhibition Overcomes Adaptive Feedback Resistance to KRAS(G12C) Inhibition. Clin Cancer Res 26, 1633– 1643. 10.1158/1078-0432.Ccr-19-3523.

28. Adachi, Y., Ito, K., Hayashi, Y., Kimura, R., Tan, T.Z., Yamaguchi, R., and Ebi, H. (2020). Epithelial-to-Mesenchymal Transition is a Cause of Both Intrinsic and Acquired Resistance to KRAS G12C Inhibitor in KRAS G12C-Mutant Non-Small Cell Lung Cancer. Clin Cancer Res 26, 5962–5973. 10.1158/1078-0432.Ccr-20-2077.

29. Hagenbeek, T.J., Zbieg, J.R., Hafner, M., Mroue, R., Lacap, J.A., Sodir, N.M., Noland, C.L., Afghani, S., Kishore, A., Bhat, K.P., et al. (2023). An allosteric pan-TEAD inhibitor blocks oncogenic YAP/TAZ signaling and overcomes KRAS G12C inhibitor resistance. Nat Cancer 4, 812–828. 10.1038/s43018-023-00577-0.

30. Tanaka, N., and Ebi, H. (2025). Mechanisms of Resistance to KRAS Inhibitors: Cancer Cells’ Strategic Use of Normal Cellular Mechanisms to Adapt. Cancer Sci 116, 600– 612. 10.1111/cas.16441.

31. Ebright, R.Y., Dilly, J., Shaw, A.T., and Aguirre, A.J. (2025). Response and Resistance to RAS Inhibition in Cancer. Cancer Discov 15, 1325–1349. 10.1158/2159-8290.Cd-25-0349.

32. Onoi, K., Chihara, Y., Uchino, J., Shimamoto, T., Morimoto, Y., Iwasaku, M., Kaneko, Y., Yamada, T., and Takayama, K. (2020). Immune Checkpoint Inhibitors for Lung Cancer Treatment: A Review. J Clin Med 9. 10.3390/jcm9051362.

33. Musaelyan, A.A., Moiseyenko, F.V., Emileva, T.E., Oganesyan, A.P., Oganyan, K.A., Urtenova, M.A., Odintsova, S.V., Chistyakov, I.V., Degtyarev, A.M., Akopov, A.L., et al. (2024). Clinical predictors of response to single-agent immune checkpoint inhibitors in chemotherapy-pretreated non-small cell lung cancer. Mol Clin Oncol 20, 32. 10.3892/mco.2024.2730.

34. Anastasiou, P., Moore, C., Rana, S., Tomaschko, M., Pillsbury, C.E., de Castro, A., Boumelha, J., Mugarza, E., de Carné Trécesson, S., Mikolajczak, A., et al. (2024). Combining RAS(ON) G12C-selective inhibitor with SHP2 inhibition sensitises lung tumours to immune checkpoint blockade. Nat Commun 15, 8146. 10.1038/s41467-024-52324-3.

35. Quintana, E., Schulze, C.J., Myers, D.R., Choy, T.J., Mordec, K., Wildes, D., Shifrin, N.T., Belwafa, A., Koltun, E.S., Gill, A.L., et al. (2020). Allosteric Inhibition of SHP2 Stimulates Antitumor Immunity by Transforming the Immunosuppressive Environment. Cancer Res 80, 2889–2902. 10.1158/0008-5472.CAN-19-3038.

36. Fedele, C., Li, S., Teng, K.W., Foster, C.J.R., Peng, D., Ran, H., Mita, P., Geer, M.J., Hattori, T., Koide, A., et al. (2021). SHP2 inhibition diminishes KRASG12C cycling and promotes tumor microenvironment remodeling. J Exp Med 218. 10.1084/jem.20201414.

37. Li, B.T., Falchook, G.S., Durm, G.A., Burns, T.F., Skoulidis, F., Ramalingam, S.S., Spira, A., Bestvina, C.M., Goldberg, S.B., Veluswamy, R., et al. (2022). OA03.06 CodeBreaK 100/101: First Report of Safety/Efficacy of Sotorasib in Combination with Pembrolizumab or Atezolizumab in Advanced KRAS p.G12C NSCLC. Journal of Thoracic Oncology 17, S10–S11. 10.1016/j.jtho.2022.07.025.

38. Day, C.P., Carter, J., Weaver Ohler, Z., Bonomi, C., El Meskini, R., Martin, P., Graff-Cherry, C., Feigenbaum, L., Tuting, T., Van Dyke, T., et al. (2014). “Glowing head” mice: a genetic tool enabling reliable preclinical image-based evaluation of cancers in immunocompetent allografts. PLoS One 9, e109956. 10.1371/journal.pone.0109956.

39. Haigis, K.M. (2017). KRAS Alleles: The Devil Is in the Detail. Trends in Cancer 3, 686– 697. 10.1016/j.trecan.2017.08.006.

40. Torrejon, D.Y., Abril-Rodriguez, G., Champhekar, A.S., Tsoi, J., Campbell, K.M., Kalbasi, A., Parisi, G., Zaretsky, J.M., Garcia-Diaz, A., Puig-Saus, C., et al. (2020). Overcoming Genetically Based Resistance Mechanisms to PD-1 Blockade. Cancer Discov 10, 1140–1157. 10.1158/2159-8290.Cd-19-1409.

41. Cook, J., and Hagemann, T. (2013). Tumour-associated macrophages and cancer. Current Opinion in Pharmacology 13, 595–601. 10.1016/j.coph.2013.05.017.

42. Singh, A., Greninger, P., Rhodes, D., Koopman, L., Violette, S., Bardeesy, N., and Settleman, J. (2009). A gene expression signature associated with “K-Ras addiction” reveals regulators of EMT and tumor cell survival. Cancer Cell 15, 489–500. 10.1016/j.ccr.2009.03.022.

43. Dilly, J., Hoffman, M.T., Abbassi, L., Li, Z., Paradiso, F., Parent, B.D., Hennessey, C.J., Jordan, A.C., Morgado, M., Dasgupta, S., et al. (2024). Mechanisms of Resistance to Oncogenic KRAS Inhibition in Pancreatic Cancer. Cancer Discov 14, 2135–2161. 10.1158/2159-8290.Cd-24-0177.

44. Kitai, H., Ebi, H., Tomida, S., Floros, K.V., Kotani, H., Adachi, Y., Oizumi, S., Nishimura, M., Faber, A.C., and Yano, S. (2016). Epithelial-to-Mesenchymal Transition Defines Feedback Activation of Receptor Tyrosine Kinase Signaling Induced by MEK Inhibition in KRAS-Mutant Lung Cancer. Cancer Discov 6, 754–769. 10.1158/2159-8290.Cd-15-1377.

45. Solanki, H.S., Welsh, E.A., Fang, B., Izumi, V., Darville, L., Stone, B., Franzese, R., Chavan, S., Kinose, F., Imbody, D., et al. (2021). Cell Type-specific Adaptive Signaling Responses to KRAS(G12C) Inhibition. Clin Cancer Res 27, 2533–2548. 10.1158/1078-0432.Ccr-20-3872.

46. Obenauf, A.C., Zou, Y., Ji, A.L., Vanharanta, S., Shu, W., Shi, H., Kong, X., Bosenberg, M.C., Wiesner, T., Rosen, N., et al. (2015). Therapy-induced tumour secretomes promote resistance and tumour progression. Nature 520, 368–372. 10.1038/nature14336.

47. Chour, A., Denis, J., Mascaux, C., Zysman, M., Bigay-Game, L., Swalduz, A., Gounant, V., Cortot, A., Darrason, M., Fallet, V., et al. (2023). Brief Report: Severe Sotorasib-Related Hepatotoxicity and Non-Liver Adverse Events Associated With Sequential Anti–Programmed Cell Death (Ligand)1 and Sotorasib Therapy in KRASG12C-Mutant Lung Cancer. Journal of Thoracic Oncology 18, 1408–1415. 10.1016/j.jtho.2023.05.013.

48. Cole, M., Anastasiou, P., Lee, C., Yu, X., de Castro, A., Roelink, J., Moore, C., Mugarza, E., Jones, M., Valand, K., et al. (2024). Spatial multiplex analysis of lung cancer reveals that regulatory T cells attenuate KRAS-G12C inhibitor-induced immune responses. Sci Adv 10, eadl6464. 10.1126/sciadv.adl6464.

49. Liu, Y., Han, J., Hsu, W.H., LaBella, K.A., Deng, P., Shang, X., Tallón de Lara, P., Cai, L., Jiang, S., and DePinho, R.A. (2025). Combined KRAS Inhibition and Immune Therapy Generates Durable Complete Responses in an Autochthonous PDAC Model. Cancer Discov 15, 162–178. 10.1158/2159-8290.Cd-24-0489.

50. Schmidt, U., Weigert, M., Broaddus, C., and Myers, G. (2018). Cell Detection with Star-Convex Polygons. held in Cham, 2018//. A.F. Frangi, J.A. Schnabel, C. Davatzikos, C. Alberola-López, and G. Fichtinger, eds. (Springer International Publishing), pp. 265–273.

51. Weigert, M., Schmidt, U., Haase, R., Sugawara, K., and Myers, G. (2020). Star-convex Polyhedra for 3D Object Detection and Segmentation in Microscopy. 1–5 March 2020. pp. 3655–3662.

52. Weigert, M., and Schmidt, U. (2022). Nuclei Instance Segmentation and Classification in Histopathology Images with Stardist. 28–31 March 2022. pp. 1–4.

