## Supplemental Figures S1-13, Supplemental Tables S1-6 for "Leveraging death of drug-sensitive cancer cells to promote immune-mediated bystander killing of subclones of drug-resistant tumour cells"

### Supplementary Material

Tomaschko et al.

27 October 2025

#### Supplementary Figures

##### Supplementary Figure 1

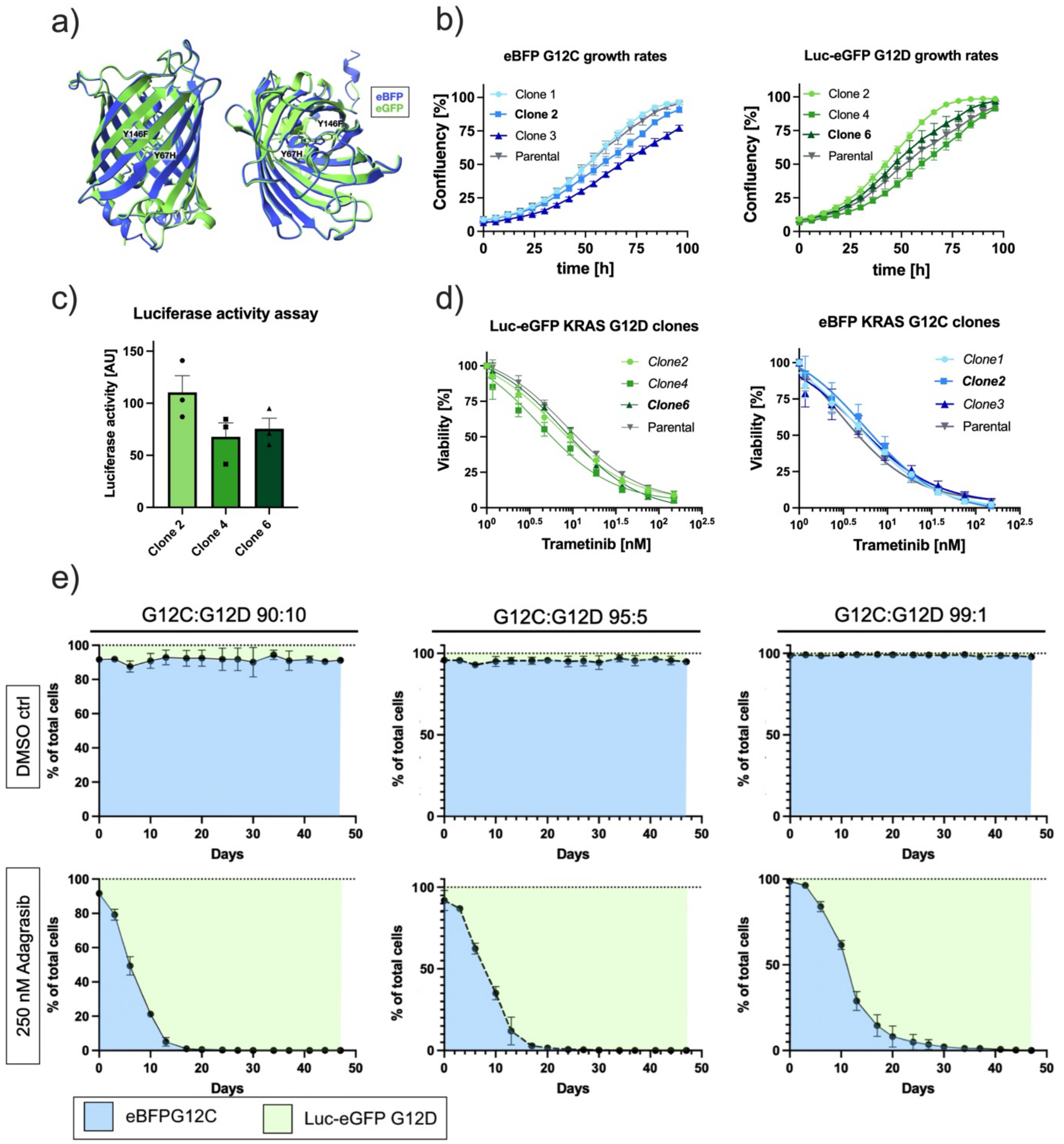

**Supplementary Figure 1. Generation of reporter-transduced KPAR1.3 cell lines**

a) Prediction of structural similarity between eBFP and eGFP. eBFP protein structure is based on AlphaFold prediction, eGFP protein based on previously solved protein crystal structure (PDB: 4EUL), alignment based on the Needleman-Wunsch algorithm.

b) Representative plots of selected Luc-eGFP KRAS G12D (left) or eBFP KRAS G12C (right) subclones, showing proliferation rates over time assessed via IncuCyte live imaging. n=3, mean ± SEM.

c) Luciferase activity of KPAR1.3 KRAS G12D Luc-eGFP sub-clonal cell lines. Data is normalised to parental KPAR1.3 cells. n=3, tr(n) = 3, mean ± SEM.

d) Viability assays of selected subclones compared to parental counterparts. Cells were treated with the MEK inhibitor trametinib for 72 hours. n = 3, tr(n) = 3, mean ± SEM.

e) Co-culture assay assessing adagrasib response over time. Mixed cultures of eBFP KRAS G12C and Luc-eGFP KRAS G12D cells were continuously treated with DMSO or 250 nM adagrasib over time and fraction of G12C and G12D cells was measured by FACS at indicated time points. Blue and green areas on graphs depict the fractions of KPAR1.3 KRAS G12C or G12D populations, respectively. n = 3, tr(n) = 3, mean ± SD.

### Supplementary Figure 2

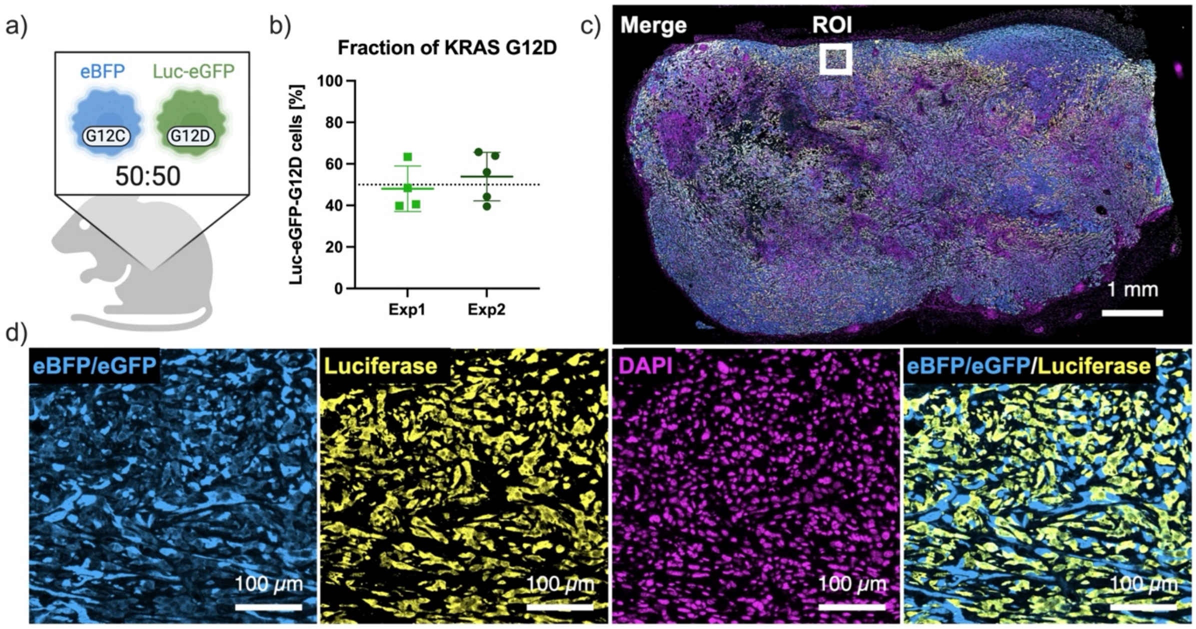


**Supplementary Figure 2. Characterisation of mixed KPAR 1.3 tumours**
a) Schematic of experiment. Luc-eGFP KRAS G12D and eBFP KRAS G12C cells were co-engrafted subcutaneously in GH-mice at a 50:50 ratio to assess the growth dynamics of co-engrafted subclones in vivo.
b) Fraction of Luc-eGFP KRAS G12D cells within all live cancer cells of two independent experiments. Flow cytometric analysis was performed 14 days after engraftment of tumours. Dots represent individual tumours, with one subcutaneous tumour per mouse; mean ± SD.
c-d) Representative immunohistochemistry staining of one tumour 14 days after co-engraftment. c) Full tumour overview (scale bar 1mm) and d) Representative ROIs of individual channels (scale bar 100 µm). eBFP/eGFP: blue (G12C and G12D cells); luciferase: yellow (G12D cells); DAPI: magenta (nucleus).

### Supplementary Figure 3

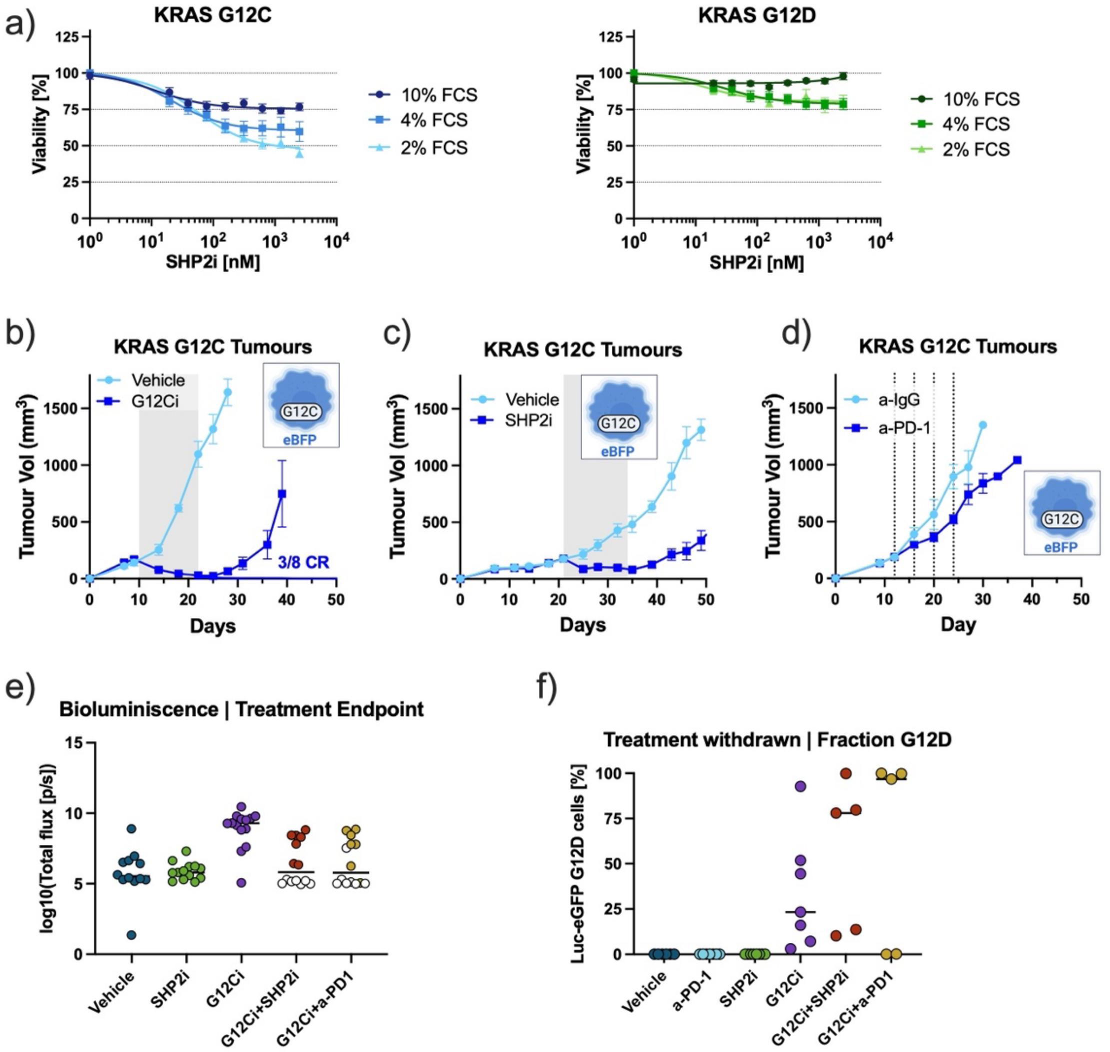


**Supplementary Figure 3. Combination strategies to enhance treatment responses**
a) Viability of eBFP KRAS G12C (left) and Luc-eGFP KRAS G12D (right) cells treated for 72 hours with serial dilutions of RMC-4550 (SHP2i). Cells were cultured with the indicated percentages of FCS. n=3-4, tr(n)=3, mean ± SEM.
b) eBFP KRAS G12C subcutaneous tumours were treated daily for two weeks (grey area) with vehicle (n=6) or the RAS G12C(ON) inhibitor RMC-4998 (100mg/kg; G12Ci; n=8). Number of complete responders (CR) is indicated. Mean ± SEM.
c) eBFP KRAS G12C subcutaneous tumours were treated daily for two weeks (grey area) with vehicle (n=8) or the SHP2 inhibitor RMC-4550 (30mg/kg; SHP2i; n=8). Mean ± SEM.
d) eBFP KRAS G12C subcutaneous tumours were treated with four doses (dotted lines) of isotype control (a-IgG, n=7) or a-PD-1 (10 mg/kg, n=8). Mean ± SEM.
e) Log10-transformed data of bioluminescence scans from mice in Figure 2e at treatment endpoint. Each dot represents one tumour; means are indicated by black lines; CR mice are highlighted in white across treatment groups.
f) Endpoint tumours from Figure 2c were isolated and fraction of Luc-eGFP KRAS G12D cells over all cancer cells was determined via flow cytometry; each dot represents one tumour, mean ± SEM.

### Supplementary Figure 4

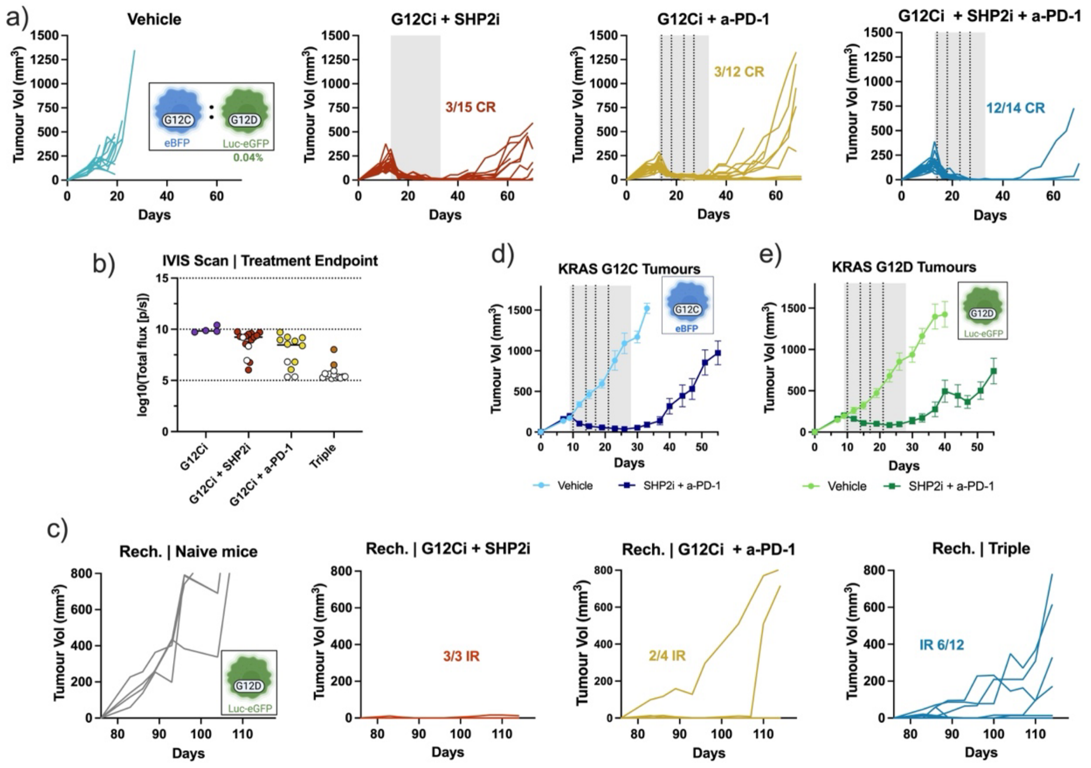


**Supplementary Figure 4. Triple combination increases the generation of complete responders**

a) Tumour growth of mixed subcutaneous tumours engrafted with BFP KRAS G12C cells plus a fraction of 0.04% Luc-eGFP KRAS G12D cells. Mice were treated for three weeks with the RAS G12C(ON) inhibitor RMC-4998 (100 mg/kg; G12Ci) with or without the SHP2 inhibitor RMC-4550 (30 mg/kg, SHP2i) and/or a-PD-1 (10 mg/kg). RMC-4998 and RMC-4550 were dosed by oral gavage daily (grey area) and a-PD1 was administered i.p. twice a week (vertical dotted lines). Complete responders (CR) are indicated.

b) Log10-transformed data of bioluminescence scans of mice in panel (a) at treatment endpoint by treatment groups. Each dot represents one tumour. Means are indicated by black vertical lines; CR mice are highlighted in white across treatment groups.

c) Complete responders in (a) were injected in the opposite flank with Luc-eGFP KRAS G12D cells. Mice that immune rejected (IR) the re-injected cancer cells are indicated. Graph titles indicate the treatment that the primary tumour received.

d-e) eBFP KRAS G12C (e) or Luc-eGFP KRAS G12D (f) subcutaneous tumours were treated for three weeks with either vehicle or a combination of RMC-4550 (30mg/kg; SHP2i) plus a-PD-1 (10mg/kg). N=12-13. Mean ± SEM.

### Supplementary Figure 5

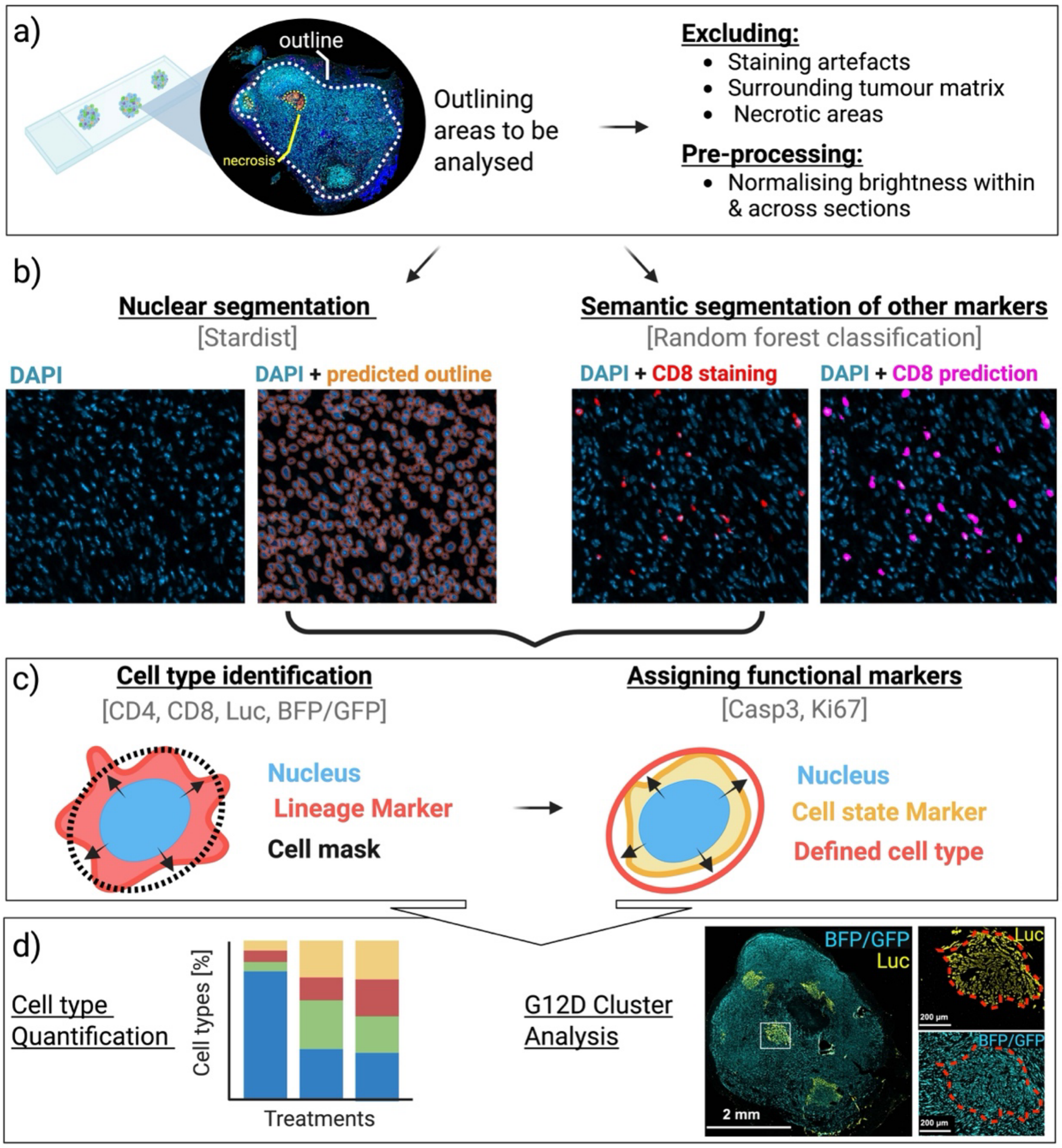


**Supplementary Figure 5. Outline of image analysis pipeline**
a) Objective slide containing several tumour sections was imaged in one frame, ROIs within image were outlined (white outline) excluding staining artefacts and the matrix surrounding the tumour; necrotic areas were also demarcated as separate entities (yellow outlines); these areas were excluded for subsequent analysis.
b) Left panel: exemplary ROI showing nuclear segmentation, right panel: exemplary ROI showing semantic segmentation of $CD8^+$ T cells.
c) Schematic of subsequent steps. Left panel: cell type identification using the lineage markers of BFP/GBP and Luciferase for cancer cell as well as CD8 and CD4 to demarcate T cells. Right panel: Subsequently, defined cell types are assigned positive or negative values for the functional state markers Ki67 and Casp3.
d) After cell type classification, respective populations can be quantified.
Figure was created with BioRender.

### Supplementary Figure 6

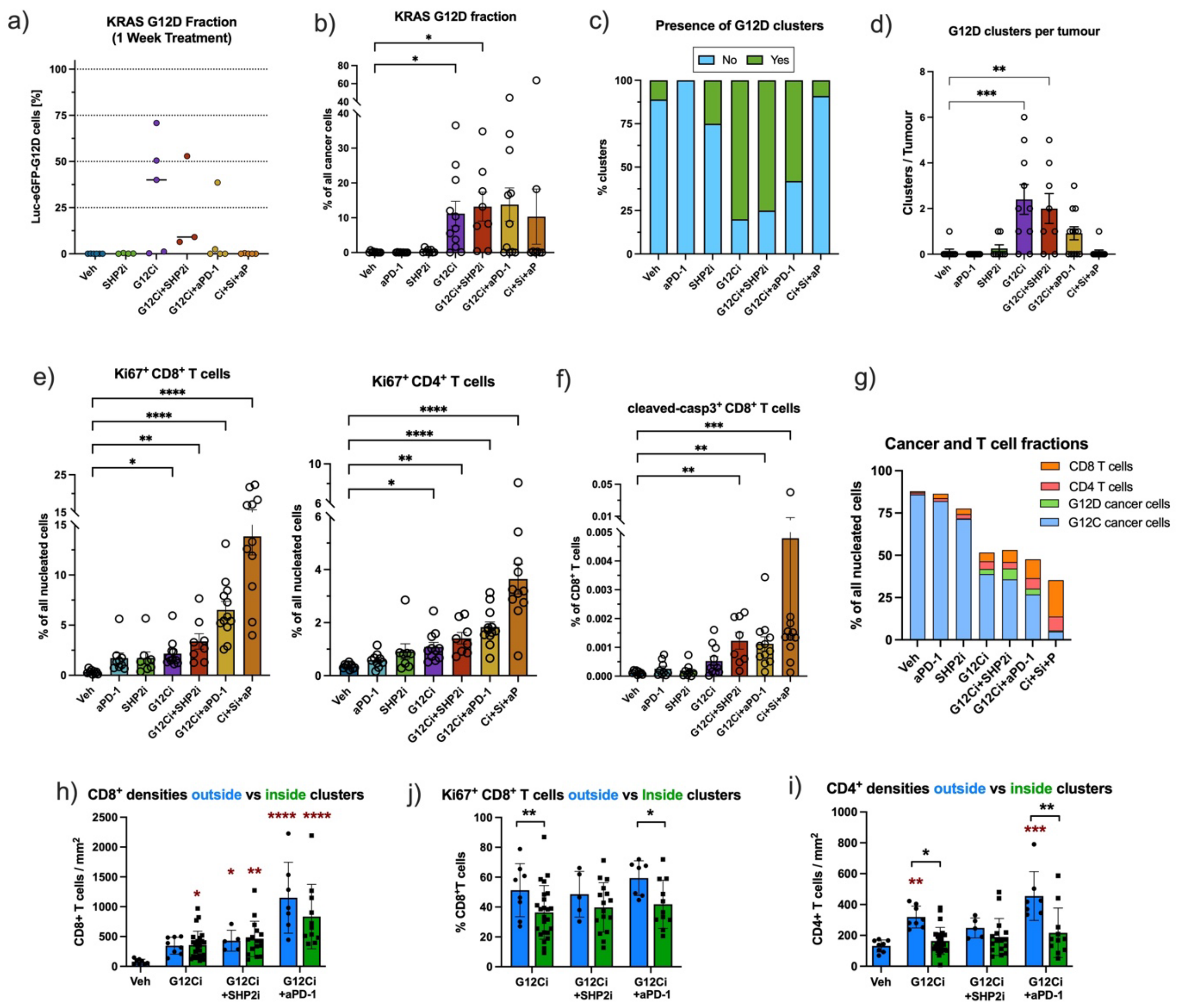

**Supplementary Figure 6. Spatial analysis of mixed KPAR1.3 subcutaneous tumours**

a) Mixed subcutaneous tumours engrafted with BFP KRAS G12C cells plus a fraction of 0.04% Luc-eGFP KRAS G12D cells. Fraction of live Luc-eGFP KRAS G12D cells within all cancer cells after one week of treatment was determined via flow cytometry. Treatments are the same as panels b-f.

(b-f) Mixed subcutaneous tumours engrafted with BFP KRAS G12C cells plus a fraction of 0.04% Luc-eGFP KRAS G12D cells were treated for six days with the RAS G12C(ON) inhibitor RMC-4998 (100 mg/kg; G12Ci / Ci) with or without the SHP2 inhibitor RMC-4550 (30 mg/kg, SHP2i / Si) and/or a-PD-1 (10 mg/kg; P). RMC-4998 and RMC-4550 were dosed by oral gavage daily and a-PD-1 was administered i.p. twice a week. Tumours were analysed by multiple immunofluorescence as indicated in Figure 3a. Unless otherwise indicated dots represented individual tumours; one section per tumour was analysed (N=8-12). Statistics indicate one-way ANOVA test comparing each of the treatment groups to the vehicle; only significant comparisons are shown.

b) Fraction of KRAS G12D mutant cancer cells out of all cancer cells.

c) Percentages of tumour sections with presence of G12D clusters, defined as cell aggregates with more than 100 G12D cells.

d) Number of G12D clusters per section.

e) Fraction of Ki-67 positive $CD8^+$ and $CD4^+$ T cells out of all nucleated cells.

f) Fraction of cleaved-caspase 3 positive $CD8^+$ T cells out of all $CD8^+$ T cells.

g) Fraction of T cells and cancer of all nucleated cells. Data is plotted as average of all the tumour sections analysed (N=8-12). Individual tumour data are plotted in Figures 3b, 3c and 3g.

h-j) Quantification of h) $CD8^+$, i) Ki67-positive $CD8^+$ T cells and j) $CD4^+$ T cells in the G12D clusters (inside; green bars) compared with a representative section outside the cluster (blue bars). The dots in the blue bars show one representative section per tumour and the dots in the green bars correspond to all the clusters identified (defined in panel c). Statistics in black indicate two-way ANOVA test comparing the inside and outside areas. For panels h and j, statistics in red indicate one-way ANOVA test comparing each of the treatment group to the vehicle. Only significant comparisons are shown.

### Supplementary Figure 7

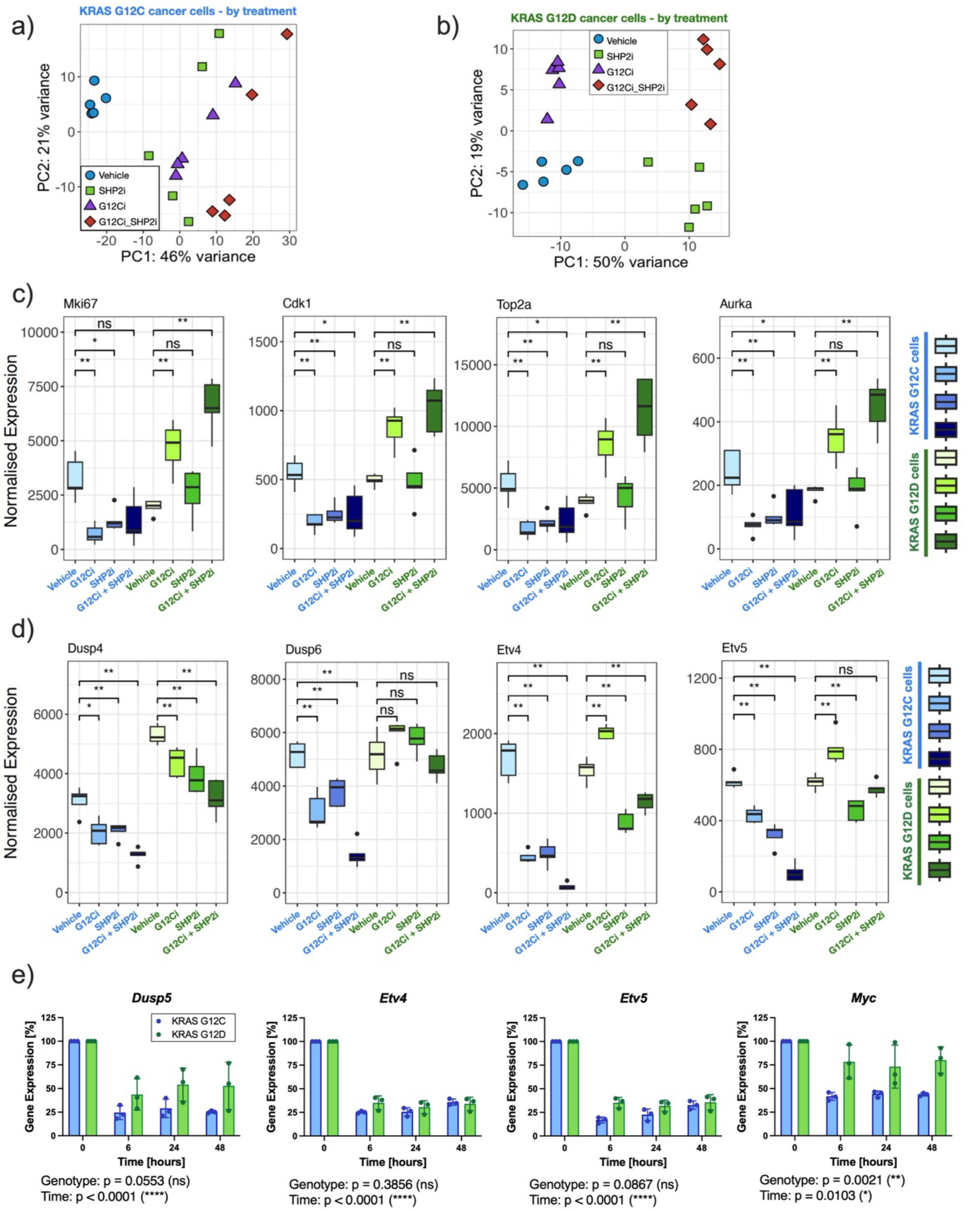

**Supplementary Figure 7. Analysis of proliferation-associated tumour cell signatures**
Mixed subcutaneous tumours were engrafted with BFP KRAS G12C cells plus 7.5% Luc-eGFP KRAS G12D cells. Mice were treated for six days with the RAS G12C(ON) inhibitor RMC-4998 (100 mg/kg; G12Ci) with or without the SHP2 inhibitor RMC-4550 (30 mg/kg; SHP2i). After treatment cancer cells were sorted and bulk RNAseq was performed.
(a-b) PCA analysis across all samples for a) KRAS G12C and b) KRAS G12D mutant tumour cell samples, with treatment groups indicated by shapes and colours. Each dot represents one independent biological replicate from a separate mouse.
(c-d) Expression of individual genes related to (c) proliferation and cell cycle progression and (d) KRAS downstream targets visualised as box plots indicating median, interquartile range. Whiskers represent the largest and smallest quartile and outliers are displayed as individual dots. Statistical significance between vehicle-treated and inhibitor-treated samples was assessed using pairwise Wilcoxon rank-sum tests (two-sided).
e) qPCR analysis of transcripts for *Dusp5*, *Etv4*, *Etv5* and *Myc* in KRAS G12C or G12D KPAR1.3 cells treated in vitro (2% FBS) with 1 µM SHP2i (RMC-4550) for indicated time points. Data was normalised to DMSO-treated cells. Mean ± SD, n=3, tr(n)=2. Two-way ANOVA test was performed to assess genotype (G12C vs G12D) and time differences.

### Supplementary Figure 8

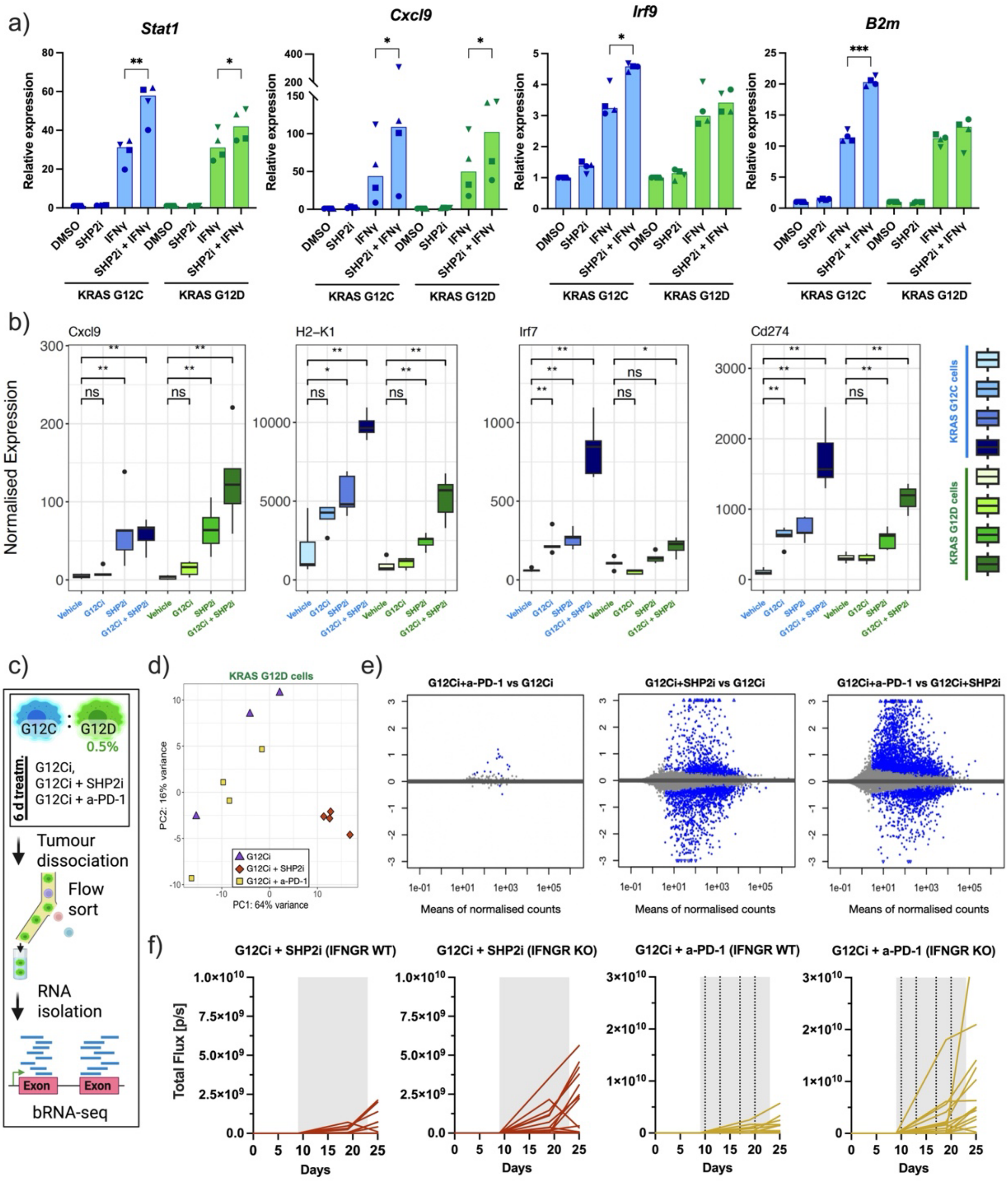

**Supplementary Figure 8. RNA-seq analysis of inflammation-associated tumour cell states**

a) KRAS G12C or G12D KPAR1.3 cells were treated in vitro for 24h with DMSO, 1μM RMC-4550 (SHP2i), IFNγ (10 ng/ml) or the combination of both and qPCR analysis of the indicated genes was performed. Mean ± SD, n=4 (indicated with different symbols), tr(n)=2. One-way ANOVA test was performed to compare IFN vs SHP2 + IFN conditions.

(b) Expression of individual genes related to interferon responses visualised from RNAseq as described in Figure 4a. Box plots indicate median, interquartile range. Whiskers represent the largest and smallest quartile and outliers are displayed as individual dots. Statistical significance between vehicle-treated and inhibitor-treated samples was assessed using pairwise Wilcoxon rank-sum tests (two-sided).

c) Schematic of experimental workflow for panels (d-e). Mixed subcutaneous tumours were engrafted with BFP KRAS G12C cells plus 0.5% Luc-eGFP KRAS G12D cells. Mice were treated for six days with the RAS G12C(ON) inhibitor RMC-4998 (100 mg/kg; C12Ci) with or without the SHP2 inhibitor RMC-4550 (30 mg/kg; SPH2i) or a-PD-1 (10 mg/kg). After treatment KRAS G12D cells were sorted and bulk RNAseq was performed.

d) PCA analysis of KRAS G12D mutant tumour cells across treatment groups.

e) MA-plots of secondary treatment effects, visualising the log2 fold changes over the mean expression of normalised gene counts. The later term in the respective plot titles is considered as “baseline” (set as 0 on x-axis). Data points on plot are coloured blue if padj ≤0.1. If data points fall out of the range of the defined y-axis they are plotted as triangles.

f) Bioluminescence scan data over time of mixed subcutaneous tumours engrafted with BFP KRAS G12C cells plus 0.04% of Luc-eGFP KRAS G12D cells expressing wild-type (WT) or knock-out (KO) IFNGR and treated for two weeks with 100 mg/kg RMC-4998 in combination with 30 mg/kg RMC-4550 (G12Ci + SHP2i) or 10 mg/kg anti-PD-1 (G12Ci + a-PD-1). Grey area indicates treatment period.

### Supplementary Figure 9

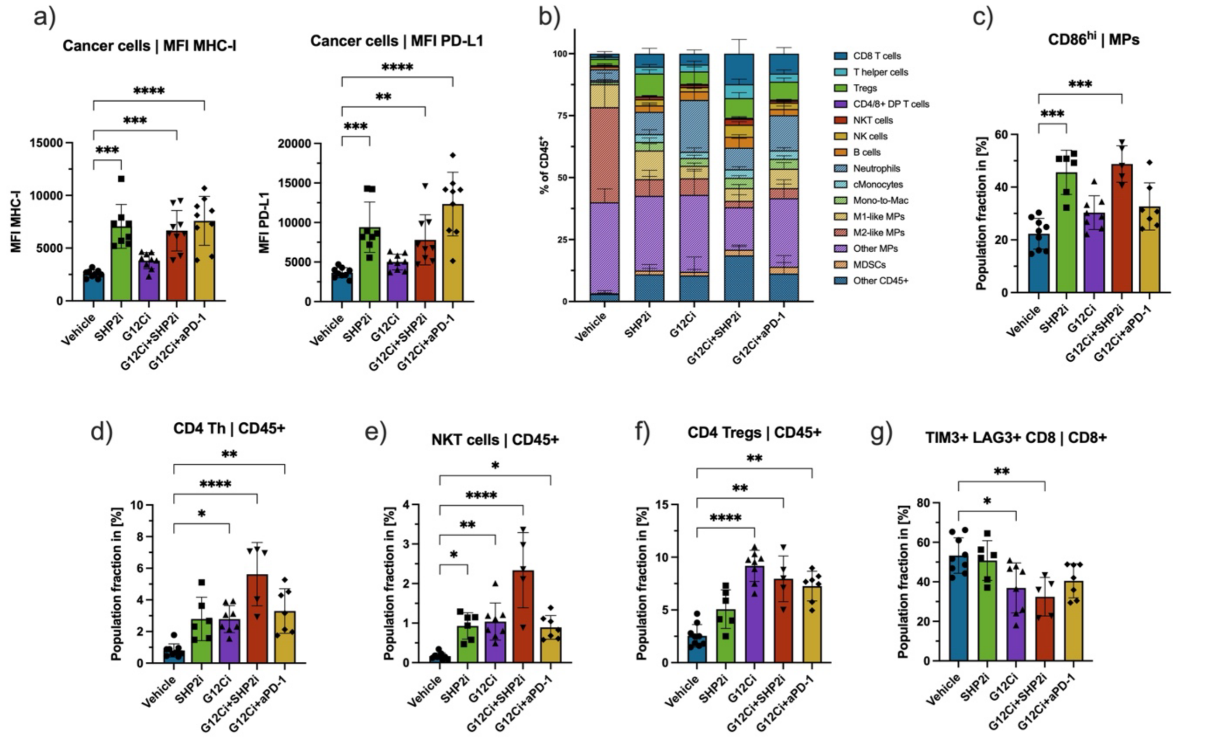


**Supplementary Figure 9. TME phenotyping via spectral flow cytometry**

Mixed subcutaneous tumours were engrafted with BFP KRAS G12C cells plus 0.2% Luc-eGFP KRAS G12D cells. Mice were treated for six days with the RAS G12C(ON) inhibitor RMC-4998 (100 mg/kg; G12Ci / Ci) with or without the SHP2 inhibitor RMC-4550 (30 mg/kg; SHP2i / Si) or a-PD-1 (10 mg/kg; P). After treatment flow cytometry of tumours was performed.

a) Surface expression (mean fluorescence intensity, MFI) of MHC-I (left) and PD-L1 (right) in cancer cells.

b) Mean percentages of $CD45^+$ immune cell sub-types by treatment groups, whiskers indicate SD within each population across individual samples. Abbreviations from top to bottom: ‘Tregs’: T regulatory cells’; ‘$CD8^+$ $CD4^+$ double-positive (DP) T cells’, ‘NKT’: natural killer T cells; ‘NK’: natural killer cells; ‘cMonocytes’: classical ($CD64^+$) Monocytes; ‘Mono-to-Mac’: a population of monocytes in transition-phase to differentiating macrophages; ‘M1-like MPs’: $Arg1^{lo}$ $MHC\text{-}II^{hi}$ Macrophages; ‘M2-like MPs’: $Arg1^{hi}$ $MHC\text{-}II^{lo}$ Macrophages; ‘Other MPs’: Other Macrophages; ‘MDSCs’: myeloid derived suppressor cells.

(c-g) For all graphs, dots represent individual tumours; bars show mean + SD; one-way ANOVA Kruskal-Wallis test was performed, calculating pairwise comparison between vehicle and individual treatment groups; only significant comparisons are indicated.

c) Fraction of CD86 high cells as percentage of all macrophages (MPs).

d) Fraction of $CD4^+$ T helper cells out of all immune cells ($CD45^+$).

e) Fraction of NKT cells out of all $CD45^+$ cells.

f) Fraction of regulatory T cells (Tregs) out of all $CD45^+$ cells.

g) Fraction of terminally differentiated/exhausted ($TIM3^+$/$LAG3^+$) $CD8^+$ T cells out of all $CD8^+$ T cells.

### Supplementary Figure 10

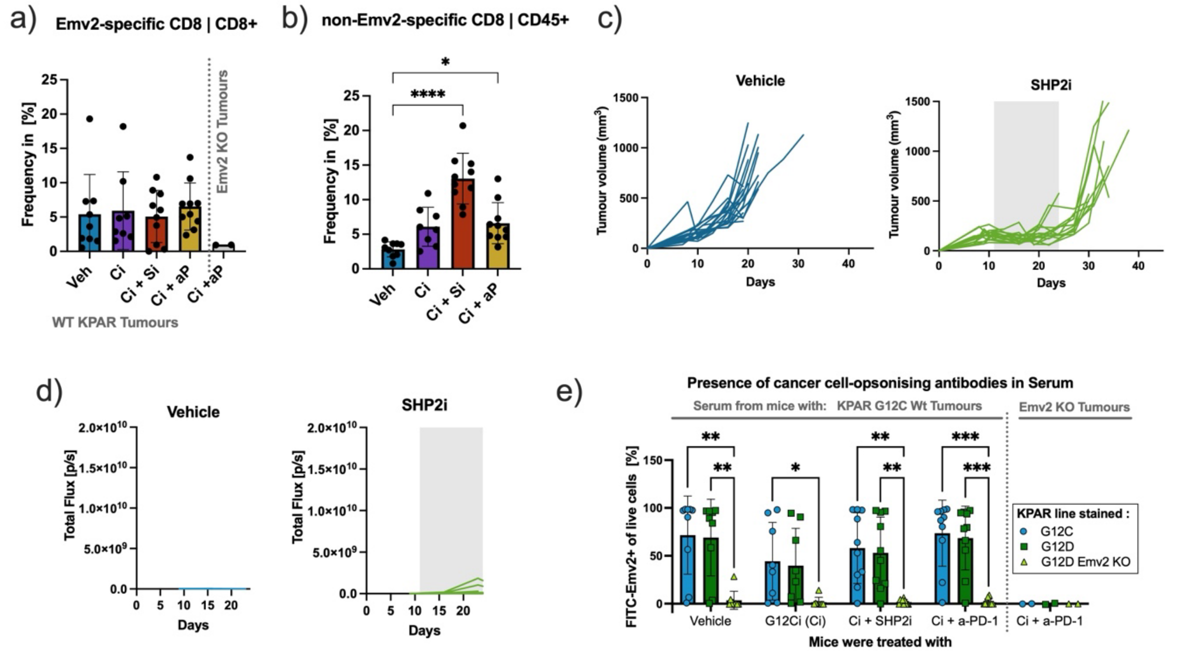


**Supplementary Figure 10. Adaptive immunity is required for durable responses**
a-b) Mice with KRAS G12C KPAR1.3 wild-type or Emv2 knockout tumours were treated for six days with the RAS G12C(ON) inhibitor RMC-4998 (100 mg/kg; G12Ci / Ci) with or without the SHP2 inhibitor RMC-4550 (30 mg/kg; SHP2i) or a-PD-1 (10 mg/kg; aP). a) Frequency of Emv2-specific $CD8^+$ T cells of all $CD8^+$ T cells. Additional control group with KPAR1.3 KRAS G12D Emv2-KO tumours is shown. b) Frequency of non-Emv2-specific $CD8^+$ T cells of all immune cells ($CD45^+$). Each dot represents one individual tumour; bar graphs indicate mean ± SD; One-way ANOVA Kruskal-Wallis test comparing vehicle to each of the treated conditions; only significant comparisons are shown.
(c-d) Mixed subcutaneous tumours were engrafted with BFP KRAS G12C cells plus a fraction of 0.04% Luc-eGFP KPAR.EP2 cells in $Rag1^{-/-}$ GH-mice. Mice were treated for two weeks with vehicle or the SHP2 inhibitor RMC-4550 (30 mg/kg; SHP2i). c) Tumour growth of individual tumours across indicated treatment groups. Treatment interval is highlighted in grey. Additional treatment groups are shown in Figure 6d. d) Bioluminescence scans, indicating the relative abundance of Luc-eGFP KRAS G12D cells over time, measured in total flux, photons per second [p/s]. Additional treatment groups are shown in Figure 6e.
e) Median fluorescent intensity (MFI) of serum-stained un-transduced KPAR1.3 KRAS G12C, G12D, or G12D Emv2 KO tumours cancer cells, assessed via flow cytometric analysis. Serum was collected from mice treated as indicated in (a). All samples were normalised to the MFI of matching serum from obtained from naïve mice. Results are coloured by genotype and sorted by treatment group. Serum from a control group of mice harbouring Emv2 KO tumours was also included as additional negative control for epitope specificity. Each dot indicates serum from one mouse, the height bars indicate mean across samples within each group + SEM. 2way ANOVA test was performed, between opsonised cell lines within each treatment condition, comparing serum samples from KPAR1.3 G12D Emv2 KO cells to the KPAR1.3 wt groups; only significant comparisons are indicated.

### Supplementary Figure 11

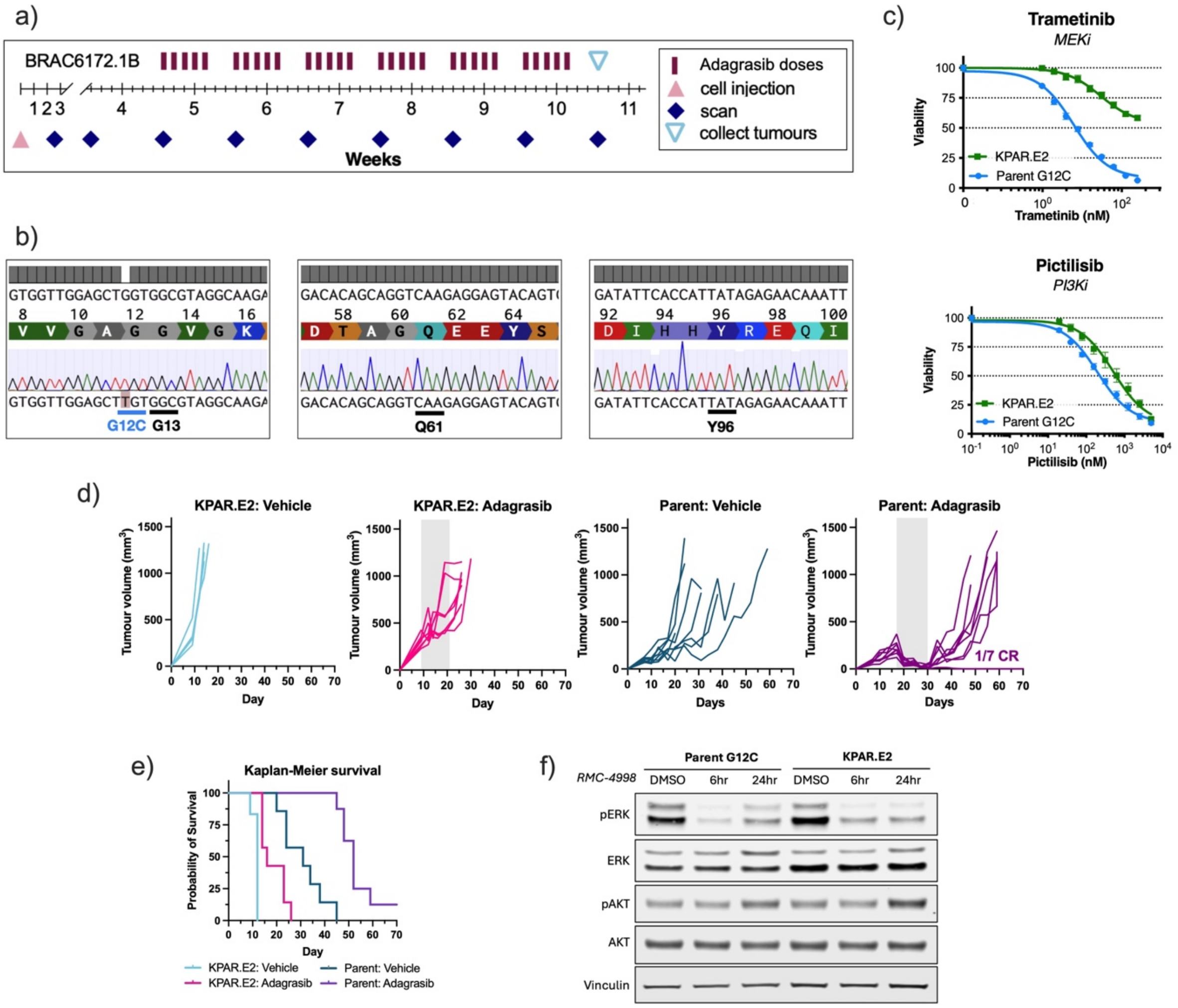


**Supplementary Figure 11. Establishment of G12Ci resistant KPAR.E2 cell line**

a) Treatment scheme to generate adagrasib-resistant KPAR1.3 KRAS G12C-derived lung tumours in immune-competent mice.

b) Sanger sequencing for KRAS transcript in KPAR.E2 cells (bottom) compared with reference sequence (top), highlighting the retained KRAS G12C mutation as well as unaltered aminoacid sequence at other hotspot sites.

c) 72-hour viability assays for targeted inhibitors trametinib (MEKi) and pictilisib (PI3Ki) at indicated drug concentrations in parental KPAR1.3 KRAS G12C cell line (Parent G12C) and G12Ci-resistant KPAR.E2 cells. Error bars indicate SEM with n=3.

d) Tumour growth of KPAR.E2 or KPAR1.3 KRAS G12C (parent) subcutaneous tumours treated for 2 weeks (grey area) with either vehicle or the KRAS G12Ci adagrasib (50 mg/kg). Complete responders (CR) indicated on respective plot.

e) Probability of survival for mice across all treatment groups from (e).

f) Westen blot of cells treated at different time points with 100 nM RMC-4998.

### Supplementary Figure 12

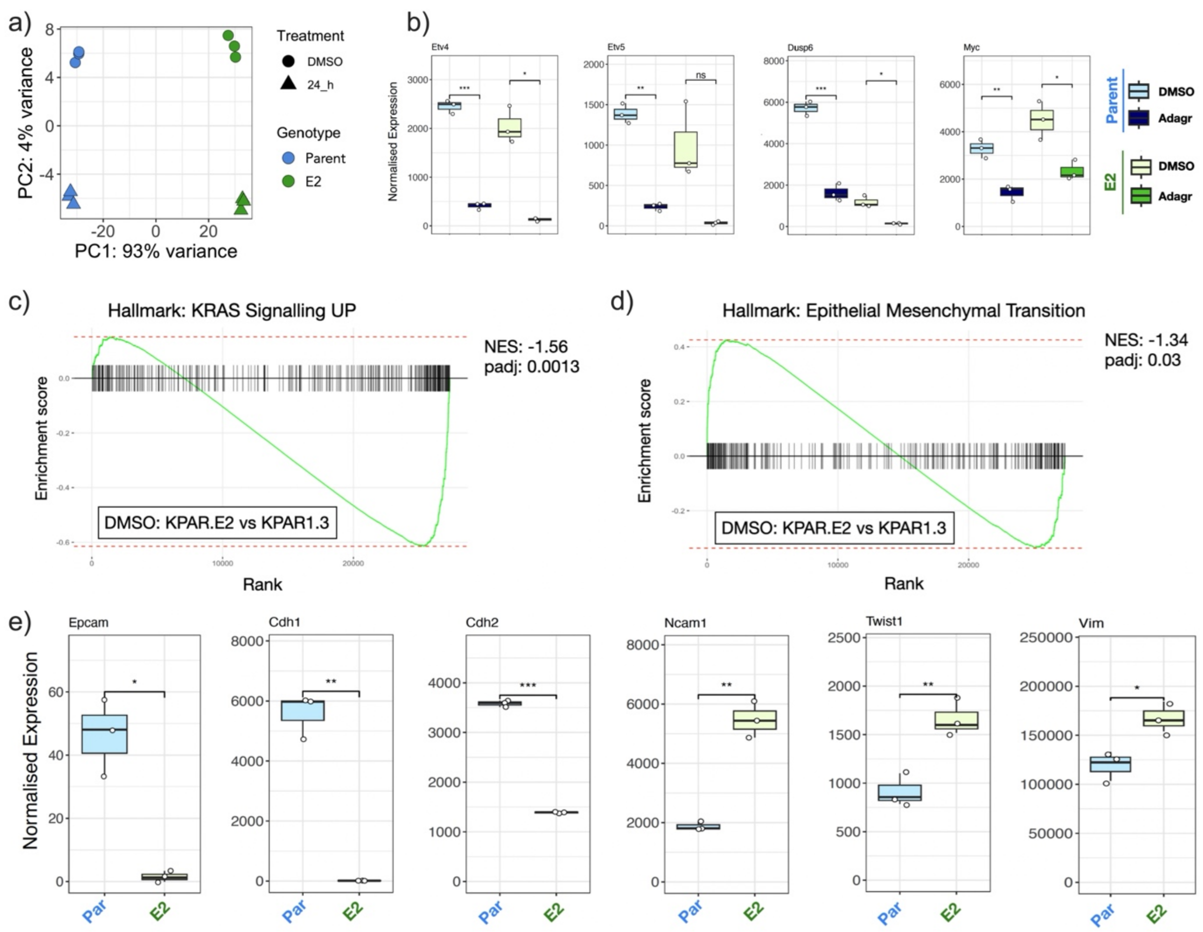


**Supplementary Figure 12. RNA-seq analysis of KPAR.E2 cell line**

a) Principal component analysis (PCA) of RNA-sequencing samples of parental KPAR1.3 KRAS G12C and KPAR.E2 cell lines treated either with DMSO or 100 nM adagrasib for 24h.

b) Expression of individual genes related to downstream transcriptional targets of KRAS signalling. Visualised as box plots, indicating median, interquartile range, whiskers represent largest and smallest quartile and outliers are displayed as individual dots.

c-d) Gene set enrichment analysis plots for indicated gene sets of the msigdb hallmarks collection, (c) KRAS Signalling UP and (d) Epithelial to mesenchymal transition.

e) Expression of individual genes related to EMT-associated genes. Visualised as box plots, indicating median, interquartile range, whiskers represent largest and smallest quartile and outliers are displayed as individual dots with x-axis labels as indicated on the right of respective panels.

### Supplementary Figure 13

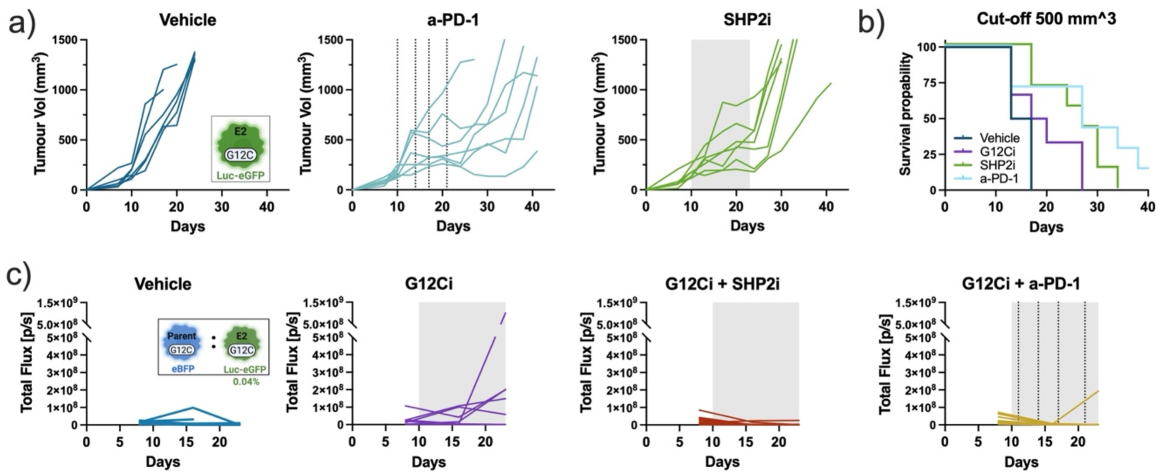


**Supplementary Figure 13. Treatment of pure and mixed KPAR.E2 tumours**
a) Tumour growth over time of individual KPAR.E2-derived subcutaneous tumours for indicated treatment groups: vehicle, a-PD-1 (10 mg/kg) and SHP2 inhibitor RMC-4550 (30mg/kg, SHP2i).
b) Probability of survival for indicated treatment groups of the same experiment. Tumour growth for G12Ci (100 mg/kg RMC-4998) is shown in Figure 7f.
c) Bioluminescence scans of co-engrafted tumours with a subpopulation of 0.04% Luc-eGFP KPAR.E2 cells by treatment group, indicating the relative abundance of KPAR.E2 cells over time, measured in total flux, photons per second [p/s]. Mice were treated for 2 weeks (grey area) with the RAS G12C(ON) inhibitor RMC-4998 (100 mg/kg; G12Ci) with or without the SHP2 inhibitor RMC-4550 (30 mg/kg; SHP2i) or a-PD-1 (10 mg/kg).

### Supplementary Tables

#### Table S1: Predicted MHC-I binding affinities.

| Peptides containing Y67H mutation | | | | | |
|---|---|---|---|---|---|
| **allele** | **start** | **end** | **length** | **peptide** | **ann_ic50** |
| H-2-Kb | 62 | 69 | 8 | VTTLT**H**GV | 7742.85 |
| H-2-Db | 67 | 76 | 10 | **H**GVQCFSRYP | 27951.49 |
| H-2-Kb | 63 | 70 | 8 | TTLT**H**GVQ | 26026.48 |
| H-2-Db | 63 | 72 | 10 | TTLT**H**GVQCF | 33228.35 |
| H-2-Db | 66 | 75 | 10 | T**H**GVQCFSRY | 41515.82 |
| H-2-Kb | 61 | 68 | 8 | LVTTLT**H**G | 34313.47 |
| H-2-Db | 60 | 69 | 10 | TLVTTLT**H**GV | 30960.01 |
| H-2-Kb | 65 | 72 | 8 | LT**H**GVQCF | 27047.65 |
| H-2-Db | 61 | 70 | 10 | LVTTLT**H**GVQ | 45032.54 |
| H-2-Kb | 67 | 74 | 8 | **H**GVQCFSR | 34708.9 |
| H-2-Kb | 66 | 73 | 8 | T**H**GVQCFS | 37290.64 |
| H-2-Db | 59 | 68 | 10 | PTLVTTLT**H**G | 44718.85 |
| H-2-Kb | 64 | 71 | 8 | TLT**H**GVQC | 34352.11 |
| H-2-Db | 64 | 73 | 10 | TLT**H**GVQCFS | 46038.05 |
| H-2-Db | 58 | 67 | 10 | WPTLVTTLT**H** | 45902.74 |
| H-2-Db | 62 | 71 | 10 | VTTLT**H**GVQC | 45221.47 |
| H-2-Kb | 60 | 67 | 8 | TLVTTLT**H** | 36699.04 |
| H-2-Db | 65 | 74 | 10 | LT**H**GVQCFSR | 47743.04 |

| Peptides containing Y146F mutation | | | | | |
|---|---|---|---|---|---|
| **allele** | **start** | **end** | **length** | **peptide** | **ann_ic50** |
| H-2-Kb | 144 | 151 | 8 | YN**F**NSHNV | 952.48 |
| H-2-Db | 144 | 153 | 10 | YN**F**NSHNVYI | 4045.8 |
| H-2-Db | 146 | 155 | 10 | **F**NSHNVYIMA | 18589.52 |
| H-2-Db | 145 | 154 | 10 | N**F**NSHNVYIM | 10562.59 |
| H-2-Db | 142 | 151 | 10 | LEYN**F**NSHNV | 21595.95 |
| H-2-Db | 140 | 149 | 10 | HKLEYN**F**NSH | 39818.49 |
| H-2-Db | 143 | 152 | 10 | EYN**F**NSHNVY | 43582.46 |
| H-2-Db | 138 | 147 | 10 | LGHKLEYN**F**N | 40727.86 |
| H-2-Db | 141 | 150 | 10 | KLEYN**F**NSHN | 42843.3 |
| H-2-Kb | 142 | 149 | 8 | LEYN**F**NSH | 26538.6 |
| H-2-Db | 137 | 146 | 10 | ILGHKLEYN**F** | 32346.83 |
| H-2-Kb | 140 | 147 | 8 | HKLEYN**F**N | 38359.12 |
| H-2-Kb | 141 | 148 | 8 | KLEYN**F**NS | 29873.42 |
| H-2-Kb | 139 | 146 | 8 | GHKLEYN**F** | 30312.32 |
| H-2-Kb | 145 | 152 | 8 | N**F**NSHNVY | 34272.29 |
| H-2-Db | 139 | 148 | 10 | GHKLEYN**F**NS | 48084.14 |
| H-2-Kb | 143 | 150 | 8 | EYN**F**NSHN | 39103.78 |

### Table S2: GSEA significant targets across all treatment groups

| KRAS | Treatment | Pathway | Exp genes | pval | padj | $log_{10}$(padj) | log2err | ES | NES |
|---|---|---|---|---|---|---|---|---|---|
| G12C | SHP2i | MYC TARGETS V1 | 254 | 8.098E-15 | 3.968E-13 | 12.401 | 0.987 | -0.826 | -2.099 |
| G12C | SHP2i | E2F TARGETS | 266 | 2.128E-14 | 5.214E-13 | 12.283 | 0.976 | -0.817 | -2.068 |
| G12C | SHP2i | G2M CHECKPOINT | 298 | 7.401E-11 | 9.066E-10 | 9.043 | 0.839 | -0.763 | -1.969 |
| G12C | SHP2i | INTERFERON GAMMARESPONSE | 311 | 3.763E-13 | 6.146E-12 | 11.211 | 0.933 | 0.850 | 1.709 |
| G12C | SHP2i | INTERFERON ALPHARESPONSE | 178 | 1.375E-09 | 1.347E-08 | 7.871 | 0.788 | 0.866 | 1.707 |
| G12C | SHP2i | MYC TARGETS V2 | 68 | 3.317E-03 | 9.029E-03 | 2.044 | 0.432 | -0.728 | -1.604 |
| G12C | SHP2i | EPITHELIALMESENCHYMALTRANSITION | 330 | 3.846E-08 | 3.141E-07 | 6.503 | 0.720 | 0.793 | 1.595 |
| G12C | SHP2i | COAGULATION | 235 | 1.963E-05 | 1.374E-04 | 3.862 | 0.576 | 0.779 | 1.553 |
| G12C | SHP2i | IL6 JAK STAT3SIGNALING | 121 | 8.151E-04 | 2.760E-03 | 2.559 | 0.477 | 0.783 | 1.501 |
| G12C | SHP2i | TNFA SIGNALING VIANFKB | 274 | 4.458E-05 | 2.731E-04 | 3.564 | 0.557 | 0.748 | 1.501 |
| G12C | SHP2i | ALLOGRAFT REJECTION | 313 | 2.986E-04 | 1.330E-03 | 2.876 | 0.498 | 0.723 | 1.455 |
| G12C | SHP2i | MYOGENESIS | 363 | 1.797E-04 | 9.782E-04 | 3.010 | 0.519 | 0.715 | 1.441 |
| G12C | SHP2i | HYPOXIA | 317 | 5.718E-04 | 2.335E-03 | 2.632 | 0.477 | 0.711 | 1.431 |
| G12C | SHP2i | XENOBIOTICMETABOLISM | 352 | 2.128E-04 | 1.043E-03 | 2.982 | 0.519 | 0.708 | 1.426 |
| G12C | SHP2i | INFLAMMATORYRESPONSE | 314 | 6.481E-04 | 2.443E-03 | 2.612 | 0.477 | 0.695 | 1.397 |
| G12C | SHP2i | APICAL JUNCTION | 373 | 8.449E-04 | 2.760E-03 | 2.559 | 0.477 | 0.684 | 1.380 |
| G12C | SHP2i | UV RESPONSE DN | 307 | 1.115E-03 | 3.414E-03 | 2.467 | 0.455 | 0.684 | 1.376 |
| G12C | SHP2i | ESTROGEN RESPONSEEARLY | 327 | 3.156E-03 | 9.029E-03 | 2.044 | 0.432 | 0.680 | 1.367 |
| G12C | SHP2i | BILE ACID METABOLISM | 171 | 1.594E-02 | 3.396E-02 | 1.469 | 0.352 | 0.696 | 1.367 |
| G12C | SHP2i | ADIPOGENESIS | 293 | 5.135E-03 | 1.198E-02 | 1.921 | 0.407 | 0.678 | 1.360 |
| G12C | SHP2i | IL2 STAT5 SIGNALING | 308 | 3.845E-03 | 9.613E-03 | 2.017 | 0.432 | 0.675 | 1.357 |
| G12C | SHP2i | COMPLEMENT | 359 | 3.924E-03 | 9.613E-03 | 2.017 | 0.407 | 0.668 | 1.346 |
| G12C | SHP2i | TGF BETA SIGNALING | 100 | 3.833E-02 | 7.825E-02 | 1.107 | 0.249 | 0.715 | 1.342 |
| G12C | SHP2i | ESTROGEN RESPONSELATE | 342 | 1.220E-02 | 2.717E-02 | 1.566 | 0.381 | 0.641 | 1.290 |
| G12C | KRASi | E2F TARGETS | 266 | 4.432E-12 | 2.172E-10 | 9.663 | 0.887 | -0.820 | -1.987 |
| G12C | KRASi | MYC TARGETS V1 | 254 | 1.705E-10 | 3.121E-09 | 8.506 | 0.827 | -0.803 | -1.937 |
| G12C | KRASi | G2M CHECKPOINT | 298 | 2.547E-10 | 3.121E-09 | 8.506 | 0.814 | -0.786 | -1.934 |
| G12C | KRASi | INTERFERON GAMMARESPONSE | 311 | 1.985E-10 | 3.121E-09 | 8.506 | 0.827 | 0.787 | 1.863 |
| G12C | KRASi | INTERFERON ALPHARESPONSE | 178 | 7.373E-07 | 7.226E-06 | 5.141 | 0.659 | 0.789 | 1.780 |
| G12C | KRASi | TNFA SIGNALING VIANFKB | 274 | 1.134E-06 | 9.260E-06 | 5.033 | 0.644 | 0.730 | 1.714 |
| G12C | KRASi | COAGULATION | 235 | 1.233E-03 | 5.491E-03 | 2.260 | 0.455 | 0.670 | 1.552 |
| G12C | KRASi | IL6 JAK STAT3SIGNALING | 121 | 5.435E-03 | 2.049E-02 | 1.689 | 0.407 | 0.702 | 1.520 |
| G12C | KRASi | ESTROGEN RESPONSEEARLY | 327 | 4.151E-04 | 2.905E-03 | 2.537 | 0.498 | 0.636 | 1.502 |
| G12C | KRASi | INFLAMMATORYRESPONSE | 314 | 8.415E-04 | 4.581E-03 | 2.339 | 0.477 | 0.626 | 1.480 |
| G12C | KRASi | ALLOGRAFT REJECTION | 313 | 7.762E-04 | 4.581E-03 | 2.339 | 0.477 | 0.619 | 1.465 |
| G12C | KRASi | HYPOXIA | 317 | 1.138E-03 | 5.491E-03 | 2.260 | 0.455 | 0.617 | 1.456 |
| G12C | KRASi | TGF BETA SIGNALING | 100 | 1.544E-02 | 4.730E-02 | 1.325 | 0.381 | 0.691 | 1.452 |
| G12C | KRASi | MYC TARGETS V2 | 68 | 3.581E-02 | 8.356E-02 | 1.078 | 0.322 | -0.687 | -1.430 |
| G12C | KRASi | ADIPOGENESIS | 293 | 6.227E-03 | 2.179E-02 | 1.662 | 0.407 | 0.594 | 1.399 |
| G12C | KRASi | EPITHELIALMESENCHYMALTRANSITION | 330 | 2.984E-03 | 1.219E-02 | 1.914 | 0.432 | 0.585 | 1.383 |
| G12C | KRASi | XENOBIOTICMETABOLISM | 352 | 9.041E-03 | 2.953E-02 | 1.530 | 0.381 | 0.557 | 1.322 |
| G12C | KRASi | IL2 STAT5 SIGNALING | 308 | 2.589E-02 | 6.677E-02 | 1.175 | 0.352 | 0.556 | 1.314 |
| G12C | KRASi | COMPLEMENT | 359 | 2.433E-02 | 6.677E-02 | 1.175 | 0.352 | 0.543 | 1.296 |
| G12C | KRASi | ESTROGEN RESPONSELATE | 342 | 2.506E-02 | 6.677E-02 | 1.175 | 0.352 | 0.540 | 1.286 |
| G12C | KRASi | MYOGENESIS | 363 | 2.867E-02 | 7.025E-02 | 1.153 | 0.352 | 0.527 | 1.260 |
| G12C | KRASi | BILE ACID METABOLISM | 171 | 7.465E-02 | 1.396E-01 | 0.855 | 0.217 | 0.561 | 1.259 |
| G12C | KRASi | UV RESPONSE DN | 307 | 6.491E-02 | 1.272E-01 | 0.895 | 0.234 | 0.520 | 1.227 |
| G12C | KRASi | APICAL JUNCTION | 373 | 3.135E-01 | 3.746E-01 | 0.426 | 0.096 | 0.437 | 1.048 |
| G12C | KRASi_SHP2i | MYC TARGETS V1 | 254 | 9.197E-13 | 1.502E-11 | 10.823 | 0.910 | -0.822 | -1.984 |
| G12C | KRASi_SHP2i | INTERFERON GAMMARESPONSE | 311 | 2.236E-20 | 1.096E-18 | 17.960 | 1.169 | 0.868 | 1.970 |
| G12C | KRASi_SHP2i | INTERFERON ALPHARESPONSE | 178 | 8.384E-15 | 2.054E-13 | 12.687 | 0.987 | 0.890 | 1.936 |
| G12C | KRASi_SHP2i | E2F TARGETS | 266 | 2.291E-10 | 2.806E-09 | 8.552 | 0.827 | -0.790 | -1.918 |
| G12C | KRASi_SHP2i | G2M CHECKPOINT | 298 | 2.749E-08 | 2.694E-07 | 6.570 | 0.734 | -0.729 | -1.796 |
| G12C | KRASi_SHP2i | MYC TARGETS V2 | 68 | 4.896E-03 | 1.599E-02 | 1.796 | 0.407 | -0.777 | -1.615 |
| G12C | KRASi_SHP2i | ALLOGRAFT REJECTION | 313 | 7.219E-05 | 5.896E-04 | 3.229 | 0.538 | 0.673 | 1.533 |
| G12C | KRASi_SHP2i | IL6 JAK STAT3SIGNALING | 121 | 4.497E-03 | 1.599E-02 | 1.796 | 0.407 | 0.718 | 1.499 |
| G12C | KRASi_SHP2i | COAGULATION | 235 | 2.786E-04 | 1.950E-03 | 2.710 | 0.498 | 0.674 | 1.496 |
| G12C | KRASi_SHP2i | ADIPOGENESIS | 293 | 4.457E-04 | 2.427E-03 | 2.615 | 0.498 | 0.657 | 1.486 |
| G12C | KRASi_SHP2i | TNFA SIGNALING VIANFKB | 274 | 4.278E-04 | 2.427E-03 | 2.615 | 0.498 | 0.656 | 1.480 |
| G12C | KRASi_SHP2i | INFLAMMATORYRESPONSE | 314 | 7.405E-04 | 3.299E-03 | 2.482 | 0.477 | 0.650 | 1.478 |
| G12C | KRASi_SHP2i | ESTROGEN RESPONSEEARLY | 327 | 5.891E-04 | 2.886E-03 | 2.540 | 0.477 | 0.634 | 1.449 |
| G12C | KRASi_SHP2i | MYOGENESIS | 363 | 1.687E-03 | 6.890E-03 | 2.162 | 0.455 | 0.611 | 1.403 |
| G12C | KRASi_SHP2i | XENOBIOTICMETABOLISM | 352 | 4.842E-03 | 1.599E-02 | 1.796 | 0.407 | 0.601 | 1.378 |
| G12C | KRASi_SHP2i | ESTROGEN RESPONSELATE | 342 | 1.077E-02 | 3.299E-02 | 1.482 | 0.381 | 0.589 | 1.348 |
| G12C | KRASi_SHP2i | BILE ACID METABOLISM | 171 | 2.311E-02 | 6.292E-02 | 1.201 | 0.352 | 0.615 | 1.335 |
| G12C | KRASi_SHP2i | HYPOXIA | 317 | 2.176E-02 | 6.273E-02 | 1.203 | 0.352 | 0.581 | 1.323 |
| G12C | KRASi_SHP2i | IL2 STAT5 SIGNALING | 308 | 2.853E-02 | 7.358E-02 | 1.133 | 0.352 | 0.567 | 1.287 |
| G12C | KRASi_SHP2i | EPITHELIALMESENCHYMALTRANSITION | 330 | 3.051E-02 | 7.474E-02 | 1.126 | 0.352 | 0.559 | 1.277 |
| G12C | KRASi_SHP2i | TGF BETA SIGNALING | 100 | 1.332E-01 | 2.088E-01 | 0.680 | 0.148 | 0.597 | 1.227 |
| G12C | KRASi_SHP2i | UV RESPONSE DN | 307 | 6.208E-02 | 1.267E-01 | 0.897 | 0.209 | 0.539 | 1.224 |
| G12C | KRASi_SHP2i | COMPLEMENT | 359 | 8.432E-02 | 1.530E-01 | 0.815 | 0.175 | 0.520 | 1.193 |
| G12C | KRASi_SHP2i | APICAL JUNCTION | 373 | 1.429E-01 | 2.121E-01 | 0.673 | 0.130 | 0.497 | 1.145 |

| | | | | | | | | | |
|---|---|---|---|---|---|---|---|---|---|
| G12D | SHP2i | MYC TARGETS V1 | 254 | 2.081E-05 | 1.020E-03 | 2.992 | 0.576 | -0.814 | -1.810 |
| G12D | SHP2i | INTERFERON ALPHARESPONSE | 178 | 4.523E-04 | 7.387E-03 | 2.132 | 0.498 | 0.830 | 1.631 |
| G12D | SHP2i | INTERFERON GAMMARESPONSE | 311 | 1.248E-04 | 3.058E-03 | 2.515 | 0.519 | 0.802 | 1.611 |
| G12D | SHP2i | MYC TARGETS V2 | 68 | 2.282E-02 | 1.062E-01 | 0.974 | 0.352 | -0.787 | -1.609 |
| G12D | SHP2i | IL6 JAK STAT3SIGNALING | 121 | 3.103E-02 | 1.165E-01 | 0.934 | 0.352 | 0.778 | 1.487 |
| G12D | SHP2i | COAGULATION | 235 | 1.167E-02 | 8.166E-02 | 1.088 | 0.381 | 0.744 | 1.471 |
| G12D | SHP2i | TNFA SIGNALING VIANFKB | 274 | 8.770E-03 | 7.162E-02 | 1.145 | 0.381 | 0.732 | 1.459 |
| G12D | SHP2i | ESTROGEN RESPONSELATE | 342 | 5.940E-03 | 7.162E-02 | 1.145 | 0.407 | 0.716 | 1.445 |
| G12D | SHP2i | EPITHELIALMESENCHYMALTRANSITION | 330 | 8.360E-03 | 7.162E-02 | 1.145 | 0.381 | 0.709 | 1.433 |
| G12D | SHP2i | ALLOGRAFT REJECTION | 313 | 1.529E-02 | 9.364E-02 | 1.029 | 0.381 | 0.704 | 1.416 |
| G12D | SHP2i | INFLAMMATORYRESPONSE | 314 | 2.413E-02 | 1.062E-01 | 0.974 | 0.352 | 0.692 | 1.392 |
| G12D | SHP2i | IL2 STAT5 SIGNALING | 308 | 2.247E-02 | 1.062E-01 | 0.974 | 0.352 | 0.691 | 1.388 |
| G12D | SHP2i | COMPLEMENT | 359 | 2.600E-02 | 1.062E-01 | 0.974 | 0.352 | 0.672 | 1.358 |
| G12D | SHP2i | ESTROGEN RESPONSEEARLY | 327 | 3.444E-02 | 1.165E-01 | 0.934 | 0.262 | 0.665 | 1.341 |
| G12D | SHP2i | XENOBIOTICMETABOLISM | 352 | 4.138E-02 | 1.267E-01 | 0.897 | 0.238 | 0.653 | 1.319 |
| G12D | SHP2i | APICAL JUNCTION | 373 | 4.886E-02 | 1.408E-01 | 0.851 | 0.217 | 0.644 | 1.306 |
| G12D | SHP2i | BILE ACID METABOLISM | 171 | 7.255E-02 | 1.580E-01 | 0.801 | 0.181 | 0.656 | 1.288 |
| G12D | SHP2i | UV RESPONSE DN | 307 | 7.317E-02 | 1.580E-01 | 0.801 | 0.177 | 0.619 | 1.243 |
| G12D | SHP2i | MYOGENESIS | 363 | 7.657E-02 | 1.580E-01 | 0.801 | 0.171 | 0.613 | 1.242 |
| G12D | SHP2i | TGF BETA SIGNALING | 100 | 1.480E-01 | 2.686E-01 | 0.571 | 0.129 | 0.649 | 1.221 |
| G12D | SHP2i | ADIPOGENESIS | 293 | 1.036E-01 | 2.031E-01 | 0.692 | 0.146 | 0.602 | 1.207 |
| G12D | SHP2i | G2M CHECKPOINT | 298 | 2.000E-01 | 3.063E-01 | 0.514 | 0.271 | -0.531 | -1.188 |
| G12D | SHP2i | E2F TARGETS | 266 | 2.550E-01 | 3.471E-01 | 0.460 | 0.231 | -0.516 | -1.149 |
| G12D | SHP2i | HYPOXIA | 317 | 6.694E-01 | 7.003E-01 | 0.155 | 0.039 | 0.455 | 0.915 |
| G12D | KRASi | E2F TARGETS | 266 | 1.166E-06 | 5.712E-05 | 4.243 | 0.644 | 0.855 | 1.963 |
| G12D | KRASi | G2M CHECKPOINT | 298 | 4.935E-04 | 1.209E-02 | 1.918 | 0.477 | 0.777 | 1.794 |
| G12D | KRASi | MYC TARGETS V1 | 254 | 5.038E-03 | 8.229E-02 | 1.085 | 0.407 | 0.740 | 1.692 |
| G12D | KRASi | HYPOXIA | 317 | 1.514E-02 | 1.854E-01 | 0.732 | 0.381 | -0.674 | -1.562 |
| G12D | KRASi | MYC TARGETS V2 | 68 | 3.208E-02 | 3.144E-01 | 0.503 | 0.322 | 0.783 | 1.552 |
| G12D | KRASi | ESTROGEN RESPONSELATE | 342 | 7.495E-02 | 4.081E-01 | 0.389 | 0.222 | 0.572 | 1.327 |
| G12D | KRASi | IL2 STAT5 SIGNALING | 308 | 1.055E-01 | 4.625E-01 | 0.335 | 0.185 | 0.510 | 1.179 |
| G12D | KRASi | ALLOGRAFT REJECTION | 313 | 1.068E-01 | 4.625E-01 | 0.335 | 0.181 | 0.506 | 1.175 |
| G12D | KRASi | INTERFERON ALPHARESPONSE | 178 | 1.438E-01 | 4.697E-01 | 0.328 | 0.172 | -0.519 | -1.163 |
| G12D | KRASi | INTERFERON GAMMARESPONSE | 311 | 1.883E-01 | 5.768E-01 | 0.239 | 0.151 | -0.482 | -1.115 |
| G12D | KRASi | TNFA SIGNALING VIANFKB | 274 | 2.103E-01 | 6.036E-01 | 0.219 | 0.142 | -0.476 | -1.085 |
| G12D | KRASi | UV RESPONSE DN | 307 | 2.600E-01 | 6.036E-01 | 0.219 | 0.126 | -0.450 | -1.041 |
| G12D | KRASi | EPITHELIALMESENCHYMALTRANSITION | 330 | 3.065E-01 | 6.036E-01 | 0.219 | 0.115 | -0.438 | -1.017 |
| G12D | KRASi | COMPLEMENT | 359 | 3.171E-01 | 6.036E-01 | 0.219 | 0.112 | -0.433 | -1.009 |
| G12D | KRASi | ADIPOGENESIS | 293 | 3.978E-01 | 6.232E-01 | 0.205 | 0.087 | 0.433 | 0.996 |
| G12D | KRASi | APICAL JUNCTION | 373 | 3.978E-01 | 6.232E-01 | 0.205 | 0.087 | 0.423 | 0.983 |
| G12D | KRASi | COAGULATION | 235 | 4.196E-01 | 6.232E-01 | 0.205 | 0.093 | -0.427 | -0.970 |
| G12D | KRASi | XENOBIOTICMETABOLISM | 352 | 4.924E-01 | 6.521E-01 | 0.186 | 0.084 | -0.404 | -0.945 |
| G12D | KRASi | INFLAMMATORYRESPONSE | 314 | 5.636E-01 | 7.268E-01 | 0.139 | 0.080 | -0.399 | -0.923 |
| G12D | KRASi | IL6 JAK STAT3SIGNALING | 121 | 5.974E-01 | 7.476E-01 | 0.126 | 0.066 | 0.429 | 0.913 |
| G12D | KRASi | TGF BETA SIGNALING | 100 | 6.103E-01 | 7.476E-01 | 0.126 | 0.072 | -0.433 | -0.907 |
| G12D | KRASi | BILE ACID METABOLISM | 171 | 6.472E-01 | 7.735E-01 | 0.112 | 0.070 | -0.404 | -0.902 |
| G12D | KRASi | ESTROGEN RESPONSEEARLY | 327 | 7.401E-01 | 7.884E-01 | 0.103 | 0.054 | 0.382 | 0.889 |
| G12D | KRASi | MYOGENESIS | 363 | 7.174E-01 | 7.884E-01 | 0.103 | 0.066 | -0.380 | -0.886 |
| G12D | KRASi_SHP2i | INTERFERON ALPHARESPONSE | 178 | 5.280E-09 | 1.294E-07 | 6.888 | 0.761 | 0.909 | 1.821 |
| G12D | KRASi_SHP2i | INTERFERON GAMMARESPONSE | 311 | 3.027E-10 | 1.483E-08 | 7.829 | 0.814 | 0.870 | 1.811 |
| G12D | KRASi_SHP2i | E2F TARGETS | 266 | 6.088E-07 | 9.943E-06 | 5.002 | 0.659 | 0.846 | 1.750 |
| G12D | KRASi_SHP2i | G2M CHECKPOINT | 298 | 5.042E-04 | 6.176E-03 | 2.209 | 0.477 | 0.772 | 1.604 |
| G12D | KRASi_SHP2i | ALLOGRAFT REJECTION | 313 | 1.040E-03 | 1.019E-02 | 1.992 | 0.455 | 0.753 | 1.566 |
| G12D | KRASi_SHP2i | IL6 JAK STAT3SIGNALING | 121 | 3.381E-02 | 2.367E-01 | 0.626 | 0.282 | 0.726 | 1.422 |
| G12D | KRASi_SHP2i | IL2 STAT5 SIGNALING | 308 | 3.191E-02 | 2.367E-01 | 0.626 | 0.277 | 0.655 | 1.359 |
| G12D | KRASi_SHP2i | HYPOXIA | 317 | 9.226E-02 | 4.740E-01 | 0.324 | 0.288 | -0.604 | -1.349 |
| G12D | KRASi_SHP2i | INFLAMMATORYRESPONSE | 314 | 5.862E-02 | 3.590E-01 | 0.445 | 0.200 | 0.615 | 1.281 |
| G12D | KRASi_SHP2i | COAGULATION | 235 | 1.247E-01 | 5.092E-01 | 0.293 | 0.135 | 0.588 | 1.201 |
| G12D | KRASi_SHP2i | MYC TARGETS V2 | 68 | 2.221E-01 | 6.175E-01 | 0.209 | 0.105 | 0.613 | 1.147 |
| G12D | KRASi_SHP2i | TNFA SIGNALING VIANFKB | 274 | 1.839E-01 | 6.034E-01 | 0.219 | 0.107 | 0.554 | 1.145 |
| G12D | KRASi_SHP2i | COMPLEMENT | 359 | 2.269E-01 | 6.175E-01 | 0.209 | 0.092 | 0.527 | 1.107 |
| G12D | KRASi_SHP2i | EPITHELIALMESENCHYMALTRANSITION | 330 | 2.847E-01 | 6.298E-01 | 0.201 | 0.080 | 0.511 | 1.068 |
| G12D | KRASi_SHP2i | XENOBIOTICMETABOLISM | 352 | 3.039E-01 | 6.298E-01 | 0.201 | 0.076 | 0.506 | 1.062 |
| G12D | KRASi_SHP2i | ESTROGEN RESPONSEEARLY | 327 | 3.714E-01 | 6.738E-01 | 0.172 | 0.067 | 0.495 | 1.033 |
| G12D | KRASi_SHP2i | ESTROGEN RESPONSELATE | 342 | 3.725E-01 | 6.738E-01 | 0.172 | 0.066 | 0.494 | 1.031 |
| G12D | KRASi_SHP2i | UV RESPONSE DN | 307 | 4.551E-01 | 6.738E-01 | 0.172 | 0.058 | 0.478 | 0.992 |
| G12D | KRASi_SHP2i | MYC TARGETS V1 | 254 | 4.921E-01 | 6.890E-01 | 0.162 | 0.055 | 0.479 | 0.985 |
| G12D | KRASi_SHP2i | ADIPOGENESIS | 293 | 5.401E-01 | 7.351E-01 | 0.134 | 0.050 | 0.466 | 0.966 |
| G12D | KRASi_SHP2i | APICAL JUNCTION | 373 | 7.867E-01 | 8.566E-01 | 0.067 | 0.031 | 0.412 | 0.866 |
| G12D | KRASi_SHP2i | BILE ACID METABOLISM | 171 | 7.675E-01 | 8.566E-01 | 0.067 | 0.036 | 0.424 | 0.850 |
| G12D | KRASi_SHP2i | MYOGENESIS | 363 | 8.916E-01 | 9.295E-01 | 0.032 | 0.024 | 0.385 | 0.807 |
| G12D | KRASi_SHP2i | TGF BETA SIGNALING | 100 | 8.583E-01 | 9.143E-01 | 0.039 | 0.035 | 0.413 | 0.791 |

### Table S3: GSEA significant targets across all treatment groups

| Treatment | pathway | Exp genes | pval | padj | $log_{10}$(padj) | log2err | ES | NES |
|---|---|---|---|---|---|---|---|---|
| Ki_Si_vs_Ki | INTERFERON ALPHARESPONSE | 178 | 8.313E-16 | 4.073E-14 | 13.390 | 1.028 | 0.904 | 2.025 |
| Ki_Si_vs_Ki | MYC TARGETS V1 | 254 | 8.810E-12 | 1.439E-10 | 9.842 | 0.875 | 0.827 | 1.947 |
| Ki_Si_vs_Ki | INTERFERON GAMMARESPONSE | 311 | 8.674E-12 | 1.439E-10 | 9.842 | 0.875 | 0.804 | 1.940 |
| Ki_Si_vs_Ki | E2F TARGETS | 266 | 1.648E-09 | 2.019E-08 | 7.695 | 0.788 | 0.792 | 1.886 |
| Ki_Si_vs_Ki | MYC TARGETS V2 | 68 | 1.452E-05 | 1.423E-04 | 3.847 | 0.593 | 0.873 | 1.718 |
| Ki_Si_vs_Ki | OXIDATIVEPHOSPHORYLATION | 211 | 4.419E-04 | 3.609E-03 | 2.443 | 0.498 | 0.692 | 1.592 |
| Ki_Si_vs_Ki | G2M CHECKPOINT | 298 | 1.804E-03 | 9.824E-03 | 2.008 | 0.455 | 0.622 | 1.486 |
| Ki_Si_vs_Ki | BILE ACID METABOLISM | 171 | 1.321E-03 | 8.369E-03 | 2.077 | 0.455 | -0.717 | -1.479 |
| Ki_Si_vs_Ki | UNFOLDED PROTEINRESPONSE | 169 | 7.890E-03 | 3.515E-02 | 1.454 | 0.381 | 0.655 | 1.455 |
| Ki_Si_vs_Ki | KRAS SIGNALING UP | 355 | 1.366E-03 | 8.369E-03 | 2.077 | 0.455 | -0.641 | -1.426 |
| Ki_Si_vs_Ki | CHOLESTEROLHOMEOSTASIS | 124 | 1.664E-02 | 6.793E-02 | 1.168 | 0.352 | -0.687 | -1.375 |
| Ki_Si_vs_Ki | APICAL JUNCTION | 373 | 7.067E-03 | 3.463E-02 | 1.461 | 0.407 | -0.611 | -1.361 |
| Ki_Si_vs_Ki | TGF BETA SIGNALING | 100 | 7.029E-02 | 2.460E-01 | 0.609 | 0.214 | -0.674 | -1.317 |
| Ki_Si_vs_Ki | DNA REPAIR | 184 | 1.071E-01 | 3.088E-01 | 0.510 | 0.238 | 0.534 | 1.210 |
| Ki_Si_vs_Ki | MITOTIC SPINDLE | 361 | 8.243E-02 | 2.693E-01 | 0.570 | 0.180 | -0.539 | -1.202 |
| Ki_Si_vs_Ki | COAGULATION | 235 | 1.223E-01 | 3.153E-01 | 0.501 | 0.151 | -0.561 | -1.201 |
| Ki_Si_vs_Ki | ALLOGRAFT REJECTION | 313 | 9.574E-02 | 2.932E-01 | 0.533 | 0.277 | 0.496 | 1.197 |
| Ki_Si_vs_Ki | COMPLEMENT | 359 | 1.949E-01 | 4.151E-01 | 0.382 | 0.111 | -0.507 | -1.129 |
| Ki_Si_vs_Ki | ANGIOGENESIS | 57 | 2.997E-01 | 4.895E-01 | 0.310 | 0.097 | -0.618 | -1.120 |
| Ki_Si_vs_Ki | INFLAMMATORYRESPONSE | 314 | 2.657E-01 | 4.650E-01 | 0.333 | 0.094 | -0.496 | -1.092 |
| Ki_Si_vs_Ki | TNFA SIGNALING VIANFKB | 274 | 6.705E-01 | 7.864E-01 | 0.104 | 0.048 | -0.420 | -0.914 |
| Ki_PD1_vs_Ki | TGF BETA SIGNALING | 100 | 4.092E-03 | 3.341E-02 | 1.476 | 0.407 | -0.832 | -1.773 |
| Ki_PD1_vs_Ki | INTERFERON GAMMARESPONSE | 311 | 3.270E-04 | 1.602E-02 | 1.795 | 0.498 | 0.935 | 1.758 |
| Ki_PD1_vs_Ki | INTERFERON ALPHARESPONSE | 178 | 1.626E-03 | 2.034E-02 | 1.692 | 0.455 | 0.935 | 1.731 |
| Ki_PD1_vs_Ki | ALLOGRAFT REJECTION | 313 | 1.273E-03 | 2.034E-02 | 1.692 | 0.455 | 0.908 | 1.709 |
| Ki_PD1_vs_Ki | E2F TARGETS | 266 | 2.270E-03 | 2.225E-02 | 1.653 | 0.432 | -0.747 | -1.646 |
| Ki_PD1_vs_Ki | MITOTIC SPINDLE | 361 | 1.660E-03 | 2.034E-02 | 1.692 | 0.455 | -0.727 | -1.588 |
| Ki_PD1_vs_Ki | G2M CHECKPOINT | 298 | 2.124E-02 | 1.387E-01 | 0.858 | 0.352 | -0.653 | -1.436 |
| Ki_PD1_vs_Ki | APICAL JUNCTION | 373 | 8.214E-02 | 3.354E-01 | 0.474 | 0.168 | 0.757 | 1.422 |
| Ki_PD1_vs_Ki | DNA REPAIR | 184 | 5.227E-02 | 2.561E-01 | 0.592 | 0.322 | -0.631 | -1.384 |
| Ki_PD1_vs_Ki | TNFA SIGNALING VIANFKB | 274 | 1.125E-01 | 4.240E-01 | 0.373 | 0.146 | 0.733 | 1.374 |
| Ki_PD1_vs_Ki | INFLAMMATORYRESPONSE | 314 | 1.746E-01 | 4.417E-01 | 0.355 | 0.112 | 0.674 | 1.269 |
| Ki_PD1_vs_Ki | COMPLEMENT | 359 | 1.748E-01 | 4.417E-01 | 0.355 | 0.110 | 0.667 | 1.254 |
| Ki_PD1_vs_Ki | MYC TARGETS V1 | 254 | 1.893E-01 | 4.417E-01 | 0.355 | 0.228 | -0.542 | -1.199 |
| Ki_PD1_vs_Ki | OXIDATIVEPHOSPHORYLATION | 211 | 2.146E-01 | 4.688E-01 | 0.329 | 0.200 | -0.527 | -1.159 |
| Ki_PD1_vs_Ki | ANGIOGENESIS | 57 | 2.560E-01 | 4.825E-01 | 0.317 | 0.105 | 0.662 | 1.158 |
| Ki_PD1_vs_Ki | COAGULATION | 235 | 2.772E-01 | 5.031E-01 | 0.298 | 0.086 | 0.606 | 1.127 |
| Ki_PD1_vs_Ki | BILE ACID METABOLISM | 171 | 3.158E-01 | 5.336E-01 | 0.273 | 0.157 | -0.478 | -1.050 |
| Ki_PD1_vs_Ki | KRAS SIGNALING UP | 355 | 3.698E-01 | 5.491E-01 | 0.260 | 0.068 | 0.549 | 1.033 |
| Ki_PD1_vs_Ki | UNFOLDED PROTEINRESPONSE | 169 | 7.338E-01 | 8.362E-01 | 0.078 | 0.041 | 0.419 | 0.777 |
| Ki_PD1_vs_Ki | MYC TARGETS V2 | 68 | 9.091E-01 | 9.366E-01 | 0.028 | 0.067 | -0.375 | -0.766 |
| Ki_PD1_vs_Ki | CHOLESTEROLHOMEOSTASIS | 124 | 9.175E-01 | 9.366E-01 | 0.028 | 0.034 | 0.375 | 0.685 |
| Ki_Si_vs_Ki_PD1 | E2F TARGETS | 266 | 2.676E-20 | 1.311E-18 | 17.882 | 1.169 | 0.778 | 2.250 |
| Ki_Si_vs_Ki_PD1 | MYC TARGETS V1 | 254 | 9.614E-17 | 2.355E-15 | 14.628 | 1.057 | 0.746 | 2.141 |
| Ki_Si_vs_Ki_PD1 | INTERFERON ALPHARESPONSE | 178 | 1.228E-14 | 2.006E-13 | 12.698 | 0.987 | 0.781 | 2.119 |
| Ki_Si_vs_Ki_PD1 | G2M CHECKPOINT | 298 | 7.725E-09 | 7.570E-08 | 7.121 | 0.748 | 0.628 | 1.826 |
| Ki_Si_vs_Ki_PD1 | OXIDATIVEPHOSPHORYLATION | 211 | 2.138E-07 | 1.497E-06 | 5.825 | 0.690 | 0.642 | 1.800 |
| Ki_Si_vs_Ki_PD1 | INTERFERON GAMMARESPONSE | 311 | 5.942E-09 | 7.279E-08 | 7.138 | 0.761 | 0.612 | 1.790 |
| Ki_Si_vs_Ki_PD1 | MYC TARGETS V2 | 68 | 1.921E-04 | 1.046E-03 | 2.981 | 0.519 | 0.731 | 1.731 |
| Ki_Si_vs_Ki_PD1 | KRAS SIGNALING UP | 355 | 9.295E-09 | 7.591E-08 | 7.120 | 0.748 | -0.688 | -1.656 |
| Ki_Si_vs_Ki_PD1 | UNFOLDED PROTEINRESPONSE | 169 | 1.385E-03 | 5.222E-03 | 2.282 | 0.455 | 0.551 | 1.486 |
| Ki_Si_vs_Ki_PD1 | ANGIOGENESIS | 57 | 1.378E-02 | 3.973E-02 | 1.401 | 0.381 | -0.718 | -1.470 |
| Ki_Si_vs_Ki_PD1 | COAGULATION | 235 | 9.022E-04 | 3.684E-03 | 2.434 | 0.477 | -0.618 | -1.457 |
| Ki_Si_vs_Ki_PD1 | INFLAMMATORYRESPONSE | 314 | 3.860E-04 | 1.720E-03 | 2.765 | 0.498 | -0.607 | -1.450 |
| Ki_Si_vs_Ki_PD1 | COMPLEMENT | 359 | 1.893E-04 | 1.046E-03 | 2.981 | 0.519 | -0.595 | -1.434 |
| Ki_Si_vs_Ki_PD1 | BILE ACID METABOLISM | 171 | 3.152E-03 | 1.103E-02 | 1.957 | 0.432 | -0.624 | -1.431 |
| Ki_Si_vs_Ki_PD1 | APICAL JUNCTION | 373 | 2.279E-04 | 1.117E-03 | 2.952 | 0.519 | -0.591 | -1.426 |
| Ki_Si_vs_Ki_PD1 | DNA REPAIR | 184 | 7.998E-03 | 2.613E-02 | 1.583 | 0.381 | 0.495 | 1.338 |
| Ki_Si_vs_Ki_PD1 | CHOLESTEROLHOMEOSTASIS | 124 | 1.898E-02 | 4.895E-02 | 1.310 | 0.352 | -0.599 | -1.333 |
| Ki_Si_vs_Ki_PD1 | TGF BETA SIGNALING | 100 | 5.563E-02 | 1.157E-01 | 0.937 | 0.217 | -0.604 | -1.314 |
| Ki_Si_vs_Ki_PD1 | TNFA SIGNALING VIANFKB | 274 | 1.255E-02 | 3.844E-02 | 1.415 | 0.381 | -0.553 | -1.312 |
| Ki_Si_vs_Ki_PD1 | ALLOGRAFT REJECTION | 313 | 1.612E-02 | 4.388E-02 | 1.358 | 0.352 | -0.539 | -1.289 |
| Ki_Si_vs_Ki_PD1 | MITOTIC SPINDLE | 361 | 4.249E-01 | 5.783E-01 | 0.238 | 0.057 | -0.424 | -1.021 |

### Table S4: Real-time PCR primers

| Primers | Forward | Reverse |
|---|---|---|
| Dusp5 | AGTGTGGAAAGCCCGTTCTC | GAAGGATTTCAACCGGGCCA |
| Etv4 | CCATGGAGAGCAGTGCCTTT | CTGCTGTTCTGCCCTGGAA |
| Etv5 | TGATGATGAGCAGTTTGTCCCA | TGGCTACAGGACGACAACTC |
| Myc | CCGGGGAGGGAATTTTTGTCT | GAGGGGCATCGTCGTGG |
| Stat1 | AAGTCTGGCAGCTGAGTTCC | TCTTCGGTGACAATGAGAGGC |
| Cxcl9 | CCAAGCCCCAATTGCAACAAA | GTCCGGATCTAGGCAGGTTT |
| Irf9 | GCCGAGTGGTGGGTAAGAC | GCAAAGGCGCTGAACAAAGAG |
| B2m | TCTCACTGACCGGCCTGTAT | ATTTCAATGTGAGGCGGGTG |
| Hsp90 | AGATTCCACTAACCGACGCC | TGCTCTTTGCTCTCACCAGT |
| Sdha | TCGACAGGGGAATGGTTTGG | TCATACTCATCGACCCGCAC |

### Table S5: Spectral flow staining panel

| Target | Fluorophore | Ab clone | Cat. No. | Company | Ab vol [ul]l / 100 ul |
|---|---|---|---|---|---|
| B220 | BUV496 | RA3-6B2 | 612950 | BD Biosciences | 0.25 |
| CD11c | BUV563 | N418 | 749040 | BD Biosciences | 0.75 |
| TIM3 | BUV661 | RMT2-23 | 753149 | BD Biosciences | 1 |
| CD62L | BUV737 | MEL-14 | 612833 | BD Biosciences | 0.5 |
| CD3 | BUV805 | 17A2 | 569192 | BD Biosciences | 0.75 |
| CD44 | BV421 | IM7 | 103039 | BioLegend | 0.5 |
| CD11b | BV480 | M1/70 | 566117 | BD Biosciences | 0.25 |
| CD69 | BV510 | H1.2F3 | 104531 | BioLegend | 0.75 |
| CD19 | BV570 | 6D5 | 115535 | BioLegend | 0.5 |
| CD103 | BV605 | 2D7 | 121433 | BioLegend | 1 |
| PD-1 | BV650 | J43 | 569506 | BD Biosciences | 0.5 |
| Ly6G | BV711 | 1A8 | 127643 | BioLegend | 0.5 |
| Ly6C | BV785 | HK1.4 | 128041 | BioLegend | 0.5 |
| CD8α | Spark Blue 550 | 53-6.7 | 100779 | BioLegend | 0.5 |
| H-2Kb | NFB610/70S | AF6-88.5.5.3 | 17836316 | Invitrogen | 0.5 |
| CD45 | PerCP | 30-F11 | 103129 | BioLegend | 0.5 |
| MHCII | PerCP-eFluor710 | M5/114.15.2 | 46-5321-82 | eBioscience | 0.15 |
| CD71 | RB780 | C2 | 755614 | BD Biosciences | 0.25 |
| LAG3 | PerCP/Fire780 | C9BV7 | 125251 | BioLegend | 0.75 |
| PD-L1 | PE | 10F.9G2 | 124307 | BioLegend | 0.25 |
| NK1.1 | PE/Cy5 | PK136 | 108715 | BioLegend | 0.5 |
| CD64 | PE/Cy7 | X54-5/7.1 | 139313 | BioLegend | 0.5 |
| Arginase 1 | eFluor450 | A1exF5 | 48-3697-82 | eBioscience | 0.75 |
| FOXP3 | eFluor660 | FJK-16 | 50-5773-82 | eBioscience | 0.5 |
| CD86 | AF-700 | GL-1 | 105023 | BioLegend | 0.25 |
| CD4 | APC/Fire 810 | GK1.5 | 100479 | BioLegend | 0.5 |
| Viab. Dye | Zombie NIR | - | 423105 | Biolegend | 0.05 |

### Table S6: Spectral flow staining panel for analysis of Emv2-specific T cells

| Antigen | Fluorophore | Clone | Cat. No | Company | Ab vol/ 100 ul | Staining |
|---|---|---|---|---|---|---|
| EMV2-peptide | PE | - | MBL-TS-M507-1 | MBL | 1 | H-2Kb MuLV p15E Tetramer-KSPWFTTL |
| CD107a | BUV395 | 1D4B | 565533 | BD Biosciences | 1 | Ab - Extracellular |
| Ki67 | BUV615 | SolA15 | 366-5698-82 | eBioscience | 0.06 | Ab -Intracellular |
| CD62L | BUV737 | MEL-14 | 612833 | BD Biosciences | 0.5 | Ab - Extracellular |
| CD3 | BUV805 | 17A2 | 569192 | BD Biosciences | 1 | Ab - Extracellular |
| CD44 | BV421 | IM7 | 103039 | BioLegend | 0.5 | Ab - Extracellular |
| CD11b | BV480 | M1/70 | 566117 | BD Biosciences | 0.25 | Ab - Extracellular |
| CD69 | BV510 | H1.2F3 | 104531 | BioLegend | 1 | Ab - Extracellular |
| NK1.1 | BV605 | PK136 | 108739 | BioLegend | 1 | Ab - Extracellular |
| PD1 | BV650 | J43 | 569506 | BD Biosciences | 0.5 | Ab - Extracellular |
| TIM3 | BV785 | RMT3-23 | 119725 | BioLegend | 1 | Ab - Extracellular |
| CD8a | FITC | KT15 | MA5-16759 | Thermofisher | 0.25 | Ab - Extracellular |
| CD45 | PerCP | 30-F11 | 103129 | BioLegend | 0.5 | Ab - Extracellular |
| LAG3 | PerCP-F780 | C9BV7 | 125251 | BioLegend | 1 | Ab - Extracellular |
| GzmB | APC | 372204 | QA16A02 | Biolegend | 0.25 | Ab -Intracellular |
| Foxp3 | eFluor 660 | FJK-16 | 50-5773-82 | eBioscience | 0.5 | Ab -Intracellular |
| CD4 | APC-F810 | GK1.5 | 100479 | BioLegend | 0.5 | Ab - Extracellular |
| Live/Dead | Zombie NIR | - | 423105 | Biolegend | 0.05 | Live/Dead dye |
